# Human cell cytoskeletal rigidity differentially influences trogocytosis and phagocytosis by *Entamoeba histolytica*

**DOI:** 10.64898/2026.03.17.712427

**Authors:** Felina P. Loya, Tianle Wang, Mary C. Irani, Rene L. Suleiman, Gang-Yu Liu, Katherine S. Ralston

**Author notes:** Correspondence (K.S. Ralston).

## Abstract

*Entamoeba histolytica* is a parasitic amoeba and the cause of amoebiasis, a common but understudied human diarrheal disease. *E. histolytica* trophozoites (“amoebae”) kill human cells through a process of cell-nibbling called trogocytosis (*trogo*-: nibble) that contributes to tissue damage. Amoebae can also perform phagocytosis, in which entire human cells are ingested. Based on studies in which human cells were artificially stiffened, it was suggested that amoebae perform phagocytosis on stiffer cells, and trogocytosis on less stiff cells. A few recent studies of macrophage ingestion of artificial targets or artificially stiffened target cells also suggested a similar relationship between target stiffness and trogocytosis/phagocytosis efficiencies. To better evaluate the impact of target cell stiffness on amoebic ingestion, instead of using artificial targets or artificial cell stiffening, we created human cell mutants in which individual genes were knocked down. We targeted seven different genes in the Rho-pathway, which have well-characterized roles in regulating actomyosin dynamics. Strikingly, amoebae performed quantitatively reduced levels of trogocytosis on all knockdown mutants, including relatively stiffer mutants and relatively less stiff mutants. In contrast, amoebic phagocytosis efficiency was inversely correlated with human cell stiffness. Thus, human cell stiffness differentially influences amoebic trogocytosis and phagocytosis. This is more complex than the conclusions of studies that used artificial targets or artificially stiffened cells. Our results emphasize that the dynamic nature of the cytoskeleton in living cells impacts trogocytosis. In addition to shedding light on the burgeoning field of eukaryotic trogocytosis, this work extends knowledge of amoebic ingestion processes that contribute to disease.

**Author Summary:** *Entamoeba histolytica* is an amoeba and the causative agent of amoebiasis, a human diarrheal disease that is particularly common in developing countries. This amoeba can kill human cells by nibbling on them, which is called trogocytosis, and it can also fully ingest human cells, which is called phagocytosis. It was not clear what caused amoebae to perform trogocytosis versus phagocytosis, but had been suggested that amoebae preferred to perform phagocytosis when human cells were artificially stiffened. Here, instead of artificially stiffening human cells, we generated mutant human cells with abnormal stiffness by targeting genes impacting the actomyosin cytoskeleton. Amoebae performed reduced levels of trogocytosis on all human cell mutants regardless of whether they were relatively stiffer or less stiff. Unlike trogocytosis, amoebic phagocytosis efficiency was inversely correlated with human cell stiffness. These findings are more complex than the conclusions of studies that used artificially stiffened cells, emphasizing that the cytoskeleton is dynamic. These findings tie into the burgeoning field of eukaryotic trogocytosis, since macrophage trogocytosis/phagocytosis has also been studied using artificial targets. As amoebic trogocytosis and phagocytosis are relevant to the pathogenesis of amoebiasis, this work also extends knowledge of amoebic processes that contribute to disease.

## Introduction

Amoebiasis is a human diarrheal disease caused by the parasitic amoeba *Entamoeba histolytica* (1). Fecal-oral transmission of *E. histolytica* is common in countries with insufficient water sanitation infrastructure (1,2). There is no vaccine and treatment options are limited (3,4). Although most of the ∼50 million infections/year are asymptomatic, *E. histolytica* trophozoites (“amoebae”) can invade and damage the colonic epithelium, causing bloody diarrhea (5). Amoebae can also spread to other organs, most frequently the liver, which results in abscesses that are fatal if left untreated (2,6). Amoebiasis results in an estimated 15,500 deaths/year in children and 67,900 deaths/year in adults (7–10). In addition to mortality, amoebiasis is a major source of morbidity. Amoebiasis is common in children in endemic areas, and is associated with malnutrition and stunted growth (3,4,11,12). Despite the global burden of amoebiasis, there is a limited understanding of the molecular pathogenesis of disease.

The profound cell killing activity (13) of amoebae likely underlies the prolific tissue damage in invasive infections. The previously accepted model was that amoebae secrete the pore-forming protein amoebapore A, and that it acts on host cell membranes (14–18). However, there is no experimental evidence that amoebapore A is secreted. Additionally, amoebapore A requires pH 5.2 for pore-forming activity (19), which is not consistent with physiological pH extracellularly (20). The lack of cell killing activity in amoebic culture supernatants, lysates, and killed amoebae (14,15,17,18,21,22), further refutes a role for secreted cell killing effectors. Establishing a new paradigm, we showed that amoebae kill human cells through a process of cell-nibbling called trogocytosis (*trogo*-: nibble) (13). During trogocytosis, amoebae pinch off and internalize bites of human cell membrane and intracellular contents, eventually resulting in death of the bitten cell (13). Inhibiting trogocytosis also inhibited amoebic invasion in a tissue explant model, suggesting relevance to the pathogenesis of invasive amoebiasis (13).

In addition to its role in *E. histolytica*, trogocytosis is a widespread and possibly fundamental form of cell-to-cell interaction in eukaryotic biology (23–28). Trogocytosis was first described as a mechanism for cell killing used by various amoebae across at least two eukaryotic supergroups (29,28,30,31). Trogocytosis was later uncovered as a mode of cell-cell interaction occurring between mammalian immune cells, that did not result in killing of the nibbled cell (27). More recent studies clearly show that trogocytosis performed by mammalian immune cells can also result in cell killing (32,33). Additionally, trogocytosis has also been shown to function in cell-cell remodeling in a variety of scenarios, where one cell nibbles and removes components of another cell (34–38). Trogocytosis can thus function in cell killing, cell-cell communication, and cell-cell remodeling, and it has roles during embryonic development, normal physiology, and disease. With the broad occurrence of trogocytosis across single and multicellular organisms in a variety of eukaryotic supergroups, it is possible that trogocytosis is a conserved process. However, the underlying molecular mechanism has not been well described in any organism, and thus the molecular details remain unclear.

Trogocytosis is clearly related to the process of phagocytosis, in which entire cells are consumed. In general, shared mechanisms exist between both processes (13,32,39–43), though aspects of the trogocytosis mechanism that are distinct from the phagocytosis mechanism are emerging (23,40,42–47). *E. histolytica* amoebae can perform both trogocytosis and phagocytosis of human cells, and both forms of ingestion require Gal/GalNAc lectin signaling, phosphatidylinositol 3-kinase (PI3K) activity, and C2 domain-containing kinase (EhC2PK) activation (48–50). In addition to the relevance of trogocytosis to tissue invasion in amoebiasis, phagocytosis is also relevant to pathogenesis (51–54). Whether amoebae perform trogocytosis or phagocytosis seems to be influenced by the state of the target cell. When amoebae ingested Jurkat T cells, they performed trogocytosis on living cells, but instead performed phagocytosis on cells that were artificially killed (13). Amoebae performed decreasing levels of trogocytosis, and increasing levels of phagocytosis, on red blood cells artificially stiffened by treatment with elevating glutaraldehyde concentrations (55). Thus, the stiffness of the human cell may influence the type of ingestion amoebae perform.

The stiffness of the human cell is driven by a dynamic, interconnected network of actin filaments, microtubules, and intermediate filaments, all of which modulate cell motility and shape, intracellular organization, and mechanical support during critical cellular processes (56–59). Cortical tension and stiffness are directly impacted by F-actin networking and architecture (60–62), which is primarily modulated by Rho-GTPase actin regulators (63) (Figure 1). Typically, in response to mechanical stimuli, Rho-GTPases Ras homolog family member A (RhoA) and Rac family small GTPase 1 or 2 (Rac1/2) are activated and induce signaling cascades, which promote the formation of actin stress fibers and focal adhesions that temporarily deform the cell membrane (63–66). Activation of Rac1/2 mediates downstream actin polymerization through actin-binding proteins like Cofilin-1 (CFL1), regulating actin fiber stability (67–72). RhoA activates Rho-associated protein kinases 1 or 2 (ROCK1/2), which phosphorylate the myosin-binding subunit 1 of myosin phosphatase (MYPT1) to modulate actomyosin contractility and mechanosensitive responses to the extracellular environment (73–79).

**Figure 1.**
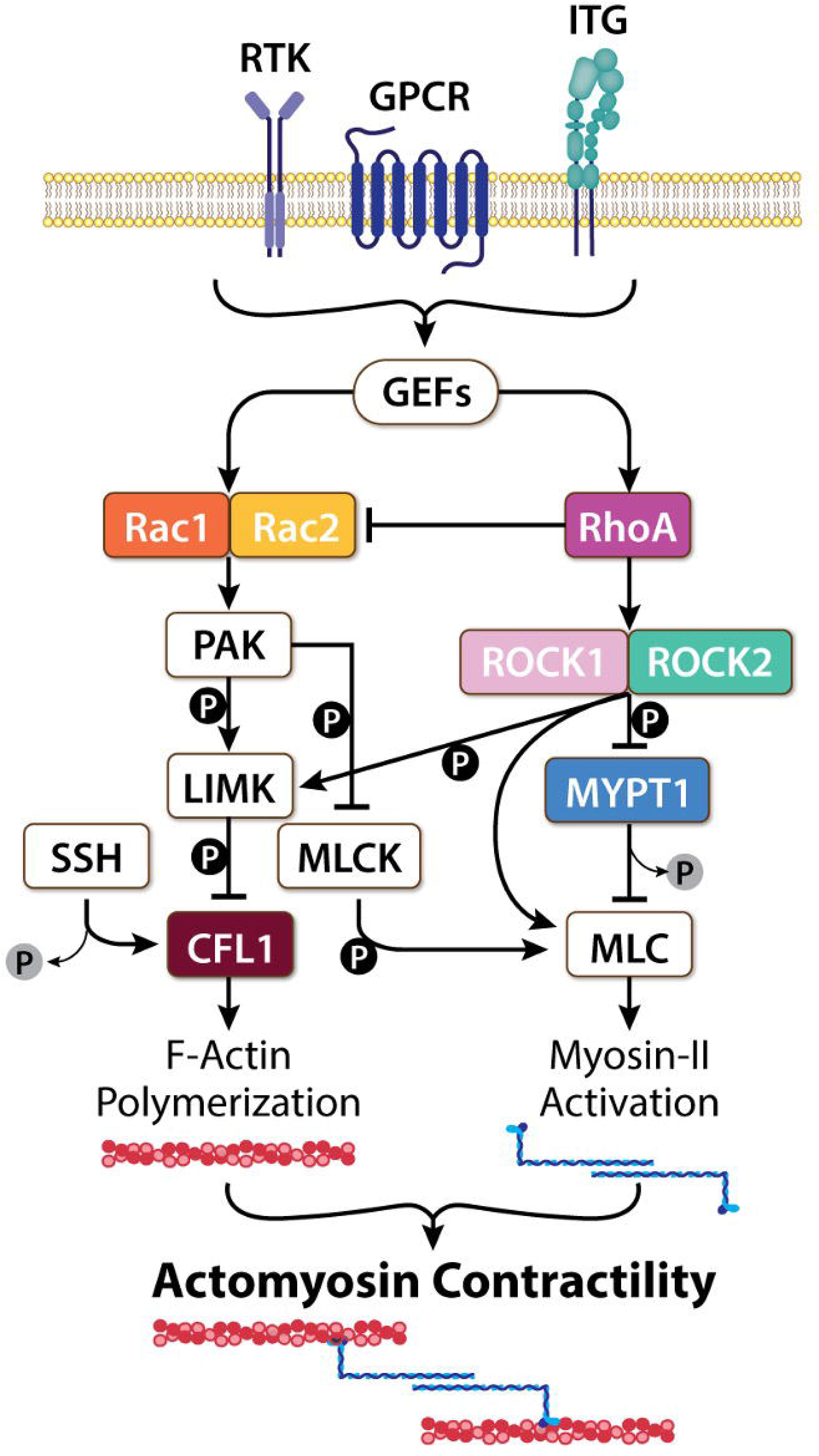
Primary Rho-pathway of actomyosin regulation. Ras homologous (Rho) guanine nucleotide exchange factors (GEFs) are activated following a cascade of intracellular signaling imposed by binding transmembrane receptors, such as receptor tyrosine kinases (RTKs), G-protein coupled receptors (GPCRs), or integrins (INTs), at the extracellular matrix (ECM). RhoGEFs promote activation of Rho-GTPases, which regulate multiple signal transduction pathways that regulate actin reorganization. The RhoA pathway is required for Myosin II activation, whereas the Rac pathway (Rac1/2) is required for actin polymerization. Crosstalk between these pathways allows for actomyosin stability and contractility throughout the cytoskeleton. Image credit: Anita Impagliazzo.

Beyond amoebae, there are several characterized scenarios where the stiffness of the target cell influenced its ingestion. For example, macrophages phagocytosed artificially stiffened red blood cells more rapidly than native red blood cells, and this was associated with greater activation of myosin-II (80). Similarly, macrophages phagocytosed stiffer polyacrylamide beads (81) or hydrogel microparticles (82) more often than softer beads or microparticles. Detailed studies of macrophage phagocytosis of stiff vs. soft hydrogel microparticles showed that phagocytic cups formed efficiently with stiff particles, which were “gulped” rapidly, but phagocytic cups tended to stall with soft particles (83). Macrophage β-integrin was identified as a key mediator for sensing target stiffness, since macrophages lacking β-integrin lost the ability to efficiently gulp stiff particles (83). Finally, in a different ingestion scenario beyond macrophage phagocytosis, physical stiffness of the target also impacted ingestion. During the process of entosis, in which epithelial cells can engulf one another, rigid epithelial cells were more likely to be ingested (i.e., to be the “losers” of entosis), while deformable epithelial cells were more likely to perform ingestion (i.e., to be the “winners” of entosis) (84).

Recent studies hint that whether macrophages perform trogocytosis vs. phagocytosis might also be influenced by physical properties of the target. When giant unilamellar vesicles (GUVs) were opsonized and used as a target for ingestion, macrophages predominantly performed phagocytosis on GUVs that had higher membrane tension and predominantly performed trogocytosis on GUVs that had lower membrane tension (85). In another recent study, macrophages were found to perform greater levels of phagocytosis on suspension cell lines and greater levels of trogocytosis on adherent cell lines (86). These observations in macrophages begin to suggest that physical properties of the target could influence the type of ingestion that is performed.

To test the potential for the cytoskeletal rigidity of human cells to influence amoebic ingestion, we first pharmacologically inhibited F-actin in human Jurkat T cells and found that amoebic trogocytosis, but not phagocytosis, was inhibited. With further consideration to the dynamic capacity of the cytoskeleton, we created seven different human cell mutants in which individual Rho-pathway genes were knocked down. Strikingly, amoebae performed quantitatively reduced levels of trogocytosis on all knockdown mutants, regardless of the relative stiffness of these mutants. In contrast, amoebic phagocytosis efficiency inversely correlated with the stiffness of human cells. Our results show, for the first time, that the human cell cytoskeleton influences amoebic phagocytosis, but that amoebic trogocytosis regulation is more complex than a simple correlation between cytoskeletal stiffness and trogocytosis efficiency.

## Results

### Pharmacological inhibition of human cell F-actin leads to impaired amoebic trogocytosis

We first used pharmacological inhibition to investigate if the stiffness of the human cell influences amoebic ingestion. Owing to its relatively long-lasting impact on actin polymerization, we opted to treat human Jurkat T cells with mycalolide B (MycB). MycB severs F-actin filaments and covalently sequesters G-actin, resulting in actin depolymerization (87–91). Compared to untreated and vehicle controls, MycB-treated Jurkat cells were rounder in shape, had less overall F-actin, and lacked secondary protrusions like filopodia (Figure 2A).

**Figure 2.**
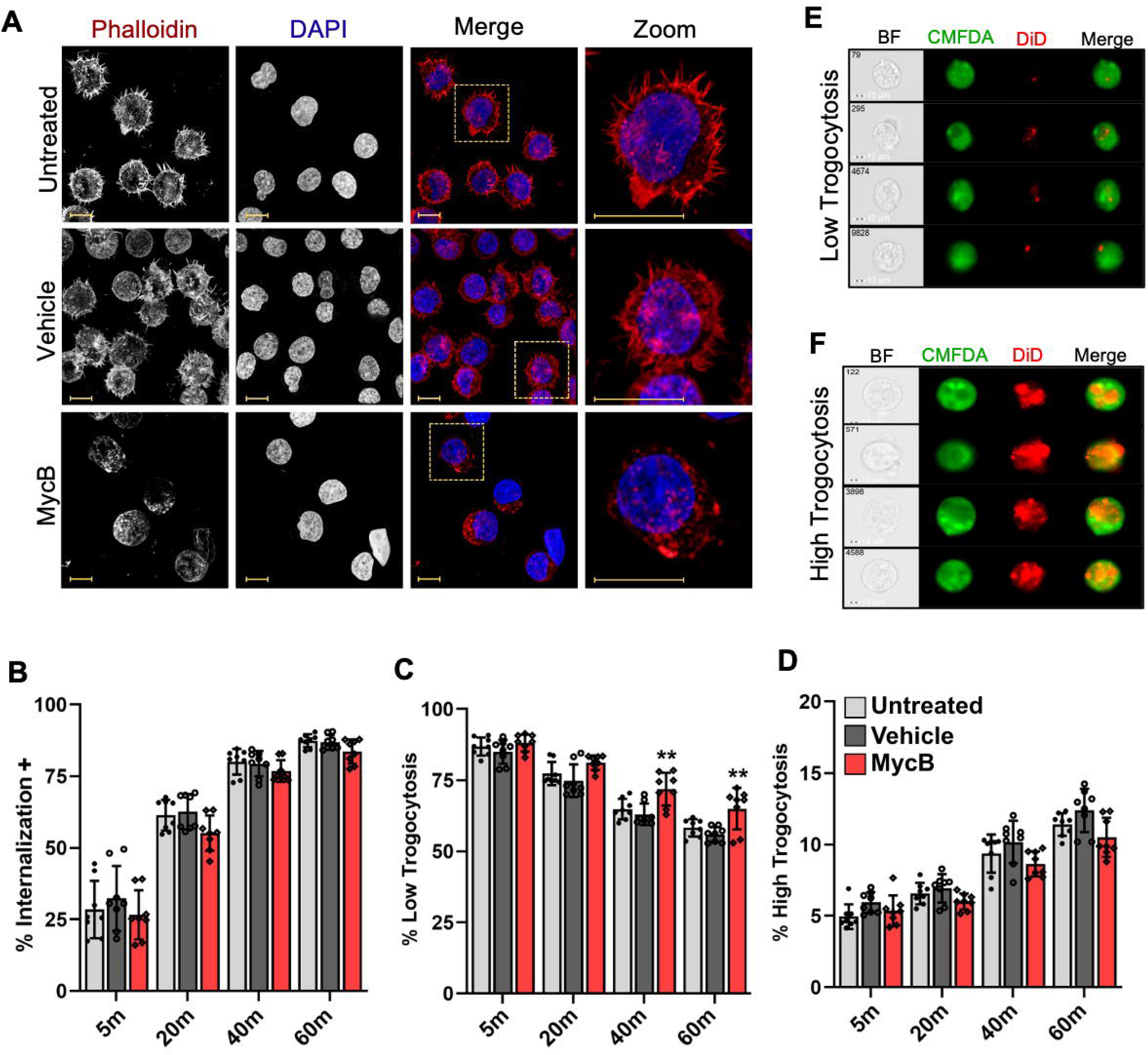
Pharmacological inhibition of human cell F-actin leads to impaired amoebic trogocytosis. A, Untreated, vehicle-treated, or MycB-treated human Jurkat T cells were stained with phalloidin to label F-actin (red) and stained with DAPI to label DNA (blue). Images are representative of >50 cells imaged for each cell line. Scale bar (yellow), 10 μm. **B-F,** Amoebae were stained with CMFDA and Jurkat cells (untreated, vehicle-treated, or MycB-treated) were stained with DiD. Cells were co-incubated for 5, 20, 40 or 60 minutes, and amoebic trogocytosis was quantified using imaging flow cytometry (see Figure S1). **B,** The percentage of amoebae that performed ingestion was quantified (as % Internalization+). **C,** Among amoebae that had performed ingestion, the proportion of amoebae that contained a low level of bites was quantified (% Low Trogocytosis). **D,** Among amoebae that had performed ingestion, the proportion of amoebae that contained a high level of bites was quantified (% High Trogocytosis). Data represent four separate biological replicates with two technical replicates per condition, for a total of eight replicates per condition. Statistical analyses were performed using ordinary 2-way ANOVA with Dunnett’s multiple comparisons test. **E-F,** Representative images of amoebae that performed low or high levels of trogocytosis. Brightfield (BF), CMFDA (green), DiD (red) and merged images are shown. Image numbers represent the order in which they were acquired during imaging flow cytometry. Scale bar, 10 μm.

Amoebae were incubated with untreated, vehicle-treated, or MycB-treated Jurkat cells, and imaging flow cytometry was used to quantify trogocytosis (Figure 2B-F; Supplemental Figure 1). Trogocytosis by amoebae requires attachment to the human cell (92,93) and an initial parameter to assess trogocytosis is the percentage of amoebae that performed ingestion (% Internalization+), (Supplemental Figure 1E). There was no significant difference in the % Internalization+ for amoebae incubated with vehicle-treated or MycB-treated cells compared to untreated cells (Figure 2B, Supplementary Figure 3A, 3D). To evaluate trogocytosis in more detail, amoebae that performed ingestion were further subdivided into populations that had performed low versus high levels of trogocytosis (Figure 2C-F; Supplemental Figure 1F-H).

Low trogocytosis corresponds to amoebae that have ingested a low number of bites, while high trogocytosis corresponds to amoebae that have ingested a high number of bites (Figure S1H). When incubated with MycB-treated cells, significantly more amoebae performed low trogocytosis (Figure 2C, 2E; Supplementary Figure 3B, 3E). However, the proportion of amoebae that performed high trogocytosis was not significantly impacted (Figure 2D, 2F; Supplementary Figure 3C, 3F). Thus, impairing actin polymerization in human cells through treatment of human cells with MycB modestly impacts amoebic trogocytosis.

Next, the impact of human cell stiffness on amoebic phagocytosis was quantified by live microscopy of amoebae incubated with untreated, vehicle-treated, or MycB-treated cells (Figure 3A-B). There was no significant difference in the percentage of amoebae that performed phagocytosis on untreated cells and vehicle- or MycB-treated cells (Figure 3C). The number of internalized cells was also not significantly different (Figure 3D). Phagocytic cup formation was variable, but differences were insignificant between groups (Figure 3E). Thus, treatment of human cells with vehicle or MycB does not significantly impact amoebic phagocytosis.

**Figure 3.**
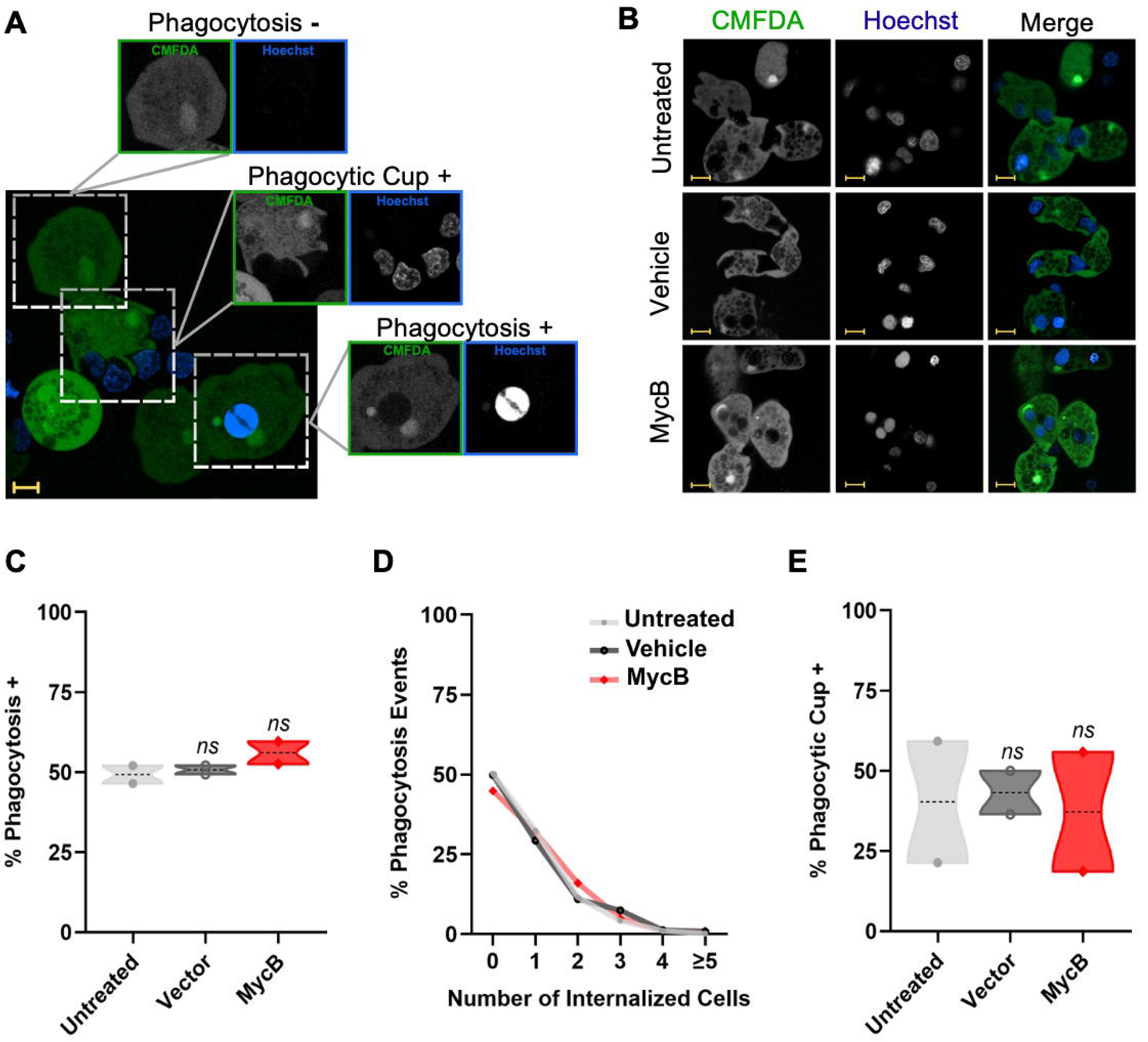
Pharmacological inhibition of human cell F-actin does not impair amoebic phagocytosis. Amoebae were stained with CMFDA and Jurkat cells (untreated, vehicle-treated, or MycB-treated) were stained with Hoechst. **A**, Representative annotated image that demonstrates scoring of ingestion events as: Phagocytosis - (no Hoechst-labeled Jurkat cells within a CMFDA-labeled amoeba), Phagocytic Cup + (a Hoechst-labeled Jurkat cell within a CMFDA-labeled phagocytic cup), and Phagocytosis + (one or more Hoechst-labeled Jurkat cells within a CMFDA-labeled amoeba). Scale bar, 10 μm. **B**, Representative images of amoebae ingesting untreated, vehicle-treated, or MycB-treated Jurkat cells. Total amoebae n = 1,513 amoebae total (503 amoebae with untreated Jurkats, 484 amoebae with vehicle-treated Jurkats, and 480 amoebae with MycB-treated Jurkats) .Scale bar, 10 μm. **C**, Percentage of amoebae with Phagocytosis + events, n = = 783 (untreated = 250; vehicle-treated = 243; MycB-treated = 290). Data shown are mean percentages of whole Phagocytosis + values. **D**, Percentage of amoebae with total phagocytosis events (Phagocytosis + or Phagocytosis -), broken down by the number of internalized Jurkat cells (0 to ≥5). Data shown are mean percentages of all phagocytosis event values described in panel C. **E**, Percentage of Phagocytic Cup + amoebae, n = 662 (untreated = 218; vehicle-treated = 222; MycB-treated = 222). Statistical analyses were performed using ordinary one-way ANOVA with Dunnett’s multiple comparisons tests.

### CRISPRi knockdown of Rho-pathway genes in Jurkat T cells

Reduced amoebic trogocytosis of MycB-treated human cells suggested that the nibbling was influenced by the human cell F-actin cytoskeleton, though these impacts on trogocytosis were relatively modest and there were no significant impacts on amoebic phagocytosis. We reasoned that inhibition of F-actin with MycB might not provide sufficient longevity to reliably assay for impacts on amoebic ingestion. Therefore, to alter actin cytoskeletal dynamics in human cells without using inhibitors, CRISPRi was used to knock down Rho-pathway genes that canonically regulate actin cytoskeletal tension (Figure 1). Seven different genes were individually targeted for knockdown: CFL1, MYPT1, Rac1, Rac2, RhoA, ROCK1, and ROCK2. Human Jurkat T cells were stably transduced with a construct for dCas9-KRAB expression, and because the proportion of intracellular dCas9-KRAB is a key parameter for successful CRISPRi (94,95), we generated clonal lines to evaluate dCas9-KRAB expression. Western blotting analysis showed that dCas9-KRAB expression was variable among clonal lines (Figure 4A; Supplemental Figure 5A). The clonal line 2D10 was selected for subsequent experimentation to ensure sufficient dCas9-KRAB expression and reduced cell-to-cell variability in knockdown efficacy.

**Figure 4.**
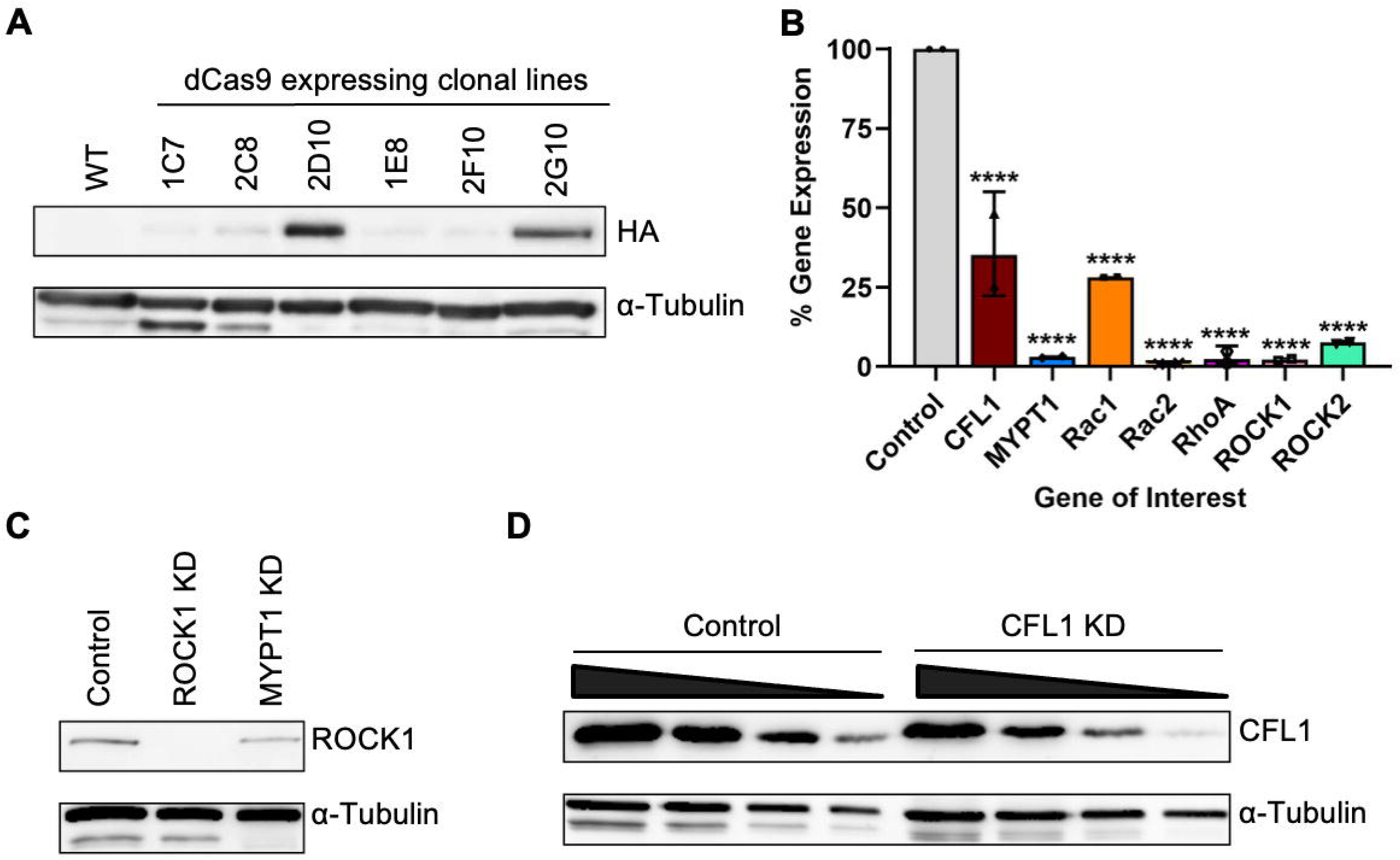
Targeted genes are efficiently knocked down in Rho-pathway knockdown mutants.A, Western blotting analysis of dCas9 expression among Jurkat T cell clonal lines that were stably transduced with a construct for expression of HA-tagged dCas9. Wild-type Jurkat T cells were used as a control. Western blots were probed with HA antibody (top) or alpha-tubulin (α-tubulin) antibody as a loading control (bottom). The clonal line “2D10” was used for subsequent experiments in which constructs for sgRNA expression were stably transduced. **B,** RT-qPCR analysis of CRISPRi knockdown mutants. Jurkat 2D10 cells stably transduced with a construct for expression of sgRNA corresponding to the ANPEP gene, which is not expressed in Jurkat cells, were used as a control. Expression of individual target genes in knockdown mutants was compared to the expression of the same gene in ANPEP control cells using unpaired, two-tailed t-tests with Welch’s correction, p<0.0001 (****). **C,** Western blotting analysis of ANPEP control, ROCK1 and MYPT1 knockdown mutants. Blots were probed with ROCK1 antibody (top) or α-tubulin antibody as a loading control (bottom). **D,** Western blotting analysis of ANPEP control cells and CFL1 knockdown mutants. Total protein samples were serially diluted. Blots were probed with CFL1 antibody (top) or α-Tubulin antibody as a loading control (bottom).

sgRNA sequences targeting CFL1, MYPT1, Rac1, Rac2, RhoA, ROCK1, and ROCK2 were individually and stably transduced in the 2D10 clonal line. A sgRNA targeting Alanyl Aminopeptidase (ANPEP), a gene not expressed in Jurkat T cells, was stably transduced as a control. RT-qPCR assays showed that all target genes were successfully knocked down in the corresponding CRISPRi knockdown mutants (Figure 4A, Supplementary Figure 4A, 4B). Western blotting analysis corroborated these findings (Figure 4C-D; Supplemental Figure 5B-C). For example, ROCK1 protein was essentially undetectable in ROCK1 knockdown mutants (Figure 4C; Supplemental Figure 5B), consistent with near complete knockdown of ROCK1 expression. CFL1 protein levels were reduced in CFL1 knockdown mutants relative to control cells (Figure 4D; Supplemental Figure 5C), consistent with partial knockdown of CFL1 expression.

### F-actin organization is abnormal in Rho-pathway knockdown mutants

All Rho-pathway CRISPRi knockdown mutants exhibited abnormal morphology of the F-actin network compared to control cells (Figure 5). Control cells were primarily spherical in shape and cortical F-actin networks were relatively homogenous and symmetrically distributed. Surface protrusions such as short filopodia and small membrane ruffles were also observed, which is characteristic of rested, wild-type Jurkat T cells (96,97). Surface protrusion patterns were similar in ROCK1 knockdown mutants, though filopodial protrusions were comparably thicker, branched, and more expansive in the mutant. ROCK1 knockdown also resulted in oblong-shaped cells with sparse central actin networking. Less total F-actin networking was apparent in both Rac1 and RhoA knockdown mutants, and both mutant cell lines retained spherical shapes. Small ruffles and sporadic, asymmetric filopodia were also prevalent. Rac2 and ROCK2 knockdown mutants formed expansive distal lamellipodial protrusions along the cortex, which were more pronounced and rosette-like in the ROCK2 knockdown mutants, and Rac2 mutants displayed especially sparse proximal nuclear F-actin. CFL1 knockdown mutants were elongated in shape with dense peripheral and distal F-actin networks composing prominent distal, asymmetric ruffling near the lamellipodia. In CFL1 knockdown mutants, filopodia were sparsely present. In contrast, MYPT1 knockdown mutants exhibited round cortical F-actin structures with condensed, ring-like distal F-actin networking. In MYPT1 knockdown mutants, peripheral and central actin networks were sparse, and cortices held few protrusions. Additionally, to determine if the cytoskeletal microtubule network was impacted in Rho-pathway knockdown mutants, immunofluorescence analysis were used to visualize α- and β-tubulin, and there were no apparent differences between the knockdown mutants (Supplemental Figure 6).

**Figure 5.**
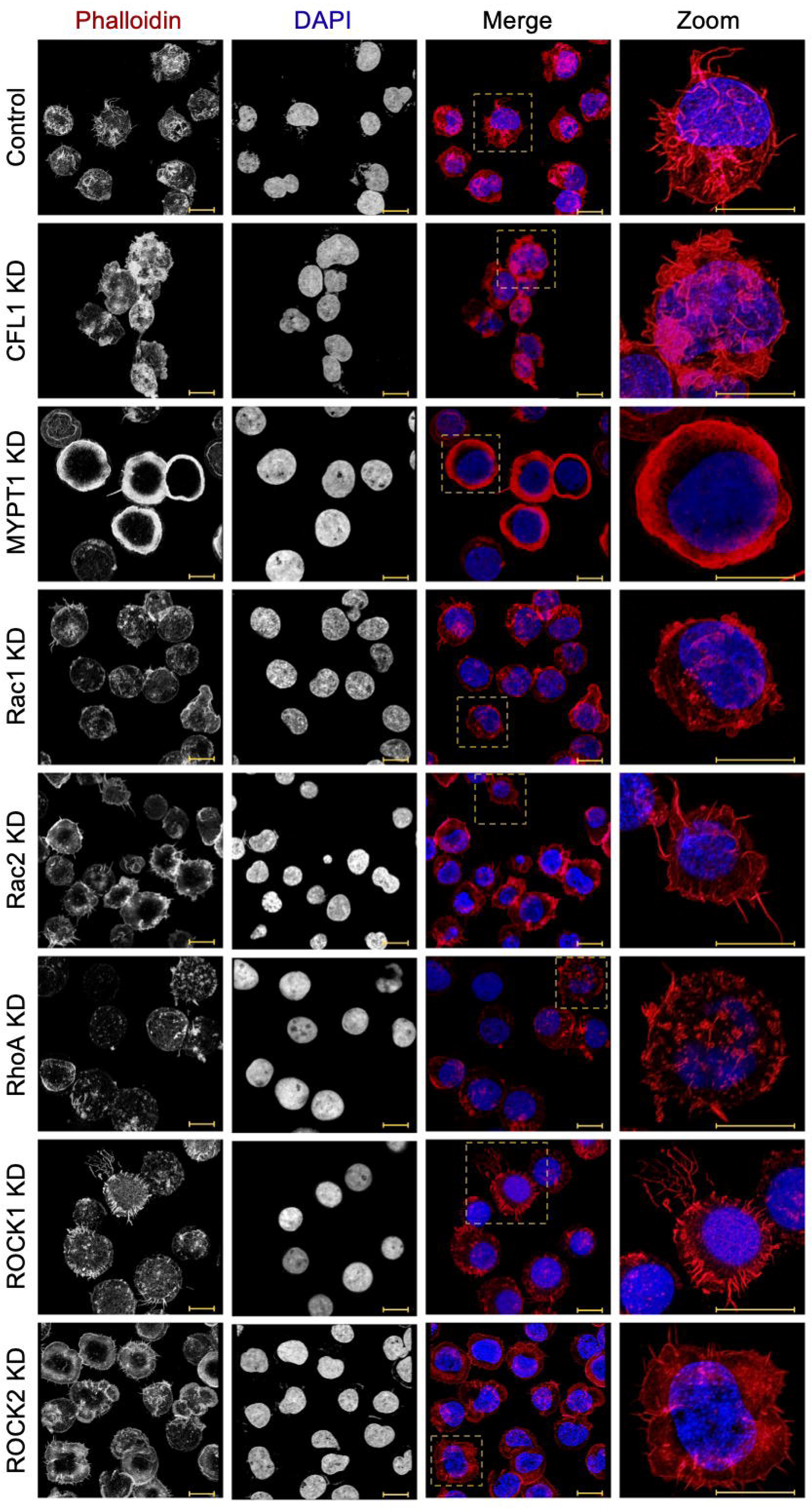
F-actin organization is abnormal in all Rho-pathway knockdown mutants. ANPEP control and Rho-pathway knockdown mutants were stained with phalloidin to label F-actin (red) and stained with DAPI to label DNA (blue). Images are representative of >50 cells imaged for each cell line. Scale bar (yellow), 10 μm.

### Amoebae perform reduced trogocytosis of all Rho-pathway knockdown mutants

Next, to ask if abnormal F-actin organization in Jurkat T cells would impact amoebic trogocytosis, amoebae were incubated with Rho-pathway knockdown mutants or control cells, and imaging flow cytometry was used to quantify amoebic trogocytosis (Figure 6; Supplemental Figure 1). Amoebic attachment to all knockdown mutants was equivalent to control cells (Supplemental Figure 7), and there was also no significant difference in the % Internalization+ for amoebae incubated with Rho-pathway knockdown mutants compared to control cells (Figure 6A, 6D, 6G; Supplemental Figure 8A, 8D). These data show that, overall, the same proportion of amoebae performed trogocytosis on all tested cell lines. However, when incubated with Rho-pathway knockdown mutants, more amoebae performed low trogocytosis (Figure 6B, 6E, 6H; Supplemental Figure 8B, 8E), showing that a greater proportion of amoebae ingested a low number of bites. Conversely, when incubated with Rho-pathway knockdown mutants, fewer amoebae performed high trogocytosis (Figure 6C, 6F, 6I; Supplemental Figure 8C, 8F), demonstrating that a smaller proportion of amoebae ingested a high number of bites. Overall, these findings show that the efficiency of amoebic trogocytosis was impacted, since amoebae ingested fewer bites of all Rho-pathway knockdown mutants compared to control cells.

**Figure 6.**
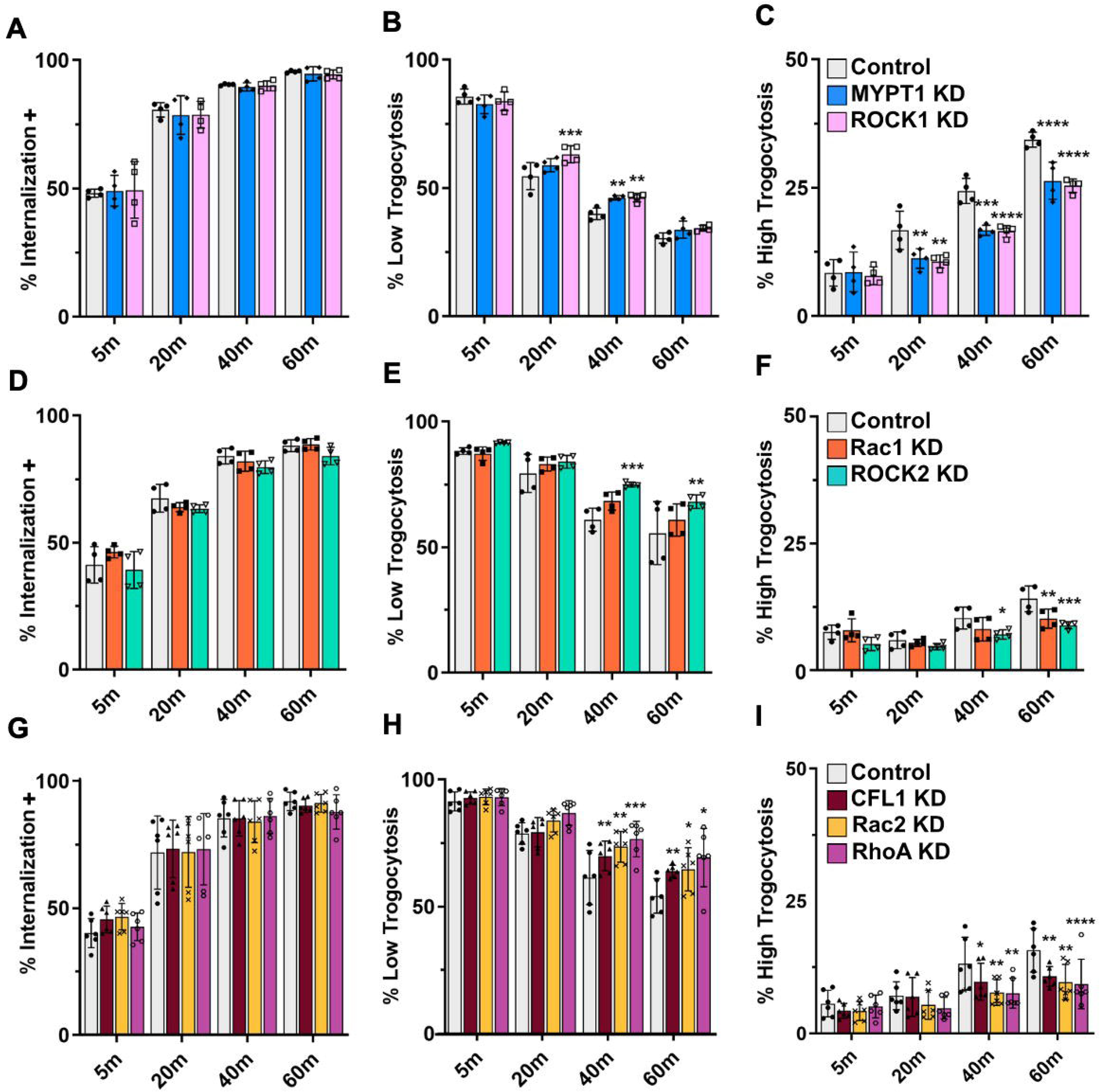
Amoebic trogocytosis of Rho-pathway knockdown mutants is impaired. A-I, Amoebae were stained with CMFDA and Jurkat cells (ANPEP control or Rho-pathway knockdown mutants) were stained with DiD. Cells were co-incubated for 5, 20, 40 or 60 minutes, and amoebic trogocytosis was quantified using imaging flow cytometry (see Figure S3). The percentage of amoebae that had performed ingestion was quantified (as % Internalization; **A**, **D**, **G**). Among amoebae that had performed ingestion, the level of trogocytosis was separated into amoebae with a low level of bites (as % Low Trogocytosis; **B**, **E**, **H**) or a high level of bites (as % High Trogocytosis; **C**, **F**, **I**). Data represent two to three separate biological replicates with two technical replicates per condition, for a total of four to six replicates per condition. Data are organized into separate panels (**A-C**, **D-F**, and **G-I**) according to experimental groups performed simultaneously. Statistical analyses were performed using ordinary 2-way ANOVA with Dunnett’s multiple comparisons test. **J-K,** Representative images of amoebae that performed low or high levels of trogocytosis. Brightfield (BF), CMFDA (green), DiD (red) and merged images are shown. Image numbers represent the order in which they were acquired during imaging flow cytometry. Scale bar, 10 μm.

### Amoebae perform increased phagocytosis of MYPT1 knockdown mutants and decreased phagocytosis of ROCK1 knockdown mutants

To determine if abnormal F-actin organization in Jurkat T cells would impact amoebic phagocytosis, amoebae were incubated with Rho-pathway knockdown mutants or control cells, and live microscopy was used to quantify amoebic phagocytosis (Figure 7). There was no significant difference in the percentage of amoebae that performed phagocytosis of Rac2 or RhoA knockdown mutants (Figure 7B). Significantly more amoebae performed phagocytosis of MYPT1 knockdown mutants, and fewer amoebae performed phagocytosis of ROCK1 knockdown mutants (Figure 7B). Interestingly, the increased amoebic phagocytosis of MYPT1 knockdown mutants was limited to events in which one Jurkat cell was ingested (Figure 7C). When phagocytic cups were quantified, amoebae interacting with ROCK1 mutants had the most phagocytic cups, consistent with stalled phagocytosis (Figure 7D). These findings show that only amoebic phagocytosis of MYPT1 and ROCK1 knockdown is altered, while amoebic phagocytosis of other tested Rho-pathway knockdown mutants is normal. Thus, amoebic phagocytosis efficiency is impacted by the stiffness of the target cell.

**Figure 7.**
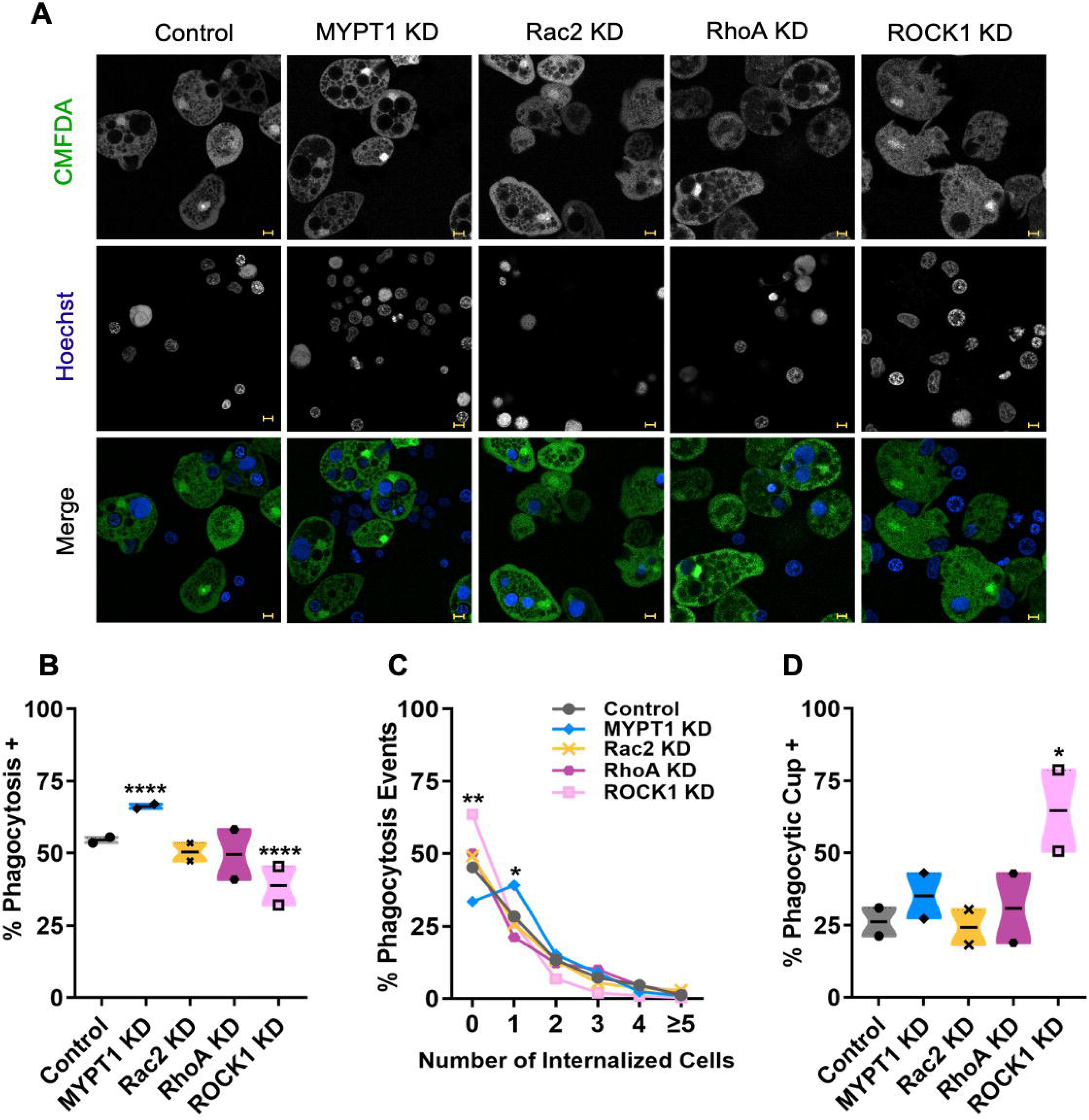
Amoebae perform increased phagocytosis of MYPT1 knockdown mutants and decreased phagocytosis of ROCK1 knockdown mutants. Amoebae were stained with CMFDA and Jurkat cells (ANPEP control or Rho-pathway knockdown mutants) were stained with Hoechst. **A**, Representative images of amoebae ingesting Rho-pathway knockdown mutants. Total amoebae n = 2,693 (Control = 826, MYPT1 KD = 463, Rac2 KD = 480, RhoA KD = 426, and ROCK1 KD = 498). Scale bar,10 μm. **B**, Percentage of amoebae with Phagocytosis + events, n = 1,398 (Control =452, MYPT1 KD = 308, Rac2 KD = 244, RhoA KD = 213, ROCK1 KD = 181). Data shown are mean percentages of whole Phagocytosis + values. **C**, Percentage of amoebae with total phagocytosis events (Phagocytosis + or Phagocytosis -), broken down by the number of internalized Jurkat cells (0 to ≥5). Data shown are mean percentages of all phagocytosis event values described in panel B. **D**, Percentage of Phagocytic Cup + amoebae. Statistical analyses were performed using ordinary one-way ANOVA with Dunnett’s multiple comparisons tests.

### Cell stiffness is significantly altered in Rho-pathway knockdown mutants

Atomic Force Microscopy-based single cell compression was used to evaluate the rigidity of control and knockdown mutants (Figure 8, Figure S9). Force *vs.* Relative Deformation profiles were obtained for successful compressions of individual cells in each cell line (Figure 8A-J). High consistency was observed among profiles, and in each profile, force increased with cellular deformation during initial contact and compression, i.e., ε = 0-40%. Stress peaks corresponded to the rupture of the cellular membrane appeared at an average ε = 60%. Continued applied compressive force induced further deformation, i.e., nuclear deformation, and the observed profiles are consistent with previously reported single T cell mechanics (98).

**Figure 8.**
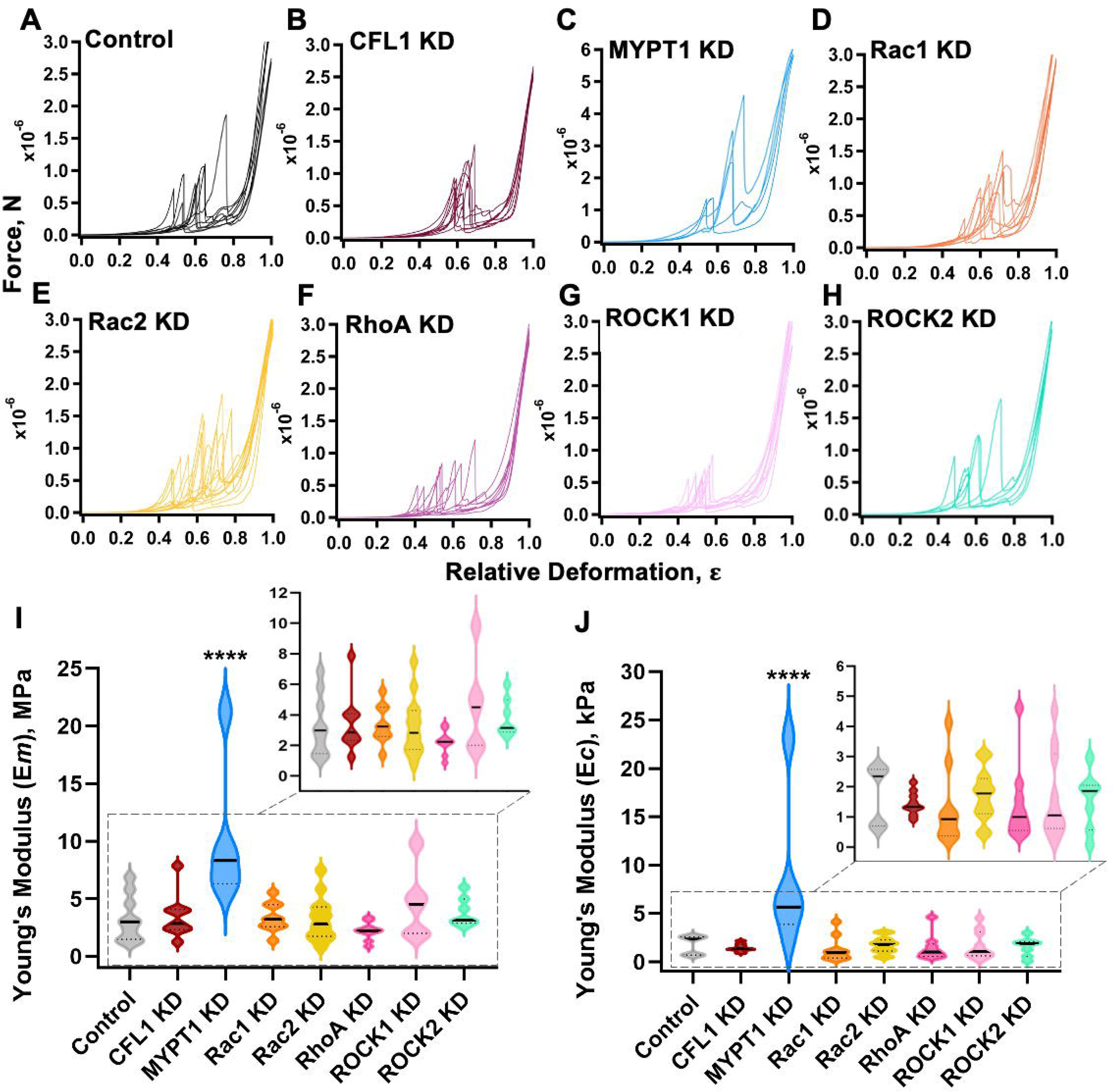
Knockdown of MYPT1 increases membrane and cytoskeletal rigidity, while other Rho-pathway mutants show decreased cytoskeletal rigidity. AFM was used to assess the stiffness of Rho-pathway knockdown mutants and ANPEP control cells. **A-H**, Force vs. Relative Deformation profiles for all analyzed cell lines. Data shown are ∼10 successful single cell compressions per group, total n =71 (Control = 9 , CFL1 KD = 9 , MYPT1 KD = 5, Rac1 KD = 9, Rac2 KD = 13 , RhoA X = 9, ROCK1 KD = 10 , ROCK2 KD = 7). **I,** Young’s moduli ( ) analyses for membrane stiffness. MYPT1 knockdown membrane rigidity is significantly greater than control. **J**, Young’s moduli ( ) analyses for cytoskeletal stiffness. MYPT1 knockdown cytoskeletal rigidity is significantly greater than control. Statistical analyses were performed using ordinary one-way ANOVA with Dunnett’s multiple comparisons tests.

Mechanical profiles for all knockdown mutants (Figures 8B-H) were similar to control (Figure 8A) but observable variations between knockdown mutants and control were noted. Stress peaks for most CFL1 (Figure 8B, S12), MYPT1 (Figure 8C, S13), Rac1 (Figure 8D, S14), and Rac2 (Figure 8E, S15) knockdown mutants were observed at slightly greater ε compared to control (Figure 8A, S11), indicating that a relatively larger applied force was required to rupture knockdown mutants. Conversely, stress peaks for most RhoA (Figure 8F, S16), ROCK1 (Figure 8G, S17), and ROCK2 (Figure 8H, S18) knockdown mutants were observed at slightly reduced ε compared to control (Figure 8A, S11), showing that a relatively smaller applied force was required to rupture knockdown mutants.

Quantification of Young’s modulus (stiffness) of the cellular membrane, E_m_, and the cytoskeleton, E_c_, was carried out following the Hertzian mechanics model (99,100) and non-linear least square fitting (Figure S9A). Number of compressions (n), average cell height, E_m_, and E_c_ are summarized in Table 1. Compared to control, the cell membrane and cell cytoskeleton of MYPT1 knockdown mutants were significantly stiffer (Figure 8I, 8J, S10). An average trend was observed with other knockdown mutant lines, where E_m_ of RhoA knockdown mutants were lower than control and both ROCK1 and ROCK2 knockdown mutants were stiffer than control (Figure 8I, Figure S10C). E_c_ for all knockdown mutants except for MYPT1 knockdown appeared lower than control (Figure 8J, Figure S10D). Thus, cell stiffness is significantly increased in MYPT1 knockdown mutants and variably decreased in other Rho-pathway knockdown mutants.

**Table 1.** Summary of AFM data.

| Cell Line | n | Height (µm) | E<sub>m</sub> (MPa) | E<sub>c</sub> (kPa) |
|---|---|---|---|---|
| Control | 9 | 8.9 ± 1.2 | 3.3 ± 2.0 | 1.73 ± 0.98 |
| CFL1 KD | 9 | 8.97 ± 0.79 | 3.5 ± 1.9 | 1.45 ± 0.39 |
| MYPT1 KD | 5 | 9.0 ± 1.1 | 10.3 ± 6.3 | 8.8 ± 8.2 |
| Rac1 KD | 9 | 7.8 ± 1.1 | 3.4 ± 1.3 | 1.3 ± 1.3 |
| Rac2 KD | 13 | 7.7 ± 1.2 | 3.22 ± 1.90 | 1.71 ± 0.86 |
| RhoA KD | 9 | 8.83 ± 1.21 | 2.22 ± 0.78 | 1.43 ± 1.31 |
| ROCK1 KD | 10 | 7.80 ± 0.84 | 4.70 ± 3.06 | 1.68 ± 1.46 |
| ROCK2 KD | 7 | 7.78 ± 0.92 | 3.83 ± 1.27 | 1.54 ± 0.98 |

For each cell line, the total number of analyzed cells (n) is indicated, along with the average cell height in µm. The average value of the calculated Young’s modulus (stiffness) of the cellular membrane, E_m_, and the cytoskeleton, E_c_, is shown, in MPa and kPa, respectively.

## Discussion

Amoebae performed quantitatively less trogocytosis on all Rho-pathway mutants compared to control cells, regardless of changes to F-actin architecture and overall stiffness. Amoebae performed more phagocytosis on stiffer MYPT1 knockdown mutants with relatively actin-rich cortexes, and less phagocytosis on less stiff ROCK1 knockdown mutants with relatively actin poor, but highly filamentous, cortexes. Thus, target cell stiffness impacts amoebic phagocytosis, but regulation of amoebic trogocytosis is more complex (Figure 9).

**Figure 9.**
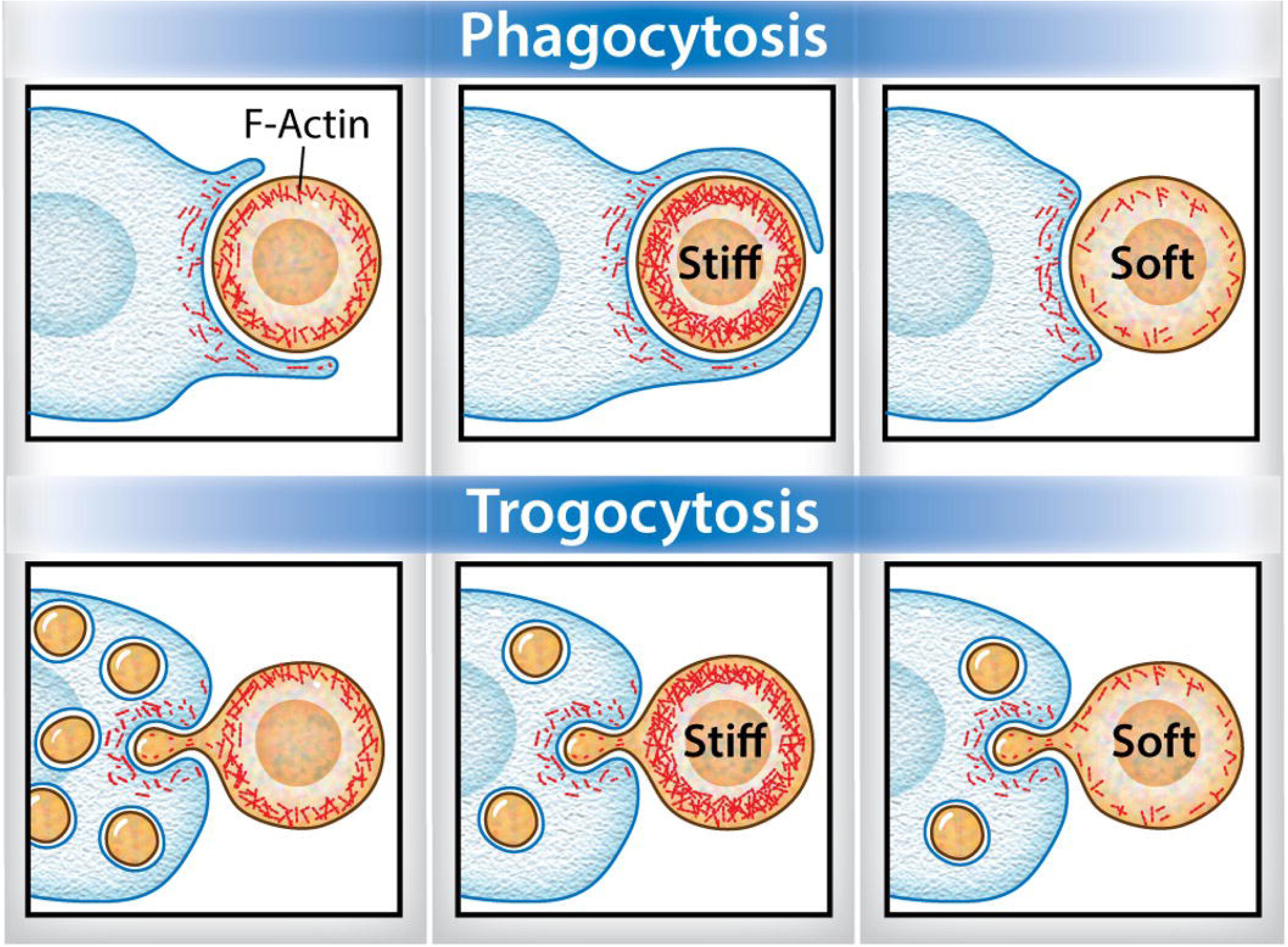
Human cell cytoskeletal rigidity differentially influences trogocytosis and phagocytosis. An amoeba (blue) is shown ingesting a human cell (orange), with F-actin indicated (red). Amoebae perform phagocytosis by attaching to a human cell and forming a phagocytic cup that surrounds the human cell. Phagocytosis efficiency is increased when amoebae ingest relatively stiff knockdown mutants. Phagocytosis efficiency is decreased when amoebae ingest relatively soft knockdown mutants, and more amoebae stall at the phagocytic cup stage. By contrast, amoebic trogocytosis efficiency is decreased with both relatively stiff and relatively soft knockdown mutants. Amoebae still perform trogocytosis but tend to take fewer bites of the stiffer and softer human cells, relative to control human cells. Image credit: Anita Impagliazzo.

We first sought to use pharmacological inhibition to interfere with human cell F-actin polymerization. This is challenging because inhibitors of the cytoskeleton typically target highly conserved aspects and therefore must be thoroughly washed out before incubating inhibitor-treated human cells with amoebae. A further challenge is that many inhibitors have half-lives that are shorter than the time needed to assay for amoebic ingestion. Treatment of human cells with MycB (87–91) led to inhibited amoebic trogocytosis, but phagocytosis was unaffected. These impacts were relatively modest, suggesting that the half-life of MycB might be too short to reliably assay for impacts on amoebic ingestion.

To produce stable, long-lasting changes to F-actin dynamics in human cells, we used CRISPRi to target Rho-pathway genes, which produced the expected phenotypic changes (65,66,63,101,102). Loss of F-actin stability was observed in Rac1 and RhoA knockdown mutants, which displayed thin cortical F-actin, minimal filopodia, and no significant membrane ruffling. Rac1 knockdown mutant phenotypes were consistent with the loss of Arp2/3-mediated F-actin stability (68,70,71,102–105,105). RhoA knockdown mutant phenotypes fit with the loss of mDia and formin-related filopodia formation (74,79,105–108). ROCK1 knockdown mutants displayed thin cortical actin networks and elongated filopodia, as expected with reduced F-actin stability, actomyosin contractility, and Rac1 antagonism (73,77,106,109). Rac2 and ROCK2 knockdown mutants maintained expansive distal lamellipodial protrusions, consistent with respective losses in F-actin polarization (110–114) and lamellipodia formation regulation (115–120). CFL1 knockdown mutants were asymmetric with accumulated cortical F-actin and large membrane ruffles, fitting with reduced actin turnover along the cortex (67,121–126). MYPT1 knockdown mutants had large, dense cortical actin and minimal membrane protrusions, consistent with reduced myosin-binding antagonism and increased actomyosin contractile interactions (78,124,127–132).

Fitting with the changes in F-actin dynamics, knockdown of Rho-pathway genes expectedly altered cytoskeletal rigidity, as assessed by AFM. At both the membrane, , and cytoskeletal, , level, MYPT1 knockdown mutants were significantly more rigid than control cells ROCK1 knockdown mutants were modestly more rigid at the membrane compared to control but were less rigid than control at the cytoskeletal level (Figure 8, Figure S10C-D). This increased membrane rigidity may be due to presence of elongated filopodial protrusions with the loss of cortical F-actin networking contributing to reduced cytoskeletal rigidity. Similarly, loss of F-actin networking membrane protrusions appeared to reduce both membrane and cytoskeletal rigidity of RhoA and Rac1 knockdown mutants. Membrane ruffles and lamellipodial protrusions did not appear to impact membrane stiffness in CFL1 and ROCK2 knockdown mutants, respectively, though apparent changes to cortical F-actin resulted in a modest reduction of cytoskeletal rigidity. Knockdown of Rac2 decreased membrane rigidity but did not disrupt cytoskeletal rigidity

In the AFM analyses, the Young’s moduli values were smaller than described previously (98). Variations in the location of stress peaks reflected single cell individuality, as T cells are known to have a strong membrane and a relatively weak cytoskeleton; thus, membrane ruptures under compression were expected (98). Additionally, variation within cell lines was possibly due to the biotinylation-streptavidin adherence of cells to the glass surfaces, as previous work did not employ coating. However, since all cells were prepared the same way, comparisons between cells are meaningful. While many of the changes that we detected were small in magnitude, this could be due to the inherently low throughput of AFM approaches, which precludes analysis of large numbers of cells. Additionally, because the mutants are knockdown rather than knockout mutants, there could be cell-to-cell variability in the level of gene knockdown, creating variability in the rigidity of individual cells.

Amoebic phagocytosis efficiency was impacted by target cell stiffness, where greater target cell stiffness correlated with increased phagocytosis, and reduced target cell stiffness correlated with reduced phagocytosis. This finding in *E. histolytica* fits nicely with studies in human macrophages, where phagocytosis was impacted by target stiffness. However, almost all studies of macrophage phagocytosis used artificially stiffened target cells or artificial targets (83–85).By contrast, we used native target cells, in which stiffness was carefully increased or decreased by individually knocking down Rho-pathway genes. In a recent study of macrophage phagocytosis of stiff vs. soft hydrogel microparticles, phagocytic cups closed quickly around stiffer microparticles, but tended to stall around softer microparticles (83). In our studies, amoebae that were ingesting relatively less stiff ROCK1 mutants had the most phagocytic cups. Thus, it is tempting to speculate that amoebic phagocytic cups also stall around softer target cells. While increased or decreased target cell stiffness correlated with increased or decreased amoebic phagocytosis, all changes in target cell stiffness correlated with decreased amoebic trogocytosis. Interestingly, trogocytosis efficiency was even impacted using knockdown mutants for which AFM was not sensitive enough to uncover significant defects in cell stiffness. It is also important to note that by using imaging flow cytometry to assay trogocytosis in our studies, we were also able to assay amoebic attachment to human cells, and there were no defects in amoebic attachment to any of the mutant human cells.

Thus, amoebae were still able to attach to mutant human cells and to initiate trogocytosis, but the efficiency of trogocytosis was impacted, since amoebae took fewer bites. Our findings highlight the importance of using native cells as targets and the importance of variably modulating stiffness (i.e., both increased and decreased) since the relationship between target stiffness and amoebic trogocytosis efficiency is not as simple as the study using glutaraldehyde fixed red blood cells suggested (55).

Beyond amoebae, studies exploring the impacts of target stiffness on trogocytosis are limited. In a recent study by Cornell et al, macrophages performed increased trogocytosis and decreased phagocytosis on cell lines with relatively low membrane tension, and conversely, decreased trogocytosis and increased phagocytosis on a cell line with relatively high membrane tension (85). It is important to note that there are many physiological differences between these cell lines, in addition to differences in membrane tension. Interestingly, macrophages predominantly performed phagocytosis on GUVs with higher membrane tension and predominantly performed trogocytosis on GUVs with lower membrane tension (85). Both the findings with cell lines and with GUVs contrast with our findings using native cells as targets for ingestion. It is interesting to speculate that these contradictory observations could be the result of the active, dynamic remodeling of the cytoskeleton in living cells.

In a recent study by Rollins et al., macrophages performed increased phagocytosis on suspension cell lines and increased trogocytosis on adherent cell lines (86). Interestingly, these findings with cell lines do not agree with the recent study by Cornell et al. (85). It is important to reiterate that there are many physiological differences between these cell lines, which renders the differences in macrophage trogocytosis/phagocytosis efficiency of these cell lines harder to interpret. Various approaches were used to increase or decrease target cell adherence, which clearly impacted macrophage phagocytosis, with little to no impact on trogocytosis (86). The only impact on trogocytosis was when SKOV3 cells were made artificially suspended, which led to a slight reduction in macrophage trogocytosis (86).

In our studies with amoebae, when amoebic trogocytosis efficiency was reduced, the magnitude of the defect was relatively small. Interestingly, this is in agreement with the recent studies of macrophage trogocytosis, where in all cases, trogocytosis defects were minor (86)(83). It is thus notable that, altogether, observed defects in trogocytosis are small in magnitude. This suggests that trogocytosis may be a robust process that is more difficult to override than phagocytosis, for which larger magnitude defects are more typically uncovered.

Our study shows, for the first time, that target cell cytoskeletal rigidity inversely correlates with *E. histolytica* phagocytosis, but does not inversely correlate with trogocytosis efficiency. Our experimental approach provides the first example in which trogocytosis and phagocytosis were studied using native target cells with variations in cytoskeletal stiffness, rather than artificial targets or artificially stiffened cells. These findings differ from the conclusions of studies of both amoebic and macrophage trogocytosis that used artificial targets or artificial cell stiffening (55,57,85). Our findings emphasize that the dynamic nature of the cytoskeleton in living cells increases the complexity of how target cell cytoskeletal stiffness influences trogocytosis. Finally, given the relevance of trogocytosis and phagocytosis to pathogenesis, in addition to shedding light on the burgeoning field of eukaryotic trogocytosis, this work extends knowledge of aspects of the host that influence the capacity of amoebae to perform ingestion processes that contribute to disease.

## Materials and Methods

### Cell Culture

*E. histolytica* HM1:IMSS (ATCC 30459) trophozoites were cultured as described previously (133–135). Briefly, amoebae were maintained at 35°C in TYI-S-33 medium supplemented with 80 U/ml penicillin, 80 µg/ml streptomycin (Gibco), 2.3% Diamond Vitamin Tween 80 solution (40X; Sigma-Aldrich), or prepared according to Diamond et al, 1978 (136,137), and 15% heat-inactivated adult bovine serum (Summa Life Sciences). Amoebae were maintained in T25 tissue culture flasks (Corning) or glass culture tubes and at 80% confluence. Amoebae were harvested for experimentation at approximately 80 to 90% confluence (133,134).

Human Jurkat T cells (ATCC TIB-152, clone E6-1) were cultured as described (133,134) at 37°C and 5% CO in RPMI 1640 medium with L-glutamine, without phenol red (Gibco), supplemented with 10 mM HEPES (Sigma-Aldrich), 100 U/mL penicillin and 100 µg/mL streptomycin (Gibco), and 10% heat-inactivated fetal bovine serum (Gibco). Jurkat cells were maintained in vented T25 or T75 tissue culture flasks between 1×10^5^ and 1×10^6^ cells/mL and passaged between 5×10^5^ and 2×10^6^ cells/mL. Jurkat cells were harvested for experimentation between 5×10^5^ and 2×10^6^ cells/mL.

Human embryonic kidney 293T/17 cells (ATCC CRL-11286) were cultured at 37°C and 5% CO in DMEM with glucose, sodium pyruvate, GlutaMAX, and phenol red (Gibco), supplemented with 10mM HEPES, 100 U/mL penicillin and 100 µg/mL streptomycin, and 10% heat-inactivated fetal bovine serum (Summa Life Sciences). 293T/17 cells were maintained in vented T25 tissue culture flasks and passaged or harvested when flasks reached 80 to 90% confluence. Cells were detached by using 0.25% (w/v) Trypsin-0.53M EDTA solution (EDTA) treatment.

For co-incubation experiments, cells were harvested and resuspended in medium M199 with Earle’s salts, L-glutamine, and 2.2 g/L sodium bicarbonate, without phenol red (Gibco) and supplemented with 5.7 mM L-cysteine, 25 mM HEPES, and 0.5% bovine serum albumin (Summa Life Sciences) (referred to as M199s) (133,134).

### DNA Constructs

The lentiviral transduction transfer plasmid SC0217_pU6-sgRNA ANPEP#1-puro (a gift from Sean R. Collins, University of California, Davis) was modified to replace puromycin with hygromycin. Restriction enzymes NheI-HF and EcoRI-HF (New England Biolabs) were used to remove the puromycin insert from the SC0217 backbone and cleanup was performed with the QIAquick gel extraction kit (QIAGEN). The hygromycin construct pEhEx_empty vector_HygR was PCR amplified using primers with overhangs corresponding to the SC0217 backbone (Table S1) Gibson assembly was performed to ligate the SC0217 backbone and the hygromycin insert and One Shot STBL3 *E. coli* (ThermoFisher) colonies were screened using restriction digest. The SC0217-Hyg construct was confirmed using Sanger sequencing (Table S1).

For creation of Jurkat cell CRISPRi knockdown mutants, single guide RNAs (sgRNAs) to target human CFL1, MYPT1, Rac1, Rac2, RhoA, ROCK1, and ROCK2 genes were selected from published data from the Human Genome-wide CRISPRi-v2 Libraries (gift from Sean R. Collins, University of California, Davis, and deposited by Jonathan S. Weissman, University of California, San Francisco; Addgene pooled libraries #83969, #1000000090) (138). The top three highest ranked sequences were chosen for each gene of interest. The SC0217-Hyg plasmid was linearized using two restriction enzymes, FastDigest Bpu1102I (BlpI) and FastDigest BstXI (ThermoFisher), then dephosphorylated with FastAP Thermosensitive Alkaline Phosphatase (ThermoFisher). Linearized SC0217-Hyg cleanup was performed with MagBio HighPrep PCR magnetic beads (MagBio). Oligos containing the sgRNA sequences with appropriate overhangs (Table S1) were annealed and phosphorylated with T4 Polynucleotide Kinase (New England Biolabs), prior to ligation to the SC0217-Hyg plasmid backbone using the Quick Ligation Kit (New England Biolabs). Bacterial colonies were screened and insertion of the sgRNA sequences was confirmed using Sanger sequencing.

### Lentiviral Transduction

To create Jurkat cells with constitutive production of dCas9, 293T/17 cells were first seeded at 3×10^5^ cells/mL in 6-well plates and grown to 80-90% confluency. 293T/17 cells were then transfected with 0.77 μg of the envelope plasmid SC0058-TEN pMD2.G Envelope 2nd G, 1.51 μg of the packaging plasmid SC0059-TEN pCMV delta R8.2 Packaging, and 2.32 μg of the transfer plasmid SC0575_phr-ucoe-ef1a-dCas9-ha-2xnls-xten80-krab-p2a-Bls (gifts from Sean R. Collins, University of California, Davis; SC0575 was modified from a plasmid created in the laboratory of Jonathan S. Weissman, University of California, San Francisco, Howard Hughes Medical Institute). All lentiviral constructs were added to Opti-MEM Reduced Serum Medium (Gibco) and TransIT-2020 Transfection Reagent (Mirus) for transfection, and transfected 293T/17 cells were incubated at 37°C for 48 hours. Following incubation, the lentiviral supernatant was harvested and added at a 1:1 ratio to wildtype Jurkat cells at 1×10^6^ cells/mL with 8 μg/ml Polybrene (ThermoFisher). Jurkat cells were incubated with lentiviral supernatants for 24 hours at 37°C, then placed under blasticidin (Gibco) selection at 10 μg/ml for 2 weeks. Following selection, transduced cells were cloned using limiting dilution in 96-well plates.

To generate CRISPRi knockdown mutants, the SC017-Hyg transfer vectors containing sgRNA sequences were each transduced into the dCas9-expressing clonal line 2D10, using the same process described above, but replacing the SC0575 plasmid with each SC017-Hyg-sgRNA plasmid. Transduced cells were placed under hygromycin (Invitrogen) selection at 800 μg/ml for 2 weeks. Mindful of the potential for compensatory changes, knockdown mutant lines were not cloned by limiting dilution, and all experimentation was performed as quickly as possible (within two weeks) after obtaining stably transduced cells. During these studies, each individual knockdown mutant line was generated by performing at least two independent lentiviral transductions, and the findings were consistent across independently generated knockdown mutant lines.

### RNA Extraction and RT-qPCR

RNA was extracted from Jurkat cells using the Direct-Zol RNA MiniPrep Plus kit (Zymo) as described previously (133). For each Jurkat cell line, RNA extraction was performed on two separate days, per cell line to form biological replicate groups. SuperScript Reverse Transcription II (Invitrogen) was performed as per the manufacturer instructions to obtain cDNA (139).

qPCR primers were designed and primer efficiency was empirically validated as previously described (135). A dilution series of wild-type Jurkat cell cDNA was (1:10, 1:100, 1:1,000, and 1:10,000 dilutions) was used as the template for qPCR amplification. The resulting cycle threshold (Ct) values were plotted against the log of the cDNA concentration, and efficiency was calculated using the linear equation for best fit line:

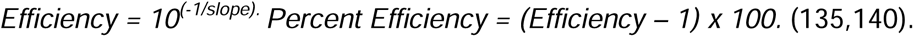

Primer efficiency values were deemed acceptable if between 90 and 110% (Table S1 and Figure S1). All assays used LightCycler 480 SYBR Green I Master Mix, 2X Conc. (Roche) and reactions were carried out using the LightCycler 480 Instrument II (Roche). Two technical replicates were carried out per RNA sample, to amplify the target gene and the housekeeping genes GAPDH and RPLP0. Relative expression of the target gene was assessed using the 2^-ΔΔCt^ method (135,141,142).

### Protein Preparation and Western Blotting

Jurkat cell protein extracts were prepared by incubation in lysis buffer containing 50 mM Tris-Cl at pH 7.4, 150 mM NaCl, 1% NP40, 0.5% DOC, 0.1% SDS, and 1X SigmaFast protease inhibitor cocktail (Sigma). Laemmli sample buffer was added to 1X final concentration and samples were denatured for 5 minutes at 100°C. To assess heterogeneity of dCas9-producing clonal lines, HA-tagged dCas9 was detected with a mouse monoclonal antibody directed to the HA epitope (Invitrogen, #26183). ROCK1 knockdown was evaluated using a rabbit monoclonal antibody directed to ROCK1 (Abcam, #ab134181), and CFL1 knockdown was evaluated using a rabbit polyclonal antibody directed to CFL1 (Abcam, #ab42824). Protein loading was assessed by using a mouse monoclonal antibody directed to alpha-tubulin (Sigma-Aldrich, #T5168). Anti-mouse and anti-rabbit secondary antibodies conjugated to Alexa Fluor 647 (Jackson Immunology, goat anti-mouse #115-605-062 and alpaca anti-rabbit #611-605-215) were used, and fluorescent images were acquired on a GE Amersham Imager 600. All samples were loaded at 20 μg total protein, except for CFL1-knockdown analysis, where samples were serially diluted.

### Phalloidin Staining

In some experiments, cells were treated with 1 µM mycalolide B (MycB) (FujiFilm Wako) or a 10% DMSO (vehicle control) for 10 minutes and washed three times in 1X PBS or M199s. To visualize filamentous actin, Jurkat cells were first incubated at 1×10^7^ cells in 1X PBS on 12 mm diameter glass coverslips (Electron Microscopy Services) that were coated with 0.1 mg/ml Poly-L Lysine (Millipore), in a 24-well plate (Corning). Cells were fixed with ice-cold 4% paraformaldehyde in 1X PBS for 15 minutes at room temperature, washed in 1X PBS, and then permeabilized with 0.5% Triton X-100 in 1X PBS for 5 minutes at room temperature. Cells were stained with 100 nM Acti-stain 488 Phalloidin (Cytoskeleton Inc., #PHDG1) for 30 minutes at room temperature. Cells were subsequently washed and stained with 100 nM DAPI (Millipore Sigma, #D9542) for 5 minutes at room temperature and washed again. Coverslips were mounted on glass slides (VWR) with 3 μl VectaShield (Vector Laboratories) and stabilized with a layer of clear nail polish around the edges.

Images were collected using a 63X objective on an Axio Observer Z1/7 (Zeiss LSM 980) with Airyscan2. Airyscan functionality was set to Super Resolution 8.3 (3D, Auto) with bidirectional scanning and z-stack acquisition to capture up to 24 z-stack slices at 1.05 - 1.35 μm per image. Start and end points for z-stack acquisitions were manually determined for each image, depending on the range of protruding actin filaments. Images were processed with Zeiss Zen imaging software, where 3-D images were then constructed using the precise maximum intensity projection of the overlaid z-stacks. Final image processing was performed using Imaris 9 (Oxford Instruments), which included a denoise step with a standard length of 1.5-2 to reduce background, and false recoloring of actin filaments (488 green to firestrm LUT red).

### Trogocytosis Assays

Amoebae and Jurkat cells were washed in M199s and independently labeled with fluorescent dyes before co-incubation. In some experiments, Jurkat cells were treated with 1 µM mycalolide B (MycB) (FujiFilm Wako) or a 10% DMSO (vehicle control) for 10 minutes and washed three times in M199s. Amoebae were labeled with CellTracker Green CMFDA (Invitrogen) at 200 nM in M199s for 15 minutes at 35°C and then washed in M199s and resuspended at 4×10^5^ cells/ml. Jurkat cells were labeled with sulphonated 1,1’-Dioctadecyl-3,3,3’,3’-Tetramethylindodicarbocyanine, 4-Chlorobenzenesulfonate Salt (DiD) (Invitrogen) at 5 μM in M199s for 5 minutes at 37°C, then 10 minutes at 4°C, washed in M199s, and resuspended at 2×10^6^ cells/ml. CMFDA-labeled amoebae and DiD-labeled Jurkat cells were combined at a 1:5 ratio in M199s and incubated at 37°C for 5, 20, 40, or 60 minutes, with two replicates per timepoint. As a control, amoebae and Jurkat cells were incubated independently for 60 minutes. Samples were transferred to ice and stained with Live/Dead fixable violet (Invitrogen) at 1 μl/ml for 30 minutes in the dark and fixed with 4% paraformaldehyde in 1X PBS for 30 minutes at room temperature. Samples were washed in 1X PBS, transferred to round-bottom 96-well plates (Corning), and sealed with pierceable plate covers (X-Pierce). Data were collected using an AutoSampler on an Amnis ImageStreamX Mark II flow cytometer using a 40X objective. 10,000 events were collected for each sample and data were analyzed using Amnis IDEAS software (Supplemental Figure 1).

### Phagocytosis Assays

Amoebae and Jurkats were washed in M199s. In some experiments, Jurkat cells were treated with 1 µM mycalolide B (MycB) (FujiFilm Wako) or a 10% DMSO (vehicle control) for 10 minutes and washed three times in M199s. Amoebae were labeled with CMFDA as described for trogocytosis assays, washed in M199s, and resuspended at 2×10^6^ cells/ml. Jurkat cells were labeled with 5 μg/mL Hoechst (ThermoFisher) at 37°C for 30 minutes, washed in M199s, washed in M199s, and resuspended at 1×10^7^ cells/ml. Amoebae and Jurkat cells were co-incubated at a 1:5 ratio on 25 mm coverslips (Chemglass) in an Attofluor Cell Chamber (ThermoFisher) at 35°C. Cells were allowed to settle for 4 minutes prior to image acquisition. For each co-incubation period, image acquisition did not surpass 20 minutes total. Images were collected using a 40X objective on an Axio Observer Z1/7 (Zeiss LSM 980) with Airyscan2. Airyscan functionality was set to SR-8Y (2D, auto) with bidirectional scanning, capturing single-plane fields at an average of 1.22 seconds.

For each knockdown mutant, phagocytosis assays were performed on two separate days. 306 image files (105 images of amoebae with ANPEP control, 50 images of amoebae with MYPT1 knockdown mutants, 53 images of amoebae with Rac2 knockdown mutants, 48 images of amoebae with RhoA knockdown mutants, and 50 images of amoebae with ROCK1 knockdown mutants) were processed with Zeiss Zen imaging software, then blindly renamed using a single substitution cypher with a randomized number naming permutation (Python).

Image files were individually analyzed using Zeiss Zen imaging software. Amoebae that were in focus and fully contained within the field of view were analyzed. Phagocytosis positive events were defined by the presence of a Hoechst-labeled Jurkat cell nucleus within a distinct phagosome (lacking CMFDA labeling) inside of an amoeba. Following manual image analysis, blinding was removed from files. In pharmacological treatment experiments, a total of 1,513 amoebae were analyzed (503 amoebae with untreated Jurkats, 484 amoebae with vehicle-treated Jurkats, and 480 amoebae with MycB-treated Jurkats). In CRISPRi-associated experiments, a total of 2,693 amoebae were analyzed (826 amoebae with ANPEP control, 463 amoebae with MYPT1 knockdown mutants, 480 amoebae with Rac2 knockdown mutants, 426 amoebae with RhoA knockdown mutants, and 498 amoebae with ROCK1 knockdown mutants).

### Atomic Force Microscopy

Atomic force microscopy (AFM) was performed to evaluate Jurkat cell rigidity. Biotinylated Jurkat cells were covalently bound to Streptavidin-coated glass substrates to allow for single-cell compression without directly stiffening the cytoskeleton. 50 mm dishes (MatTek) were first prepared through silanization of the glass bottom substrates with 1 M NaOH (aqueous) for 20 minutes at room temperature, washed in molecular grade water, and air dried. Dishes were then incubated with 2% (3-Aminopropyl)triethoxysilane (APTES)(Fisher) in 95% Ethanol for 2 hours at room temperature, washed in molecular grade water, and air dried. Next, dishes were coated with 0.1 mg/mL Streptavidin (ThermoFisher) in 1X PBS pH 7.4 at 4°C overnight, washed in 1X PBS with gentle agitation on a LSE low speed orbital shaker (Corning), and stored in 1X PBS at 4°C for up to three days before AFM. On the day of AFM experimentation, Jurkat cells were washed three times in ice-cold 1X PBS and incubated in 10 mM Sulfo-NHS-SS-Biotin (ThermoFisher) for 30 min at room temperature. Cells were washed once in ice-cold 50 mM Tris-Cl, twice in ice-cold 1X PBS, and resuspended at 1×10^6^ cells/mL in ice-cold 1X PBS pH 7.4. Biotinylated Jurkat cells were incubated on Streptavidin-coated dishes for 1 hour at room temperature. Just before mounting the dishes for compression, non-bound Jurkat cells were gently removed by aspiration, and 1X PBS was replaced with warm, complete RPMI.

Atomic Force Microscopy (AFM)-based single cell compression was developed and reported by Lulevich *et al*. previously (143,98) (Figure S9A). Single-cell mechanics were measured with a MFP-3D AFM (Asylum Research Corporation) integrated with an inverted optical microscope IX50 (Olympus America) used to guide the AFM microsphere and visualize single cells for compression. Prior to experimentation, an Au-coated silicon cantilever AC240TS (Olympus America) was modified by gluing a 40 ± 2 μm diameter glass sphere (Duke Scientific Corporation) to its apex using Devcon Epoxy 5 min (ITW Performance Polymers). The spring constant of the modified cantilever was determined using the adding mass method (144). = 2.8 N/m in these experiments. Before single-cell compression, cells were visualized and inspected using a brightfield 60X oil immersion objective, from which the majority population phenotypes of cells were identified. Round-shaped cells represented the phenotypic majority across all cell lines and thus were chosen for compression. Additionally, the photodiode sensitivity was calibrated and a force curve was acquired on the bare glass region adjacent to the target cell before each compression to serve as a surface reference point (98,145).

Upon selecting a target cell for compression, the center of the nucleus was aligned with the center of the glass sphere. During compression, the modified cantilever probe moved toward the cell at 2 μm/s, until 100% compression was achieved. Retraction of the probe occurred at the same μm/s. The force versus z-movement were recorded to attain the Force versus Relative Deformation profiles, while the cellular responses were monitored and recorded through the 60X objective and a CCD camera (Sony Corporation of America). All AFM data were acquired and analyzed using Igor Pro 6.34A (WaveMetrics, Inc.).

### Quantification of Membrane ( ) and Cytoskeletal ( ) Young’s Moduli

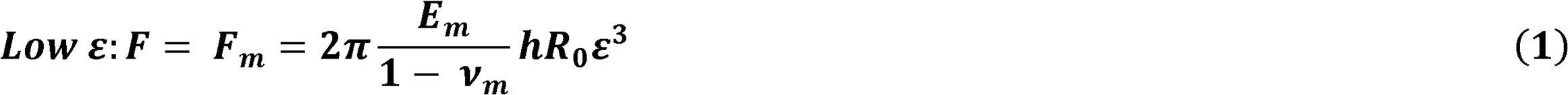

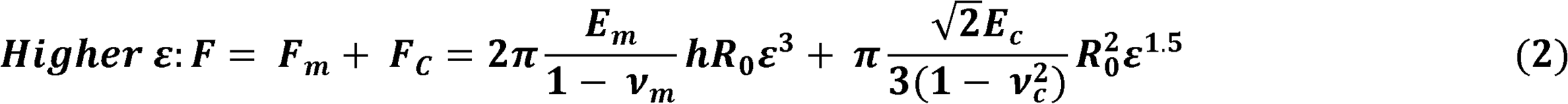

Evaluation of cell deformation during single cell compression was performed using a previously described incompressible fluid-filled balloon model (98,143). This model assumes the following: (a) during compression, the cell may be treated as a spherical membrane (balloon) filled with an incompressible fluid pressed between 2 parallel plates, (b) at low deformations, the cell membrane (shell of balloon) is impermeable, (c) a cell’s mechanical resistance arises from membrane stretching and cytoskeletal deformation with negligible contributions from other cellular components such as the nucleus at low deformations (98,143). Equations (1) and (2) were used to extract *E_m_* and *E_c_* respectively from Force *vs.* Relative Deformation profiles, where *F* is the total force, *F_m_* is the force contributed from cell membrane, *F_c_* is the force contributed from cytoskeleton, *E_m_* is the Young’s modulus of the cell membrane, *v_m_* is the Poisson ratio of the cell membrane *E_c_* is the Young’s modulus of the cytoskeleton, *v_c_* is the Poisson ratio of the cytoskeleton, *h*is the membrane thickness of the cell (Figures S10A, S10B), *R*_0_ is the radius of the cell before compression, and *ε* is the relative deformation of the cell *v_m_* was assumed to be 0.5, *v_c_* was assumed to be 0, *h* was assumed to be 4 *nm*, and *R*_0_ was assumed to be 5 *µm*.

First, *E_m_* was obtained by using Eq. (1) and the described assumptions to fit in the low *ε* region of the profile. Various *ε* regions were examined until the Eq. (1) fitness curve significantly deviated from the profile, and the low *ε* region was determined by the point at which the curve began to significantly deviate from the profile. Then, was obtained by using Eq. (2) with *E_m_* from Eq. (1) and the assumptions to fit in the higher *ε* region of the profile. Various *ε* regions were examined using Eq. (2) to determine the high *ε* region, as described with low *ε*. Supplemental Figure S9B-D shows a representative profile of a CFL1 knockdown cell, where the box represents the region of curve fitting. In this example, *E_m_* = 3.83 MPa (*x*^2=2.69×10^−16^) and = 1.68 kPa (*x*^2=6.73×10^−15^). This line fitting procedure was applied to all cells that underwent successful compressions.

### Statistical Analyses

Analyses were performed in Prism (GraphPad). Linear regression with best-fit values ± SE were applied to independent qPCR primer validation analyses. Nonlinear regression with line fitting was applied to atomic force microscopy data. Unpaired two-tailed t-tests with Welch’s correction were performed on normalized ΔΔCt qPCR data. Ordinary two-way ANOVA with Dunnett’s multiple comparisons tests were used in analyzing RT-qPCR, trogocytosis assays, and atomic force microscopy data. Fisher’s exact tests were used in analyzing yes/no parameters in phagocytosis assays. For all applicable statistical analyses, *ns,* P > 0.05; *, P ≤ 0.05; **, P ≤ 0.01; ***, P ≤ 0.001; ****, P ≤ 0.0001).

## Acknowledgments

We thank Anita Impagliazzo of Anita Impagliazzo Medical Illustration for the images in Figure 1 and Figure 9. We thank Sean Collins for the plasmid backbones and the protocol for lentiviral transduction. We thank Maura Ruyechan for modifying the lentiviral transduction transfer plasmid to replace puromycin with hygromycin. Imaging flow cytometry and microscopy were performed using shared instrumentation in the UC Davis MCB Light Microscopy Imaging Facility, of which the Zeiss 980 LSM was funded by NIH S10OD026702. We thank Thomas Wilkop for technical training and support in the use of these instruments. We thank the members of our laboratory, as well as Sean Collins and Scott Dawson, for helpful discussions, and we thank Chun-Yi Wu for advice on atomic force microscopy. This work was supported by NIH R01AI146914 and NIH R21AI193905 awarded to K.S.R..

## Financial Disclosure Statement

The funders had no role in study design, data collection and analysis, decision to publish, or preparation of the manuscript.

## Conflict of Interest Statement

The authors have declared that no competing interests exist.

## Author Contributions

F.P.L.: Methodology, Formal analysis, Investigation, Visualization, Writing – original draft, and Writing – review and editing. T.W.: Formal analysis, Investigation, Writing – original draft. M.C.I.: Investigation. R.L.S.: Investigation. G-Y.L.: Resources, Formal Analysis, Writing – original draft. K.S.R.: Conceptualization, Funding acquisition, Project administration, Supervision, Visualization, Writing – original draft, and Writing – review and editing.

## Supplementary Table Legends

Table S1. Primers used in these studies.

Sequences of primers that were used in these studies, for cloning of plasmid constructs, Sanger sequencing, cloning of sgRNA, and qPCR. The purpose of each primer is indicated, along with the primer name and sequence.

## Supplementary Figure Legends

**Figure S1. Imaging flow cytometry gating for trogocytosis assays.**

Amoebae were stained with CMFDA and Jurkat cells were stained with DiD. Cells were co-incubated for 5, 20, 40 or 60 minutes, and amoebic trogocytosis was quantified using imaging flow cytometry. **A**, All images were assessed using the gradient RMS of the masked brightfield to remove out of focus images from analysis. **B**, Images in focus were gated using a masked brightfield spot count to remove images that contained multiple cells that were not touching. **C**, Images that contained single cells or cell clusters were gated by aspect ratio and intensity of channel 2, corresponding to CMFDA dye, to identify images that contained only one amoeba per image (Single Amoebae). **D**, Images that contained single amoebae were then gated by maximum pixel intensity for channel 11, corresponding to DiD dye, to identify images that contained Jurkat cell material (Jurkat+). **E**, In images that contained single amoebae and Jurkat cell material, the internalization of Jurkat cell material was analyzed (i.e., the extent to which DiD-labeled material was within the CMFDA-labeled amoeba), to identify amoebae that had internalized Jurkat cell material (Internalization+). **F**, Finally, amoebae with internalized Jurkat cell material were analyzed using bright detail intensity and maximum pixel intensity for channel 11, which assess the degree of fragmentation of the DiD signal. Events were gated as low, mid, and high, corresponding to the levels of trogocytosis. **G**, Representative gating of for low, mid, and high levels of trogocytosis across four time points (5, 20, 40, 60 min. of co-incubation). **H,** Representative gating for low, mid, and high trogocytosis analysis, with representative images (a-f) shown. The position of each image is indicated in the scatter plot. Brightfield (BF), CMFDA (green), DiD (red), and merged images are shown. Scale bar, 10 μm.

**Figure S2. Analysis of amoebic attachment to untreated, vehicle-treated, or MycB-treated human Jurkat T cells.**

Amoebae were stained with CMFDA and Jurkat cells (untreated, vehicle-treated, or MycB-treated) were stained with DiD. Cells were co-incubated for 5, 20, 40 or 60 minutes, and amoebic trogocytosis was quantified using imaging flow cytometry (see Figure 2B-F and Supplementary Figure 1). Images that contained single amoebae were then gated by maximum pixel intensity for channel 11, corresponding to DiD dye, to identify images that contained Jurkat cell material (Jurkat+).

**Figure S3. Normalized analysis of amoebic trogocytosis of MycB-treated human cells.** The trogocytosis assay data shown in Figure 1B – 1D were normalized such that amoebic ingestion of untreated human cells was set to 100%. Amoebae were stained with CMFDA and Jurkat cells (untreated, vehicle-treated, or MycB-treated) were stained with DiD. Cells were co-incubated for 5, 20, 40 or 60 minutes, and amoebic trogocytosis was quantified using imaging flow cytometry (see Figure S1). **A,** The percentage of amoebae that performed ingestion was quantified (as % Internalization+), and amoebic ingestion of untreated human cells was set to 100%. **B,** Among amoebae that had performed ingestion, the proportion of amoebae that contained a low level of bites was quantified (% Low Trogocytosis), and low trogocytosis of untreated human cells was set to 100%. **C,** Among amoebae that had performed ingestion, the proportion of amoebae that contained a high level of bites was quantified (% High Trogocytosis) and high trogocytosis of untreated human cells was set to 100. **D-F**, Data shown are the fold change of amoebic ingestion of MycB-treated and vehicle-treated human cells relative to amoebic ingestion of untreated human cells, which is set to 1 and represented by a dashed line. Data represent four separate biological replicates with two technical replicates per condition, for a total of eight replicates per condition. Statistical analyses were performed using ordinary 2-way ANOVA with Dunnett’s multiple comparisons test.

**Figure S4. Primer efficiency analysis of qPCR primers.**

**A**, Primer efficiency analysis of housekeeping genes GAPDH and RPLP0. **B**, Primer efficiency analysis of genes of interest explored in this study, Data shown are plotted qPCR Ct values (Y axis) vs. log cDNA (X axis).

**Figure S5. Full Western blot images to accompany Figure 4**.

**A**, Western blotting analysis of dCas9 expression among Jurkat T cell clonal lines that were stably transduced with a construct for expression of HA-tagged dCas9. Wild-type Jurkat T cells were used as a control. Western blots were probed with HA antibody (right) or alpha-tubulin (α-tubulin) antibody as a loading control (left). **B,** Western blotting analysis of ANPEP control, ROCK1 and MYPT1 knockdown mutants. Blots were probed with ROCK1 antibody (right) or α-tubulin antibody as a loading control (left). **C,** Western blotting analysis of ANPEP control cells and CFL1 knockdown mutants. Total protein samples were serially diluted. Blots were probed with CFL1 antibody (right) or α-Tubulin antibody as a loading control (left).

**Figure S6. Imaging of microtubules in Rho-pathway knockdown mutants.** Immunofluorescence was performed with an antibody recognizing α- and β-tubulin (yellow) to assess the microtubule cytoskeleton in ANPEP control and Rho-pathway knockdown mutants. Cells were also stained with DAPI to label DNA (blue). Images are representative of >50 cells imaged for each cell line. Scale bar (yellow), 10 μm.

**Figure S7. Analysis of amoebic attachment to Rho-pathway knockdown mutants.** Amoebae were stained with CMFDA and Jurkat cells (ANPEP control or Rho-pathway knockdown mutants) were stained with DiD. Cells were co-incubated for 5, 20, 40 or 60 minutes, and amoebic trogocytosis was quantified using imaging flow cytometry (see Figure 6 and Supplementary Figure 1). Images that contained single amoebae were then gated by maximum pixel intensity for channel 11, corresponding to DiD dye, to identify images that contained Jurkat cell material (Jurkat+).

**Figure S8. Normalized analysis of amoebic trogocytosis of Rho-pathway knockdown mutants.**

The trogocytosis assay data shown in Figure 6 were normalized such that amoebic ingestion of ANPEP control human cells was set to 100%. Amoebae were stained with CMFDA and Jurkat cells (ANPEP control or Rho-pathway knockdown mutants) were stained with DiD. Cells were co-incubated for 5, 20, 40 or 60 minutes, and amoebic trogocytosis was quantified using imaging flow cytometry (see Figure S1). **A,** The percentage of amoebae that performed ingestion was quantified (as % Internalization+), and amoebic ingestion of ANPEP control cells was set to 100%. **B,** Among amoebae that had performed ingestion, the proportion of amoebae that contained a low level of bites was quantified (% Low Trogocytosis), and low trogocytosis of ANPEP control cells was set to 100%. **C,** Among amoebae that had performed ingestion, the proportion of amoebae that contained a high level of bites was quantified (% High Trogocytosis) and high trogocytosis of ANPEP control cells was set to 100. **D-F**, Data shown are the fold change of amoebic ingestion of Rho-pathway knockdown mutants relative to amoebic ingestion of ANPEP control cells, which is set to 1 and represented by a dashed line. Data represent two to three separate biological replicates with two technical replicates per condition, for a total of four to six replicates per condition. Statistical analyses were performed using ordinary 2-way ANOVA with Dunnett’s multiple comparisons test.

**Figure S9. AFM technology and fitting of Young’s moduli.**

**A,** Schematic diagram of AFM-based measurements of single-cell mechanics**. B-D**, Representative line fitting process using a single compression of a CFL1 knockdown mutant. **B,** Representative Force *vs.* Relative Deformation curve. Red box represents area selected for line fitting. **C**, Nonlinear least-square fitting of the profile in (B) using Eq. (1), = 0.00 – 0.30; = 3.83 MPa (*x*^2^.69 * 10 ) (red). **D**, Nonlinear least-square fitting of the same cell using Eq. (2), = 0.00 – 0.45; = 1.68 kPa (*x*^2^ 6.73 * 10 ) (red).

**Figure S10. Cell** height and fold change comparisons of control *vs.* mutant single-cell compressions.

Individual AFM data comparisons of Control vs. Rho-pathway knockdown mutant single-cell compressions. **A**, Cell height distribution for all cell lines. Rac2 knockdown mutants are significantly shorter than control. Statistical analyses were performed using ordinary one-way ANOVA with Dunnett’s multiple comparisons tests. **B,** Fold change (Log_2_) of height of the cell, μm. **C**, Fold change (Log_2_) of Young’s Modulus of the cell membrane, . **D**, Fold change (Log_2_) of Young’s Modulus of the cytoskeleton, .

**Figure S11. AFM analysis of control cells.**

9 single cell AFM compressions for control Jurkat cells were acquired. Line fittings for each individual cell are labeled horizontally: **A,** Force *vs.* relative deformation profile of compressed control cells. **B,** Nonlinear least-square fitting of the profile in A using Eq. (1), varies. **C,** Nonlinear least-square fitting of the profile in A using eq. (2), varies.

**Figure S12. AFM analysis of CFL1 knockdown mutants.**

8 single cell AFM compressions for CFL1 knockdown mutants were acquired. Line fittings for each individual cell are labeled horizontally: **A,** Force *vs.* relative deformation profile of compressed control cells. **B,** Nonlinear least-square fitting of the profile in A using Eq. (1), varies. **C,** Nonlinear least-square fitting of the profile in A using eq. (2), varies. The 4^th^ cell compression/line fitting is presented in the main text.

**Figure S13. AFM analysis of MYPT1 knockdown mutants.**

5 single cell AFM compressions for MYPT1 knockdown mutants were acquired. Line fittings for each individual cell are labeled horizontally: **A,** Force *vs.* relative deformation profile of compressed control cells. **B,** Nonlinear least-square fitting of the profile in A using Eq. (1), varies. **C,** Nonlinear least-square fitting of the profile in A using eq. (2), varies.

**Figure S14. AFM analysis of Rac1 knockdown mutants.**

9 single cell AFM compressions for Rac1 knockdown mutants were acquired. Line fittings for each individual cell are labeled horizontally: **A,** Force *vs.* relative deformation profile of compressed control cells. **B,** Nonlinear least-square fitting of the profile in A using Eq. (1), varies. **C,** Nonlinear least-square fitting of the profile in A using eq. (2), varies.

**Figure S15. AFM analysis of Rac2 knockdown mutants.**

13 single cell AFM compressions for Rac2 KD knockdown mutants were acquired. Line fittings for each individual cell are labeled horizontally: **A,** Force *vs.* relative deformation profile of compressed control cells. **B,** Nonlinear least-square fitting of the profile in A using Eq. (1), varies. **C,** Nonlinear least-square fitting of the profile in A using eq. (2), varies.

**Figure S16. AFM analysis of RhoA knockdown mutants.**

9 single cell AFM compressions for RhoA KD knockdown mutants were acquired. Line fittings for each individual cell are labeled horizontally: **A,** Force *vs.* relative deformation profile of compressed control cells. **B,** Nonlinear least-square fitting of the profile in A using Eq. (1), varies. **C,** Nonlinear least-square fitting of the profile in A using eq. (2), varies.

**Figure S17. AFM analysis of ROCK1 knockdown mutants.**

10 single cell AFM compressions for ROCK1 knockdown mutants were acquired. Line fittings for each individual cell are labeled horizontally: **A,** Force *vs.* relative deformation profile of compressed control cells. **B,** Nonlinear least-square fitting of the profile in A using Eq. (1), varies. **C,** Nonlinear least-square fitting of the profile in A using eq. (2), varies.

**Figure S18. AFM analysis of ROCK2 knockdown mutants.**

7 single cell AFM compressions for ROCK2 knockdown mutants were acquired. Line fittings for each individual cell are labeled horizontally: **A,** Force *vs.* relative deformation profile of compressed control cells. **B,** Nonlinear least-square fitting of the profile in A using Eq. (1), varies. **C,** Nonlinear least-square fitting of the profile in A using eq. (2), varies.

## Notes

### Competing Interest Statement

The authors have declared no competing interest.

### Summary of Updates

In this revision, we added several new findings that strengthen the mechanistic, molecular-level explanation linking altered F-actin organization in Jurkat cell knockdown mutants to changes in amoebic trogocytosis. In new Figures 2 and 3, we used pharmacological inhibition of F-actin in Jurkat cells. This impacted amoebic trogocytosis in the same manner as knockdown mutants, thus corroborating our findings using knockdown mutants. To strengthen the connection between altered cytoskeletal rigidity of knockdown mutants and the alterations in amoebic trogocytosis, we directly measured the stiffness of the knockdown mutants by using atomic force microscopy (new Figure 8, new Table 1, new Supplementary Figure S9-S18). We provide new analysis of the microtubule cytoskeleton in knockdown mutants (new Figure S6), showing that there are no major differences between these mutants. In addition to these key new figures, we also provided additional new analysis (new Supplementary Figure S2, S3, S7, S8) and a new figure to summarize the overall model (new Figure 9).

## References

1. Shirley DAT, Farr L, Watanabe K, Moonah S. A Review of the Global Burden, New Diagnostics, and Current Therapeutics for Amebiasis. Open Forum Infect Dis. 2018 Jul;5(7):ofy161. doi:10.1093/ofid/ofy161 PubMed PMID: 30046644; PubMed Central PMCID: PMC6055529.

2. Haque R, Huston CD, Hughes M, Houpt E, Petri WA. Amebiasis. N Engl J Med. 2003 Apr 17;348(16):1565–73. doi:10.1056/NEJMra022710 PubMed PMID: 12700377.

3. Shrivastav MT, Malik Z, Somlata null. Revisiting Drug Development Against the Neglected Tropical Disease, Amebiasis. Front Cell Infect Microbiol. 2020;10:628257. doi:10.3389/fcimb.2020.628257 PubMed PMID: 33718258; PubMed Central PMCID: PMC7943716.

4. Gonzales MLM, Dans LF, Sio-Aguilar J. Antiamoebic drugs for treating amoebic colitis. Cochrane Database Syst Rev. 2019 Jan 9;1(1):CD006085. doi:10.1002/14651858.CD006085.pub3 PubMed PMID: 30624763; PubMed Central PMCID: PMC6326239.

5. Kantor M, Abrantes A, Estevez A, Schiller A, Torrent J, Gascon J, et al. Entamoeba Histolytica: Updates in Clinical Manifestation, Pathogenesis, and Vaccine Development. Can J Gastroenterol Hepatol. 2018 Dec 2;2018:4601420. doi:10.1155/2018/4601420 PubMed PMID: 30631758; PubMed Central PMCID: PMC6304615.

6. Stanley SL. Amoebiasis. Lancet Lond Engl. 2003 Mar 22;361(9362):1025–34. doi:10.1016/S0140-6736(03)12830-9 PubMed PMID: 12660071.

7. Chou A, Austin RL. Entamoeba histolytica Infection. In: StatPearls [Internet]. Treasure Island (FL): StatPearls Publishing; 2023 [cited 2023 Jul 27]. Available from: http://www.ncbi.nlm.nih.gov/books/NBK557718/ PubMed PMID: 32491650.

8. Marie C, Petri WA. Amoebic dysentery. BMJ Clin Evid. 2013 Aug 30;2013:0918. PubMed PMID: 23991750; PubMed Central PMCID: PMC3758071.

9. Stauffer W, Ravdin JI. Entamoeba histolytica: an update. Curr Opin Infect Dis. 2003 Oct;16(5):479–85. doi:10.1097/00001432-200310000-00016 PubMed PMID: 14502002.

10. Wang H, Naghavi M, Allen C, Barber RM, Bhutta ZA, Carter A, et al. Global, regional, and national life expectancy, all-cause mortality, and cause-specific mortality for 249 causes of death, 1980–2015: a systematic analysis for the Global Burden of Disease Study 2015. The Lancet. 2016 Oct 8;388(10053):1459–544. doi:10.1016/S0140-6736(16)31012-1

11. Gilchrist CA, Petri SE, Schneider BN, Reichman DJ, Jiang N, Begum S, et al. Role of the Gut Microbiota of Children in Diarrhea Due to the Protozoan Parasite Entamoeba histolytica. J Infect Dis. 2016 May 15;213(10):1579–85. doi:10.1093/infdis/jiv772 PubMed PMID: 26712950; PubMed Central PMCID: PMC4837909.

12. Mondal D, Petri WA Jr, Sack RB, Kirkpatrick BD, Haque R. Entamoeba histolytica-associated diarrheal illness is negatively associated with the growth of preschool children: evidence from a prospective study. Trans R Soc Trop Med Hyg. 2006 Nov 1;100(11):1032–8. doi:10.1016/j.trstmh.2005.12.012

13. Ralston KS, Solga MD, Mackey-Lawrence NM, Somlata, Bhattacharya A, Petri WA. Trogocytosis by Entamoeba histolytica contributes to cell killing and tissue invasion. Nature. 2014 Apr;508(7497):7497. doi:10.1038/nature13242

14. Bracha R, Nuchamowitz Y, Mirelman D. Transcriptional Silencing of an Amoebapore Gene in Entamoeba histolytica: Molecular Analysis and Effect on Pathogenicity. Eukaryot Cell. 2003 Apr;2(2):295–305. doi:10.1128/EC.2.2.295-305.2003 PubMed PMID: 12684379; PubMed Central PMCID: PMC154849.

15. Bracha R, Nuchamowitz Y, Leippe M, Mirelman D. Antisense inhibition of amoebapore expression in Entamoeba histolytica causes a decrease in amoebic virulence. Mol Microbiol. 1999 Nov;34(3):463–72. doi:10.1046/j.1365-2958.1999.01607.x PubMed PMID: 10564488.

16. Leippe M. Amoebapores. Parasitol Today. 1997 May 1;13(5):178–83. doi:10.1016/S0169-4758(97)01038-7

17. Leippe M, Andrä J, Müller-Eberhard HJ. Cytolytic and antibacterial activity of synthetic peptides derived from amoebapore, the pore-forming peptide of Entamoeba histolytica. Proc Natl Acad Sci U S A. 1994 Mar 29;91(7):2602–6. doi:10.1073/pnas.91.7.2602 PubMed PMID: 8146160; PubMed Central PMCID: PMC43417.

18. Leippe M, Andrä J, Nickel R, Tannich E, Müller-Eberhard HJ. Amoebapores, a family of membranolytic peptides from cytoplasmic granules of Entamoeba histolytica: isolation, primary structure, and pore formation in bacterial cytoplasmic membranes. Mol Microbiol. 1994 Dec;14(5):895–904. doi:10.1111/j.1365-2958.1994.tb01325.x PubMed PMID: 7715451.

19. Hecht O, van Nuland NA, Schleinkofer K, Dingley AJ, Bruhn H, Leippe M, et al. Solution Structure of the Pore-forming Protein of *Entamoeba histolytica*\*. J Biol Chem. 2004 Apr 23;279(17):17834–41. doi:10.1074/jbc.M312978200

20. Evans DF, Pye G, Bramley R, Clark AG, Dyson TJ, Hardcastle JD. Measurement of gastrointestinal pH profiles in normal ambulant human subjects. Gut. 1988 Aug;29(8):1035–41. doi:10.1136/gut.29.8.1035 PubMed PMID: 3410329; PubMed Central PMCID: PMC1433896.

21. Ralston KS, Petri WA. Tissue destruction and invasion by Entamoeba histolytica. Trends Parasitol. 2011 Jun 1;27(6):254–63. doi:10.1016/j.pt.2011.02.006

22. Leippe M. Ancient weapons: NK-lysin, is a mammalian homolog to pore-forming peptides of a protozoan parasite. Cell. 1995 Oct 6;83(1):17–8. doi:10.1016/0092-8674(95)90229-5 PubMed PMID: 7553868.

23. Miyake K, Karasuyama H. The Role of Trogocytosis in the Modulation of Immune Cell Functions. Cells. 2021 May;10(5):5. doi:10.3390/cells10051255

24. Nakada-Tsukui K, Nozaki T. Trogocytosis in Unicellular Eukaryotes. Cells. 2021 Nov;10(11):11. doi:10.3390/cells10112975

25. Nakayama M, Hori A, Toyoura S, Yamaguchi SI. Shaping of T Cell Functions by Trogocytosis. Cells. 2021 May;10(5):5. doi:10.3390/cells10051155

26. Bettadapur A, Miller HW, Ralston KS. Biting Off What Can Be Chewed: Trogocytosis in Health, Infection, and Disease. Infect Immun. 2020 Jun 22;88(7):e00930–19. doi:10.1128/IAI.00930-19

27. Batista FD, Iber D, Neuberger MS. B cells acquire antigen from target cells after synapse formation. Nature. 2001 May;411(6836):6836. doi:10.1038/35078099

28. Brown T. Observations by immunofluorescence microscopy and electron microscopy on the cytopathogenicity of Naegleria fowleri in mouse embryo-cell cultures. J Med Microbiol. 1979 Aug;12(3):363–71. doi:10.1099/00222615-12-3-363 PubMed PMID: 381667.

29. Waddell DR, Vogel G. Phagocytic behavior of the predatory slime mold, Dictyostelium caveatum. Cell nibbling. Exp Cell Res. 1985 Aug;159(2):323–34. doi:10.1016/s0014-4827(85)80006-9 PubMed PMID: 4029272.

30. Culbertson CG. The pathogenicity of soil amebas. Annu Rev Microbiol. 1971;25:231–54. doi:10.1146/annurev.mi.25.100171.001311 PubMed PMID: 5005026.

31. Culbertson CG. Pathogenic Naegleria and Hartmannella (Acenthamoeba). Ann N Y Acad Sci. 1970 Oct 30;174(2):1018–22. doi:10.1111/j.1749-6632.1970.tb45623.x PubMed PMID: 5278125.

32. Matlung HL, Babes L, Zhao XW, Houdt M van, Treffers LW, Rees DJ van, et al. Neutrophils Kill Antibody-Opsonized Cancer Cells by Trogoptosis. Cell Rep. 2018 Jun 26;23(13):3946–3959.e6. doi:10.1016/j.celrep.2018.05.082 PubMed PMID: 29949776.

33. Velmurugan R, Challa DK, Ram S, Ober RJ, Ward ES. Macrophage-Mediated Trogocytosis Leads to Death of Antibody-Opsonized Tumor Cells. Mol Cancer Ther. 2016 Aug;15(8):1879–89. doi:10.1158/1535-7163.MCT-15-0335 PubMed PMID: 27226489; PubMed Central PMCID: PMC4975628.

34. Kim J, Park S, Kim J, Kim Y, Yoon HM, Rayhan BR, et al. Trogocytosis-mediated immune evasion in the tumor microenvironment. Exp Mol Med. 2025 Jan;57(1):1–12. doi:10.1038/s12276-024-01364-2

35. Chen Y, Xin Q, Zhu M, Qiu J, Qiu J, Li R, et al. Trogocytosis in CAR immune cell therapy: a key mechanism of tumor immune escape. Cell Commun Signal. 2024 Oct 28;22(1):521. doi:10.1186/s12964-024-01894-2

36. Lim TK, Ruthazer ES. Microglial trogocytosis and the complement system regulate axonal pruning in vivo. eLife. 2021 Mar 16;10:e62167. doi:10.7554/eLife.62167 PubMed PMID: 33724186; PubMed Central PMCID: PMC7963485.

37. Weinhard L, di Bartolomei G, Bolasco G, Machado P, Schieber NL, Neniskyte U, et al. Microglia remodel synapses by presynaptic trogocytosis and spine head filopodia induction. Nat Commun. 2018 Mar 26;9(1):1228. doi:10.1038/s41467-018-03566-5 PubMed PMID: 29581545; PubMed Central PMCID: PMC5964317.

38. Abdu Y, Maniscalco C, Heddleston JM, Chew TL, Nance J. Developmentally programmed germ cell remodelling by endodermal cell cannibalism. Nat Cell Biol. 2016 Dec;18(12):1302–10. doi:10.1038/ncb3439 PubMed PMID: 27842058; PubMed Central PMCID: PMC5129868.

39. Saito-Nakano Y, Wahyuni R, Nakada-Tsukui K, Tomii K, Nozaki T. Rab7D small GTPase is involved in phago-, trogocytosis and cytoskeletal reorganization in the enteric protozoan Entamoeba histolytica. Cell Microbiol. 2021 Jan;23(1):e13267. doi:10.1111/cmi.13267 PubMed PMID: 32975360; PubMed Central PMCID: PMC7757265.

40. Gilmartin AA, Ralston KS, Petri WA. Inhibition of Amebic Cysteine Proteases Blocks Amebic Trogocytosis but Not Phagocytosis. J Infect Dis. 2020 May 15;221(10):1734–9. doi:10.1093/infdis/jiz671 PubMed PMID: 31999350; PubMed Central PMCID: PMC7184912.

41. Morrissey MA, Williamson AP, Steinbach AM, Roberts EW, Kern N, Headley MB, et al. Chimeric antigen receptors that trigger phagocytosis. eLife. 2018 Jun 4;7:e36688. doi:10.7554/eLife.36688 PubMed PMID: 29862966; PubMed Central PMCID: PMC6008046.

42. Gilmartin AA, Ralston KS, Petri WA. Inhibition of Amebic Lysosomal Acidification Blocks Amebic Trogocytosis and Cell Killing. mBio. 2017 Aug 29;8(4):e01187–17. doi:10.1128/mBio.01187-17 PubMed PMID: 28851845; PubMed Central PMCID: PMC5574710.

43. Martínez-Martín N, Fernández-Arenas E, Cemerski S, Delgado P, Turner M, Heuser J, et al. T cell receptor internalization from the immunological synapse is mediated by TC21 and RhoG GTPase-dependent phagocytosis. Immunity. 2011 Aug 26;35(2):208–22. doi:10.1016/j.immuni.2011.06.003 PubMed PMID: 21820331; PubMed Central PMCID: PMC4033310.

44. Gong J, Gaitanos TN, Luu O, Huang Y, Gaitanos L, Lindner J, et al. Gulp1 controls Eph/ephrin trogocytosis and is important for cell rearrangements during development. J Cell Biol. 2019 Oct 7;218(10):3455–71. doi:10.1083/jcb.201901032 PubMed PMID: 31409653; PubMed Central PMCID: PMC6781437.

45. Wetzel SA, McKeithan TW, Parker DC. Peptide-specific intercellular transfer of MHC class II to CD4+ T cells directly from the immunological synapse upon cellular dissociation. J Immunol Baltim Md 1950. 2005 Jan 1;174(1):80–9. doi:10.4049/jimmunol.174.1.80 PubMed PMID: 15611230.

46. Vanherberghen B, Andersson K, Carlin LM, Nolte-’t Hoen ENM, Williams GS, Höglund P, et al. Human and murine inhibitory natural killer cell receptors transfer from natural killer cells to target cells. Proc Natl Acad Sci U S A. 2004 Nov 30;101(48):16873–8. doi:10.1073/pnas.0406240101 PubMed PMID: 15550544; PubMed Central PMCID: PMC534731.

47. Hudrisier D, Riond J, Mazarguil H, Gairin JE, Joly E. Cutting edge: CTLs rapidly capture membrane fragments from target cells in a TCR signaling-dependent manner. J Immunol Baltim Md 1950. 2001 Mar 15;166(6):3645–9. doi:10.4049/jimmunol.166.6.3645 PubMed PMID: 11238601.

48. Becker SM, Cho KN, Guo X, Fendig K, Oosman MN, Whitehead R, et al. Epithelial Cell Apoptosis Facilitates Entamoeba histolytica Infection in the Gut. Am J Pathol. 2010 Mar 1;176(3):1316–22. doi:10.2353/ajpath.2010.090740

49. Ravdin JI, Moreau F, Sullivan JA, Petri WA, Mandell GL. Relationship of free intracellular calcium to the cytolytic activity of Entamoeba histolytica. Infect Immun. 1988 Jun;56(6):1505–12. doi:10.1128/iai.56.6.1505-1512.1988 PubMed PMID: 2897335; PubMed Central PMCID: PMC259428.

50. Petri WA, Smith RD, Schlesinger PH, Murphy CF, Ravdin JI. Isolation of the galactose-binding lectin that mediates the in vitro adherence of Entamoeba histolytica. J Clin Invest. 1987 Nov;80(5):1238–44. doi:10.1172/JCI113198 PubMed PMID: 2890654; PubMed Central PMCID: PMC442376.

51. Trissl D, Martínez-Palomo A, de la Torre M, de la Hoz R, Pérez de Suárez E. Surface properties of Entamoeba: increased rates of human erythrocyte phagocytosis in pathogenic strains. J Exp Med. 1978 Nov 1;148(5):1137–43. doi:10.1084/jem.148.5.1137 PubMed PMID: 722237; PubMed Central PMCID: PMC2185040.

52. Rodríguez MA, Orozco E. Isolation and characterization of phagocytosis- and virulence-deficient mutants of Entamoeba histolytica. J Infect Dis. 1986 Jul 1;154(1):27–32. doi:10.1093/infdis/154.1.27 PubMed PMID: 2872254.

53. Orozco E, Suárez ME, Sánchez T. Differences in adhesion, phagocytosis and virulence of clones from Entamoeba histolytica, strain HM1:IMSS. Int J Parasitol. 1985 Dec;15(6):655–60. doi:10.1016/0020-7519(85)90012-8 PubMed PMID: 2869003.

54. E O, G G, A MP, T S. Entamoeba histolytica. Phagocytosis as a virulence factor. J Exp Med. 1983 Nov 1;158(5). doi:10.1084/jem.158.5.1511 PubMed PMID: 6313842.

55. Lejeune A, Gicquaud C. Target cell deformability determines the type of phagocytic mechanism used by Entamoeba histolytica-like, Laredo strain. Biol Cell. 1992 Jan 1;74:211–6. doi:10.1016/0248-4900(92)90027-X

56. Cai X, Xing X, Cai J, Chen Q, Wu S, Huang F. Connection between biomechanics and cytoskeleton structure of lymphocyte and Jurkat cells: An AFM study. Micron Oxf Engl 1993. 2010 Apr;41(3):257–62. doi:10.1016/j.micron.2009.08.011 PubMed PMID: 20060729.

57. Fletcher DA, Mullins RD. Cell mechanics and the cytoskeleton. Nature. 2010 Jan;463(7280):7280. doi:10.1038/nature08908

58. Olson EN, Nordheim A. Linking actin dynamics and gene transcription to drive cellular motile functions. Nat Rev Mol Cell Biol. 2010 May;11(5):5. doi:10.1038/nrm2890

59. Pollard TD, Borisy GG. Cellular Motility Driven by Assembly and Disassembly of Actin Filaments. Cell. 2003 Feb 21;112(4):453–65. doi:10.1016/S0092-8674(03)00120-X PubMed PMID: 12600310.

60. Akenuwa OH, Gu J, Nebenführ A, Abel SM. Morphometric analysis of actin networks. Mol Biol Cell. 2024 Dec;35(12):ar146. doi:10.1091/mbc.E24-06-0248

61. Sun J, Zhong X, Fu X, Miller H, Lee P, Yu B, et al. The Actin Regulators Involved in the Function and Related Diseases of Lymphocytes. Front Immunol. 2022 Mar 16;13:799309. doi:10.3389/fimmu.2022.799309 PubMed PMID: 35371070; PubMed Central PMCID: PMC8965893.

62. Fallqvist B, Fielden ML, Pettersson T, Nordgren N, Kroon M, Gad AKB. Experimental and computational assessment of F-actin influence in regulating cellular stiffness and relaxation behaviour of fibroblasts. J Mech Behav Biomed Mater. 2016 Jun 1;59:168–84. doi:10.1016/j.jmbbm.2015.11.039

63. Tapon N, Hall A. Rho, Rac and Cdc42 GTPases regulate the organization of the actin cytoskeleton. Curr Opin Cell Biol. 1997 Feb;9(1):86–92. doi:10.1016/s0955-0674(97)80156-1 PubMed PMID: 9013670.

64. Chauhan BK, Lou M, Zheng Y, Lang RA. Balanced Rac1 and RhoA activities regulate cell shape and drive invagination morphogenesis in epithelia. Proc Natl Acad Sci. 2011 Nov 8;108(45):18289–94. doi:10.1073/pnas.1108993108

65. Abreu-Blanco MT, Verboon JM, Parkhurst SM. Coordination of Rho Family GTPase Activities to Orchestrate Cytoskeleton Responses during Cell Wound Repair. Curr Biol. 2014 Jan 20;24(2):144–55. doi:10.1016/j.cub.2013.11.048

66. Ridley AJ. Rho GTPases and actin dynamics in membrane protrusions and vesicle trafficking. Trends Cell Biol. 2006 Oct;16(10):522–9. doi:10.1016/j.tcb.2006.08.006

67. Tsai CH, Chang CY, Lin BZ, Wu YL, Wu MH, Lin LT, et al. Up-regulation of cofilin-1 in cell senescence associates with morphological change and p27kip1-mediated growth delay. Aging Cell. 2021;20(1):e13288. doi:10.1111/acel.13288

68. Kunschmann T, Puder S, Fischer T, Steffen A, Rottner K, Mierke CT. The Small GTPase Rac1 Increases Cell Surface Stiffness and Enhances 3D Migration Into Extracellular Matrices. Sci Rep. 2019 May 22;9(1):1. doi:10.1038/s41598-019-43975-0

69. Wiggan O, Schroder B, Krapf D, Bamburg JR, DeLuca JG. Cofilin Regulates Nuclear Architecture through a Myosin-II Dependent Mechanotransduction Module. Sci Rep. 2017 Jan 19;7(1):1. doi:10.1038/srep40953

70. Galkin VE, Orlova A, Kudryashov DS, Solodukhin A, Reisler E, Schröder GF, et al. Remodeling of actin filaments by ADF/cofilin proteins. Proc Natl Acad Sci U S A. 2011 Dec 20;108(51):20568–72. doi:10.1073/pnas.1110109108 PubMed PMID: 22158895; PubMed Central PMCID: PMC3251117.

71. Sun CX, Magalhães MAO, Glogauer M. Rac1 and Rac2 differentially regulate actin free barbed end formation downstream of the fMLP receptor. J Cell Biol. 2007 Oct 22;179(2):239–45. doi:10.1083/jcb.200705122 PubMed PMID: 17954607; PubMed Central PMCID: PMC2064760.

72. Ghosh M, Song X, Mouneimne G, Sidani M, Lawrence DS, Condeelis JS. Cofilin Promotes Actin Polymerization and Defines the Direction of Cell Motility. Science. 2004 Apr 30;304(5671):743–6. doi:10.1126/science.1094561

73. Tixeira R, Phan TK, Caruso S, Shi B, Atkin-Smith GK, Nedeva C, et al. ROCK1 but not LIMK1 or PAK2 is a key regulator of apoptotic membrane blebbing and cell disassembly. Cell Death Differ. 2020 Jan;27(1):1. doi:10.1038/s41418-019-0342-5

74. Bros M, Haas K, Moll L, Grabbe S. RhoA as a Key Regulator of Innate and Adaptive Immunity. Cells. 2019 Jul 17;8(7):733. doi:10.3390/cells8070733 PubMed PMID: 31319592; PubMed Central PMCID: PMC6678964.

75. Schiffhauer ES, Ren Y, Iglesias VA, Kothari P, Iglesias PA, Robinson DN. Myosin IIB assembly state determines its mechanosensitive dynamics. J Cell Biol. 2019 Mar 4;218(3):895–908. doi:10.1083/jcb.201806058 PubMed PMID: 30655296; PubMed Central PMCID: PMC6400566.

76. Huse M. Mechanical forces in the immune system. Nat Rev Immunol. 2017 Nov;17(11):11. doi:10.1038/nri.2017.74

77. Shi J, Wu X, Surma M, Vemula S, Zhang L, Yang Y, et al. Distinct roles for ROCK1 and ROCK2 in the regulation of cell detachment. Cell Death Dis. 2013 Feb;4(2):2. doi:10.1038/cddis.2013.10

78. Koga Y, Ikebe M. A Novel Regulatory Mechanism of Myosin Light Chain Phosphorylation via Binding of 14-3-3 to Myosin Phosphatase. Mol Biol Cell. 2008 Mar;19(3):1062–71. doi:10.1091/mbc.e07-07-0668

79. Maddox AS, Burridge K. RhoA is required for cortical retraction and rigidity during mitotic cell rounding. J Cell Biol. 2003 Jan 20;160(2):255–65. doi:10.1083/jcb.200207130 PubMed PMID: 12538643; PubMed Central PMCID: PMC2172639.

80. Sosale NG, Rouhiparkouhi T, Bradshaw AM, Dimova R, Lipowsky R, Discher DE. Cell rigidity and shape override CD47’s “self”-signaling in phagocytosis by hyperactivating myosin-II. Blood. 2015 Jan 15;125(3):542–52. doi:10.1182/blood-2014-06-585299 PubMed PMID: 25411427; PubMed Central PMCID: PMC4296014.

81. Beningo KA, Wang Y li. Fc-receptor-mediated phagocytosis is regulated by mechanical properties of the target. J Cell Sci. 2002 Feb 15;115(4):849–56. doi:10.1242/jcs.115.4.849

82. Vorselen D, Wang Y, de Jesus MM, Shah PK, Footer MJ, Huse M, et al. Microparticle traction force microscopy reveals subcellular force exertion patterns in immune cell–target interactions. Nat Commun. 2020 Jan 7;11:20. doi:10.1038/s41467-019-13804-z PubMed PMID: 31911639; PubMed Central PMCID: PMC6946705.

83. Settle AH, Winer BY, Jesus MM de, Seeman L, Wang Z, Chan E, et al. β2 integrins impose a mechanical checkpoint on macrophage phagocytosis [Internet]. bioRxiv; 2024 [cited 2024 Sep 10]. p. 2024.02.20.580845. Available from: https://www.biorxiv.org/content/10.1101/2024.02.20.580845v1 doi:10.1101/2024.02.20.580845

84. Sun Q, Luo T, Ren Y, Florey O, Shirasawa S, Sasazuki T, et al. Competition between human cells by entosis. Cell Res. 2014 Nov;24(11):11. doi:10.1038/cr.2014.138

85. Cornell CE, Chorlay A, Krishnamurthy D, Martin NR, Baldauf L, Fletcher DA. Target cell cortical tension regulates macrophage trogocytosis. Nat Cell Biol. 2025 Dec;27(12):2078–88. doi:10.1038/s41556-025-01807-6

86. Rollins KR, Fiaz S, Datta I, Morrissey MA. Target cell adhesion limits macrophage phagocytosis and promotes trogocytosis. J Cell Biol. 2025 Nov 3;224(11):e202502034. doi:10.1083/jcb.202502034 PubMed PMID: 41042177.

87. Mandal S, Melo M, Gordiichuk P, Acharya S, Poh YC, Li N, et al. WASP facilitates tumor mechanosensitivity in T lymphocytes. bioRxiv : the preprint server for biology. 2023. doi:10.1101/2023.10.02.560434

88. Leithner A, Altenburger LM, Hauschild R, Assen FP, Rottner K, Stradal TEB, et al. Dendritic cell actin dynamics control contact duration and priming efficiency at the immunological synapse. J Cell Biol. 2021 Feb 3;220(4):e202006081. doi:10.1083/jcb.202006081

89. Vaahtomeri K, Brown M, Hauschild R, De Vries I, Leithner AF, Mehling M, et al. Locally Triggered Release of the Chemokine CCL21 Promotes Dendritic Cell Transmigration across Lymphatic Endothelia. Cell Rep. 2017 May 2;19(5):902–9. doi:10.1016/j.celrep.2017.04.027 PubMed PMID: 28467903; PubMed Central PMCID: PMC5437727.

90. Saito S, Watabe S, Ozaki H, Fusetani N, Karaki H. Mycalolide B, a novel actin depolymerizing agent. J Biol Chem. 1994 Nov 25;269(47):29710–4. PubMed PMID: 7961961.

91. Hori M, Saito S, Shin YZ, Ozaki H, Fusetani N, Karaki H. Mycalolide-B, a novel and specific inhibitor of actomyosin ATPase isolated from marine sponge. FEBS Lett. 1993 May 10;322(2):151–4. doi:10.1016/0014-5793(93)81557-g PubMed PMID: 8482383.

92. Saffer LD, Petri WA. Role of the galactose lectin of Entamoeba histolytica in adherence-dependent killing of mammalian cells. Infect Immun. 1991 Dec;59(12):4681–3. doi:10.1128/iai.59.12.4681-4683.1991

93. Ravdin JI, Croft BY, Guerrant RL. Cytopathogenic mechanisms of Entamoeba histolytica. J Exp Med. 1980 Aug 1;152(2):377–90. doi:10.1084/jem.152.2.377 PubMed PMID: 6249882; PubMed Central PMCID: PMC2185944.

94. Larson MH, Gilbert LA, Wang X, Lim WA, Weissman JS, Qi LS. CRISPR interference (CRISPRi) for sequence-specific control of gene expression. Nat Protoc. 2013 Nov;8(11):2180–96. doi:10.1038/nprot.2013.132 PubMed PMID: 24136345; PubMed Central PMCID: PMC3922765.

95. Jinek M, Chylinski K, Fonfara I, Hauer M, Doudna JA, Charpentier E. A Programmable Dual-RNA–Guided DNA Endonuclease in Adaptive Bacterial Immunity. Science. 2012 Aug 17;337(6096):816–21. doi:10.1126/science.1225829

96. Murugesan S, Hong J, Yi J, Li D, Beach JR, Shao L, et al. Formin-generated actomyosin arcs propel T cell receptor microcluster movement at the immune synapse. J Cell Biol. 2016 Nov 7;215(3):383–99. doi:10.1083/jcb.201603080 PubMed PMID: 27799367; PubMed Central PMCID: PMC5100289.

97. Yi J, Wu XS, Crites T, Hammer JA. Actin retrograde flow and actomyosin II arc contraction drive receptor cluster dynamics at the immunological synapse in Jurkat T cells. Mol Biol Cell. 2012 Mar 1;23(5):834–52. doi:10.1091/mbc.E11-08-0731 PubMed PMID: 22219382; PubMed Central PMCID: PMC3290643.

98. Lulevich V, Zink T, Chen HY, Liu FT, Liu G yu. Cell Mechanics Using Atomic Force Microscopy-Based Single-Cell Compression. Langmuir. 2006 Sep 1;22(19):8151–5. doi:10.1021/la060561p

99. Krieg M, Fläschner G, Alsteens D, Gaub BM, Roos WH, Wuite GJL, et al. Atomic force microscopy-based mechanobiology. Nat Rev Phys. 2019 Jan;1(1):41–57. doi:10.1038/s42254-018-0001-7

100. Sneddon IN. The relation between load and penetration in the axisymmetric boussinesq problem for a punch of arbitrary profile. Int J Eng Sci. 1965 May 1;3(1):47–57. doi:10.1016/0020-7225(65)90019-4

101. Zhang Y, Shen H, Liu H, Feng H, Liu Y, Zhu X, et al. Arp2/3 complex controls T cell homeostasis by maintaining surface TCR levels via regulating TCR+ endosome trafficking. Sci Rep. 2017 Aug 21;7(1):8952. doi:10.1038/s41598-017-08357-4

102. Nolz JC, Gomez TS, Zhu P, Li S, Medeiros RB, Shimizu Y, et al. The WAVE2 Complex Regulates Actin Cytoskeletal Reorganization and CRAC-Mediated Calcium Entry during T Cell Activation. Curr Biol. 2006 Jan 10;16(1):24–34. doi:10.1016/j.cub.2005.11.036

103. Blanchoin L, Boujemaa-Paterski R, Sykes C, Plastino J. Actin Dynamics, Architecture, and Mechanics in Cell Motility. Physiol Rev. 2014 Jan;94(1):235–63. doi:10.1152/physrev.00018.2013

104. Yan C, Martinez-Quiles N, Eden S, Shibata T, Takeshima F, Shinkura R, et al. WAVE2 deficiency reveals distinct roles in embryogenesis and Rac_-_mediated actin_-_based motility. EMBO J. 2003 Jul 15;22(14):3602–12. doi:10.1093/emboj/cdg350

105. Worthylake RA, Burridge K. Leukocyte transendothelial migration: orchestrating the underlying molecular machinery. Curr Opin Cell Biol. 2001 Oct;13(5):569–77. doi:10.1016/s0955-0674(00)00253-2 PubMed PMID: 11544025.

106. Watanabe N, Kato T, Fujita A, Ishizaki T, Narumiya S. Cooperation between mDia1 and ROCK in Rho-induced actin reorganization. Nat Cell Biol. 1999 Jul;1(3):136–43. doi:10.1038/11056

107. Watanabe N, Madaule P, Reid T, Ishizaki T, Watanabe G, Kakizuka A, et al. p140mDia, a mammalian homolog of Drosophila diaphanous,is a target protein for Rho small GTPase and is a ligand for profilin. EMBO J. 1997 Jun;16(11):3044–56. doi:10.1093/emboj/16.11.3044

108. Heasman SJ, Ridley AJ. Multiple roles for RhoA during T cell transendothelial migration. Small GTPases. 2010;1(3):174–9. doi:10.4161/sgtp.1.3.14724 PubMed PMID: 21686273; PubMed Central PMCID: PMC3116607.

109. Lee MH, Kundu JK, Chae JI, Shim JH. Targeting ROCK/LIMK/cofilin signaling pathway in cancer. Arch Pharm Res. 2019 Jun 1;42(6):481–91. doi:10.1007/s12272-019-01153-w

110. Wan YJ, Yang Y, Leng QL, Lan B, Jia HY, Liu YH, et al. Vav1 increases Bcl-2 expression by selective activation of Rac2–Akt in leukemia T cells. Cell Signal. 2014 Oct 1;26(10):2202–9. doi:10.1016/j.cellsig.2014.05.015

111. Steffen A, Ladwein M, Dimchev GA, Hein A, Schwenkmezger L, Arens S, et al. Rac function is crucial for cell migration but is not required for spreading and focal adhesion formation. J Cell Sci. 2013 Oct 15;126(20):4572–88. doi:10.1242/jcs.118232

112. Arana E, Vehlow A, Harwood NE, Vigorito E, Henderson R, Turner M, et al. Activation of the small GTPase Rac2 via the B cell receptor regulates B cell adhesion and immunological-synapse formation. Immunity. 2008 Jan;28(1):88–99. doi:10.1016/j.immuni.2007.12.003 PubMed PMID: 18191593.

113. D’Souza-Schorey C, Boettner B, Van Aelst L. Rac Regulates Integrin-Mediated Spreading and Increased Adhesion of T Lymphocytes. Mol Cell Biol. 1998 Jul;18(7):3936–46. doi:10.1128/mcb.18.7.3936 PubMed PMID: 9632778; PubMed Central PMCID: PMC108978.

114. Didsbury J, Weber RF, Bokoch GM, Evans T, Snyderman R. rac, a novel ras-related family of proteins that are botulinum toxin substrates. J Biol Chem. 1989 Oct 5;264(28):16378–82. PubMed PMID: 2674130.

115. Jerrell RJ, Leih MJ, Parekh A. The ROCK isoforms differentially regulate the morphological characteristics of carcinoma cells. Small GTPases. 2017 Sep 18;11(2):131–7. doi:10.1080/21541248.2017.1341366 PubMed PMID: 28650698; PubMed Central PMCID: PMC7053931.

116. Jerrell RJ, Parekh A. Matrix rigidity differentially regulates invadopodia activity through ROCK1 and ROCK2. Biomaterials. 2016 Apr 1;84:119–29. doi:10.1016/j.biomaterials.2016.01.028

117. Lock FE, Ryan KR, Poulter NS, Parsons M, Hotchin NA. Differential Regulation of Adhesion Complex Turnover by ROCK1 and ROCK2. PLOS ONE. 2012 Feb 13;7(2):e31423. doi:10.1371/journal.pone.0031423

118. Amano M, Nakayama M, Kaibuchi K. Rho-kinase/ROCK: A key regulator of the cytoskeleton and cell polarity. Cytoskeleton. 2010;67(9):545–54. doi:10.1002/cm.20472

119. Lock FE, Hotchin NA. Distinct Roles for ROCK1 and ROCK2 in the Regulation of Keratinocyte Differentiation. PLOS ONE. 2009 Dec 4;4(12):e8190. doi:10.1371/journal.pone.0008190

120. Darenfed H, Dayanandan B, Zhang T, Hsieh SHK, Fournier AE, Mandato CA. Molecular characterization of the effects of Y-27632. Cell Motil Cytoskeleton. 2007 Feb;64(2):97–109. doi:10.1002/cm.20168 PubMed PMID: 17009325.

121. Seeland I, Xiong Y, Orlik C, Deibel D, Prokosch S, Küblbeck G, et al. The actin remodeling protein cofilin is crucial for thymic αβ but not γδ T-cell development. Bhandoola A, editor. PLOS Biol. 2018 Jul 9;16(7):e2005380. doi:10.1371/journal.pbio.2005380

122. Thauland TJ, Hu KH, Bruce MA, Butte MJ. Cytoskeletal adaptivity regulates T cell receptor signaling. Sci Signal. 2017 Mar 7;10(469):eaah3737. doi:10.1126/scisignal.aah3737 PubMed PMID: 28270556; PubMed Central PMCID: PMC5854469.

123. Samstag Y, John I, Wabnitz GH. Cofilin: a redox sensitive mediator of actin dynamics during T-cell activation and migration. Immunol Rev. 2013;256(1):30–47. doi:10.1111/imr.12115

124. Babich A, Li S, O’Connor RS, Milone MC, Freedman BD, Burkhardt JK. F-actin polymerization and retrograde flow drive sustained PLCγ1 signaling during T cell activation. J Cell Biol. 2012 Jun 4;197(6):775–87. doi:10.1083/jcb.201201018

125. Ishaq M, Lin BR, Bosche M, Zheng X, Yang J, Huang D, et al. LIM kinase 1 - dependent cofilin 1 pathway and actin dynamics mediate nuclear retinoid receptor function in T lymphocytes. BMC Mol Biol. 2011 Sep 16;12:41. doi:10.1186/1471-2199-12-41 PubMed PMID: 21923909; PubMed Central PMCID: PMC3187726.

126. Lee KH, Meuer SC, Samstag Y. Cofilin: a missing link between T cell co-stimulation and rearrangement of the actin cytoskeleton. Eur J Immunol. 2000 Mar;30(3):892–9. doi:10.1002/1521-4141(200003)30:3<892::AID-IMMU892>3.0.CO;2-U PubMed PMID: 10741406.

127. Dupré L, Boztug K, Pfajfer L. Actin Dynamics at the T Cell Synapse as Revealed by Immune-Related Actinopathies. Front Cell Dev Biol. 2021 Jun 24;9:665519. doi:10.3389/fcell.2021.665519 PubMed PMID: 34249918; PubMed Central PMCID: PMC8266300.

128. Fowell DJ, Kim M. The spatio-temporal control of effector T cell migration. Nat Rev Immunol. 2021 Sep;21(9):582–96. doi:10.1038/s41577-021-00507-0

129. Hong J, Murugesan S, Betzig E, Hammer JA. Contractile actomyosin arcs promote the activation of primary mouse T cells in a ligand-dependent manner. PLOS ONE. 2017 Aug 17;12(8):e0183174. doi:10.1371/journal.pone.0183174

130. Joo EE, Yamada KM. MYPT1 regulates contractility and microtubule acetylation to modulate integrin adhesions and matrix assembly. Nat Commun. 2014 Mar 25;5:3510. doi:10.1038/ncomms4510 PubMed PMID: 24667306; PubMed Central PMCID: PMC4190669.

131. Khasnis M, Nakatomi A, Gumpper K, Eto M. Reconstituted Human Myosin Light Chain Phosphatase Reveals Distinct Roles of Two Inhibitory Phosphorylation Sites of the Regulatory Subunit, MYPT1. Biochemistry. 2014 Apr 29;53(16):2701–9. doi:10.1021/bi5001728

132. Kumari S, Vardhana S, Cammer M, Curado S, Santos L, Sheetz M, et al. T Lymphocyte Myosin IIA is Required for Maturation of the Immunological Synapse. Front Immunol [Internet]. 2012 [cited 2023 Oct 6];3. Available from: https://www.frontiersin.org/articles/10.3389/fimmu.2012.00230

133. Miller HW, Tam TSY, Ralston KS. Entamoeba histolytica Develops Resistance to Complement Deposition and Lysis after Acquisition of Human Complement-Regulatory Proteins through Trogocytosis. mBio. 2022 Mar;13(2):e03163–21. doi:10.1128/mbio.03163-21

134. Miller HW, Suleiman RL, Ralston KS. Trogocytosis by Entamoeba histolytica Mediates Acquisition and Display of Human Cell Membrane Proteins and Evasion of Lysis by Human Serum. mBio. 2019 Apr 30;10(2):e00068–19. doi:10.1128/mBio.00068-19

135. Growth and Genetic Manipulation of Entamoeba histolytica [Internet]. doi:10.1002/cpz1.327

136. Diamond LS, Harlow DR, Cunnick CC. A new medium for the axenic cultivation of Entamoeba histolytica and other Entamoeba. Trans R Soc Trop Med Hyg. 1978;72(4):431–2. doi:10.1016/0035-9203(78)90144-x PubMed PMID: 212851.

137. Diamond LS. Techniques of axenic cultivation of Entamoeba histolytica Schaudinn, 1903 and E. histolytica-like amebae. J Parasitol. 1968 Oct;54(5):1047–56. PubMed PMID: 4319346.

138. Horlbeck MA, Gilbert LA, Villalta JE, Adamson B, Pak RA, Chen Y, et al. Compact and highly active next-generation libraries for CRISPR-mediated gene repression and activation. Adelman K, editor. eLife. 2016 Sep 23;5:e19760. doi:10.7554/eLife.19760

139. Levesque-Sergerie JP, Duquette M, Thibault C, Delbecchi L, Bissonnette N. Detection limits of several commercial reverse transcriptase enzymes: impact on the low- and high-abundance transcript levels assessed by quantitative RT-PCR. BMC Mol Biol. 2007 Oct 22;8:93. doi:10.1186/1471-2199-8-93 PubMed PMID: 17953766; PubMed Central PMCID: PMC2151766.

140. Ruijter JM, Ramakers C, Hoogaars WMH, Karlen Y, Bakker O, van den Hoff MJB, et al. Amplification efficiency: linking baseline and bias in the analysis of quantitative PCR data. Nucleic Acids Res. 2009 Apr;37(6):e45. doi:10.1093/nar/gkp045 PubMed PMID: 19237396; PubMed Central PMCID: PMC2665230.

141. Livak KJ, Schmittgen TD. Analysis of Relative Gene Expression Data Using Real-Time Quantitative PCR and the 2−ΔΔCT Method. Methods. 2001 Dec 1;25(4):402–8. doi:10.1006/meth.2001.1262

142. Rao X, Huang X, Zhou Z, Lin X. An improvement of the 2^(–delta delta CT) method for quantitative real-time polymerase chain reaction data analysis. Biostat Bioinforma Biomath. 2013 Aug;3(3):71–85. PubMed PMID: 25558171; PubMed Central PMCID: PMC4280562.

143. Morales Maldonado AD, Diloretto DA, Timofeyev V, Karsai A, Ogorodnik E, Huang Y, et al. Protocol for measuring membrane elasticity of mouse cardiomyocytes using atomic force microscopy. STAR Protoc. 2025 Apr 11;6(2):103746. doi:10.1016/j.xpro.2025.103746 PubMed PMID: 40222013; PubMed Central PMCID: PMC12018553.

144. Cleveland JP, Manne S, Bocek D, Hansma PK. A nondestructive method for determining the spring constant of cantilevers for scanning force microscopy. Rev Sci Instrum. 1993 Feb 1;64(2):403–5. doi:10.1063/1.1144209

145. Lulevich V, Shih YP, Lo SH, Liu G yu. Cell Tracing Dyes Significantly Change Single Cell Mechanics. J Phys Chem B. 2009 May 7;113(18):6511–9. doi:10.1021/jp8103358

