## Supplemental Information for "Human cell cytoskeletal rigidity differentially influences trogocytosis and phagocytosis by *Entamoeba histolytica*"

### Supplementary Information

#### Supplemental Methods

##### Immunofluorescence Assays

To visualize microtubules, Jurkat cells were prepared, fixed, and permeabilized as described for phalloidin staining. Blocking was then performed with 1% bovine serum albumin (BSA) (GeminiBio) and 0.3% Triton X-100 in 1X PBS for 2 hours at room temperature. Samples were incubated with a 1:500 dilution of a sheep polyclonal antibody reactive to α- and β-tubulin (Cytoskeleton Inc., #ATN02-S) overnight at 4°C. After washing, secondary antibody staining was performed in the dark for 1 hour at room temperature using a donkey anti-sheep polyclonal antibody conjugated to Alexa Fluor 647 (Abcam, #ab150179). Cells were subsequently washed and stained with 100 nM DAPI (Millipore Sigma) for 5 minutes at room temperature, washed again, and were mounted on glass slides as described for phalloidin staining. Images were collected and processed as described for phalloidin staining, with false recoloring of microtubules from 647 red to yellow.

#### Supplementary Table 1.

| Experimental Purpose | Name | Sequence (5' → 3') |
|---|---|---|
| PCR Amplification of Hygromycin from pEhEx_empty-vector_HygR | GHP 5F | CTCAAGCCTCAGACAGTGGTTCAAAGTTTTTTTCTTCCA TTTCAGGTGTCGTGAGATGAAAAAGCCTGAACTCAC |
| | GHP 5R | TGTATAACTTCGTATAATGTATGCTATACGAAGTTATTAG GTCCCTCGACGAATTCTATTCCTTTGCCCTCGGAC |
| Sanger sequencing of SC0217_pU6-sgRNA ANPEP#1-Hyg | SC0217 Hyg Seq1F | TTCCATTTCAGGTGTCGTGA |
| | SC0217 Hyg Seq1R | GACATATCCACGCCCTCCTA |
| | SC0217 Hyg Seq2F | GAATTCAGCGAGAGCCTGAC |
| | SC0217 Hyg Seq2R | AGCAATCGCGCATATGAAAT |
| | SC0217 Hyg Seq3F | TCACTGGCAAACTGTGATGG |
| | SC0217 Hyg Seq3R | GATGTTGGCGACCTCGTATT |
| | SC0217 Hyg Seq4F | ATACGAGGTCGCCAACATCT |
| | SC0217 Hyg Seq4R | ATTTGTGTACGCCCGACAGT |
| sgRNA Cloning with BstXI and BlpI Overhang Sequences | CFL1 Forward | TTGGGCCGGGACCCGACTGAACGGTTTAAGAGC |
| | CFL1 Reverse | TTAGCTCTTAAACCGTTCAGTCGGGTCCCGGCCCAACAAG |
| | MYPT1 Forward | TTGGATGAGTGCGGGCCAGAGGAGTTTAAGAGC |
| | MYPT1 Reverse | TTAGCTCTTAAACTCCTCTGGCCCGCACTCATCCAACAAG |
| | Rac1 Forward | TTGGCTACGAGCGCTCCCGAGGAGTTTAAGAGC |
| | Rac1 Reverse | TTAGCTCTTAAACTCCTCGGGAGCGCTCGTAGCCAACAAG |
| | Rac2 Forward | TTGGCTGAGGAGCAGCGGTGGTGGTTTAAGAGC |
| | Rac2 Reverse | TTAGCTCTTAAACCACCACCGCTGCTCCTCAGCCAACAAG |
| | RhoA Forward | TTGGGTAGCTGAAGACCAGACCGGTTTAAGAGC |
| | RhoA Reverse | TTAGCTCTTAAACCGGTCTGGTCTTCAGCTACCCAACAAG |
| | ROCK1 Forward | TTGGGACTCCCTCCGGGCAACAAGTTTAAGAGC |
| | ROCK1 Reverse | TTAGCTCTTAAACTTGTTGCCCGGAGGGAGTCCCAACAAG |
| | ROCK2 Forward | TTGGGCGCGCGGCCGGAGACGGGGTTTAAGAGC |
| | ROCK2 Reverse | TTAGCTCTTAAACCCCGTCTCCGGCCGCGCGCCCAACAAG |
| Sanger sequencing of sgRNAs (U6 Promoter) | U6_seq | GGTACAGTGCAGGGGAAAGA |
| qPCR Housekeeping Genes | GAPDH Forward | AATCCCATCACCATCTTCCA |
| | GAPDH Reverse | AAATGAGCCCCAGCCTTC |
| | RPLP0 Forward | ATCATCAACGGGTACAAACGAGTC |
| | RPLP0 Reverse | GCAGATGGATCAGCCAAGAAGG |
| qPCR Cytoskeletal Knockdown Genes of Interest | CFL1 Forward | CTGCAGCGCTCTCGTCTT |
| | CFL1 Reverse | ACACCGGAGGCCATGTTTC |
| | MYPT1 Forward | TTGGAAATGGAAAAAAGGGAACG |
| | MYPT1 Reverse | CCCATTTTCATCCTTTAGCCTCT |
| | Rac1 Forward | TCCGCAAACAGATGTGTTCTTA |
| | Rac1 Reverse | CGCACCTCAGGATACCACTTT |
| | Rac2 Forward | TGGCCAAGGAGATTGACTCG |
| | Rac2 Reverse | GCCTCGTCGAACACGGTTT |
| | RhoA Forward | TCGTTAGTCCACGGTCTGGT |
| | RhoA Reverse | CAGCCATTGCTCAGGCAAC |
| | ROCK1 Forward | GAGGCAAAGTCTGTGGCAAT |
| | ROCK1 Reverse | TCAGCCTTCTCTCGAGCTTC |
| | ROCK2 Forward | CAACTGTGAGGCTTGTATGAAG |
| | ROCK2 Reverse | TGCAAGGTGCTATAATCTCCTC |

### Figure S1

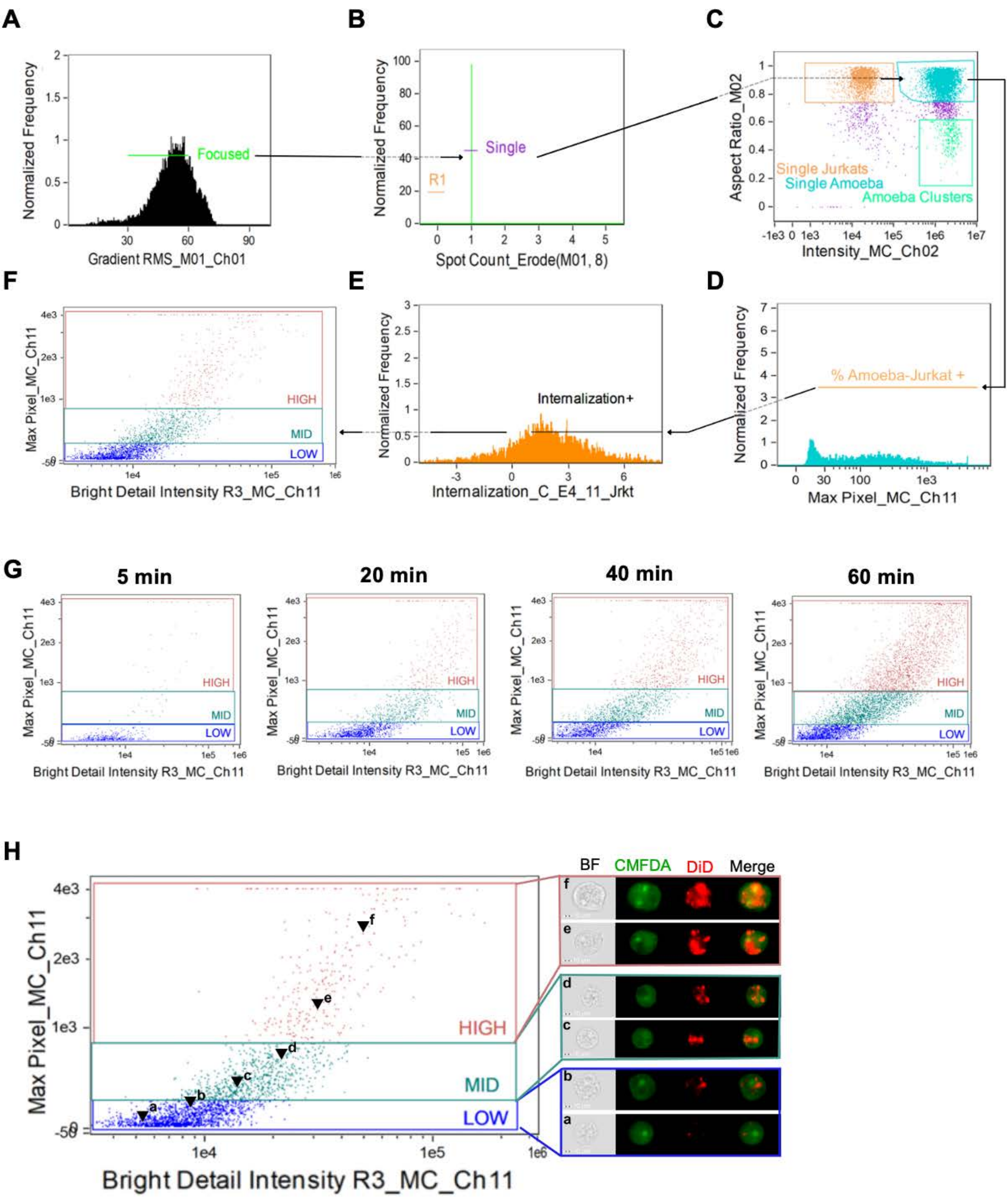

A
Normalized Frequency
Focused
Gradient RMS_M01_Ch01
B
Normalized Frequency
Single
R1
Spot Count_Erode(M01, 8)
C
Aspect Ratio_M02
Single Jurkats
Single Amoeba
Amoeba Clusters
Intensity_MC_Ch02
D
Normalized Frequency
% Amoeba-Jurkat +
Max Pixel_MC_Ch11
E
Normalized Frequency
Internalization+
Internalization_C_E4_11_Jrkt
F
Max Pixel_MC_Ch11
HIGH
MID
LOW
Bright Detail Intensity R3_MC_Ch11
G
5 min
20 min
40 min
60 min
H
Max Pixel_MC_Ch11
HIGH
MID
LOW
Bright Detail Intensity R3_MC_Ch11
BF
CMFDA
DiD
Merge

### Figure S2

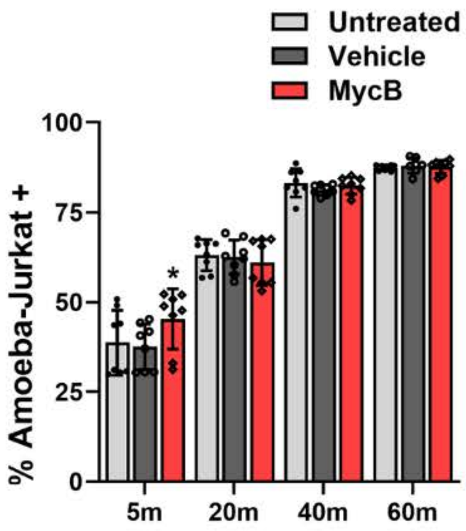

Untreated
Vehicle
MycB
% Amoeba-Jurkat +
100
75
50
25
0
*
5m
20m
40m
60m

**Figure S3**

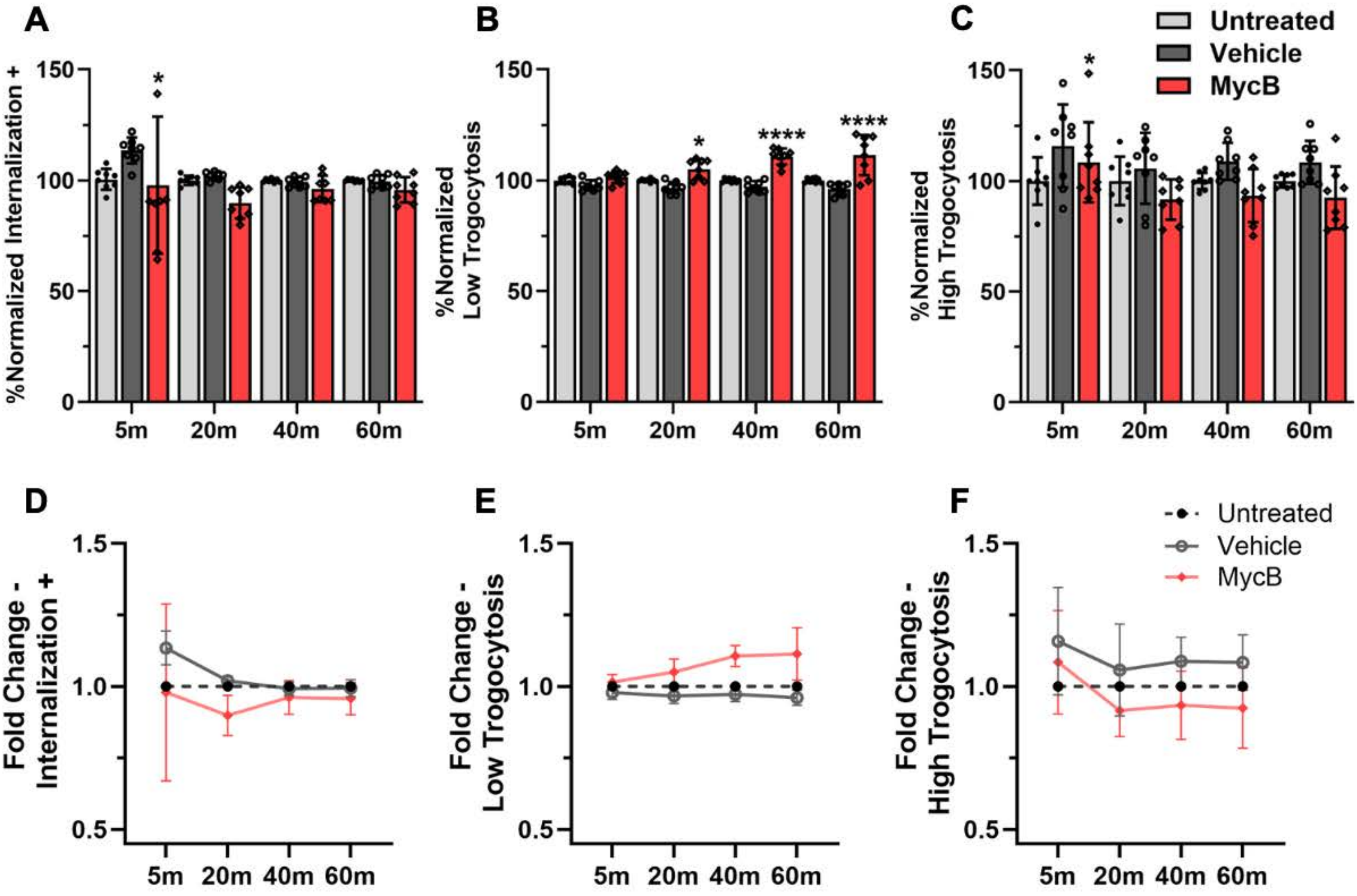

A
B
C
Untreated
Vehicle
MycB
%Normalized Internalization +
%Normalized Low Trogocytosis
%Normalized High Trogocytosis
150
100
50
0
5m
20m
40m
60m
*
****
D
E
F
Fold Change - Internalization +
Fold Change - Low Trogocytosis
Fold Change - High Trogocytosis
1.5
1.0
0.5

**Figure S4**

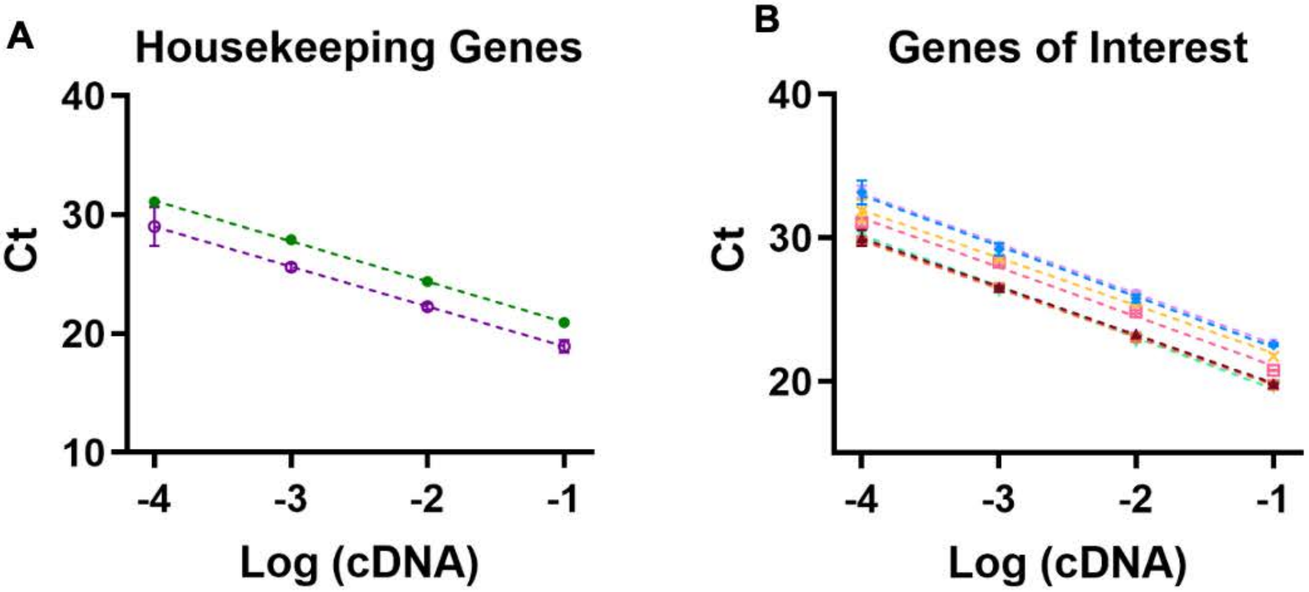


| PRIMER NAME | LINEAR EQUATION | qPCR EFFICIENCY |
|---|---|---|
| GAPDH | Y = -3.40*X + 17.59 | 96.86 % |
| RPLP0 | Y = -3.36*X + 15.58 | 98.54 % |

| PRIMER NAME | LINEAR EQUATION | qPCR EFFICIENCY |
|---|---|---|
| CFL1 | Y = -3.37*X + 16.49 | 97.93 % |
| MYPT1 | Y = -3.53*X + 18.85 | 92.05 % |
| Rac1 | Y = -3.35*X + 16.40 | 98.68 % |
| Rac2 | Y = -3.33*X + 18.64 | 99.81 % |
| RhoA | Y = -3.50*X + 19.10 | 93.07 % |
| ROCK1 | Y = -3.44*X + 17.64 | 95.28 % |
| ROCK2 | Y = -3.58*X + 15.89 | 90.18 % |

#### Figure S5

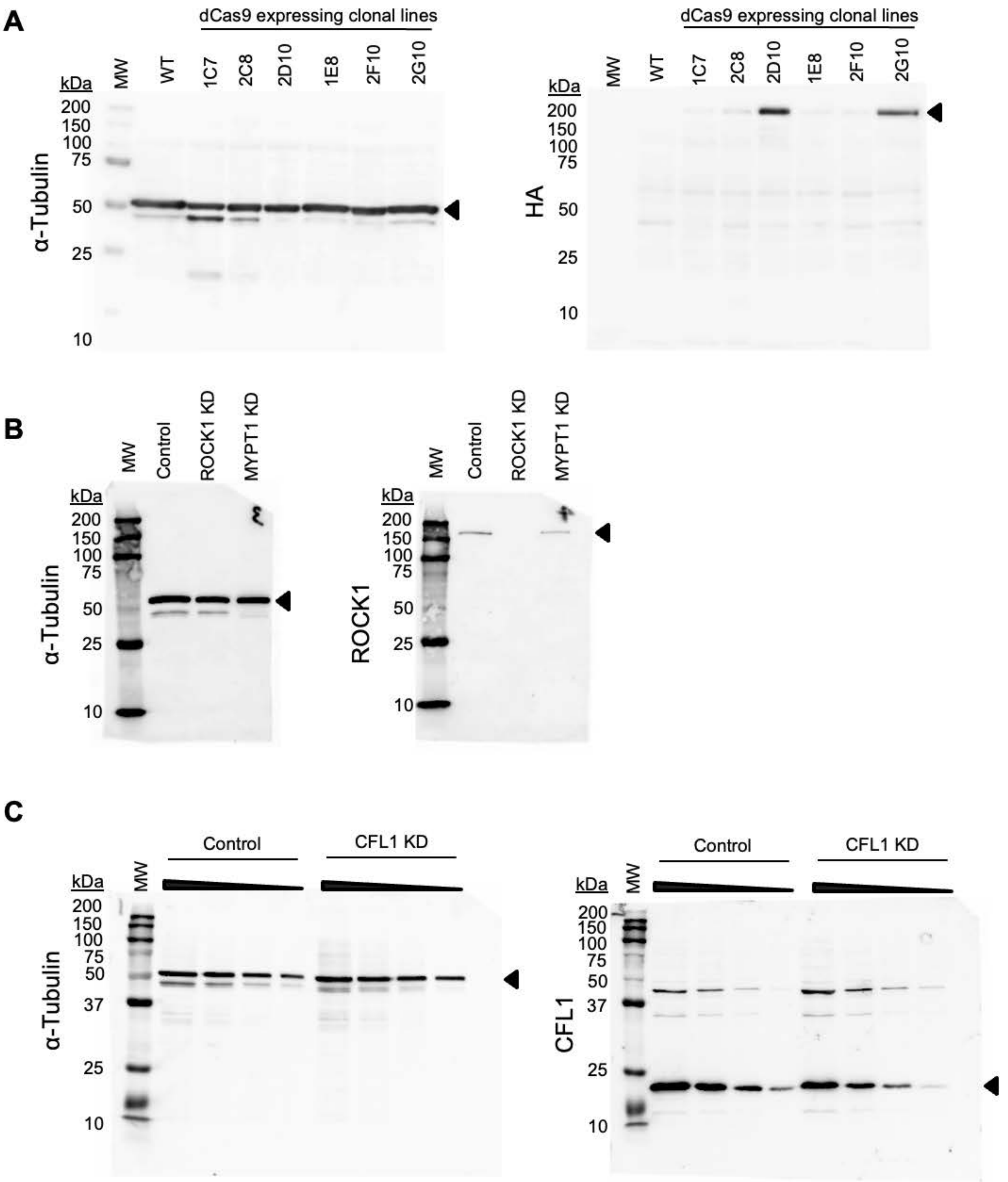
A
dCas9 expressing clonal lines
MW
WT
1C7
2C8
2D10
1E8
2F10
2G10
kDa
200
150
100
75
50
25
10
α-Tubulin
HA
B
Control
ROCK1 KD
MYPT1 KD
ROCK1
C
Control
CFL1 KD
37
CFL1

**Figure S6**

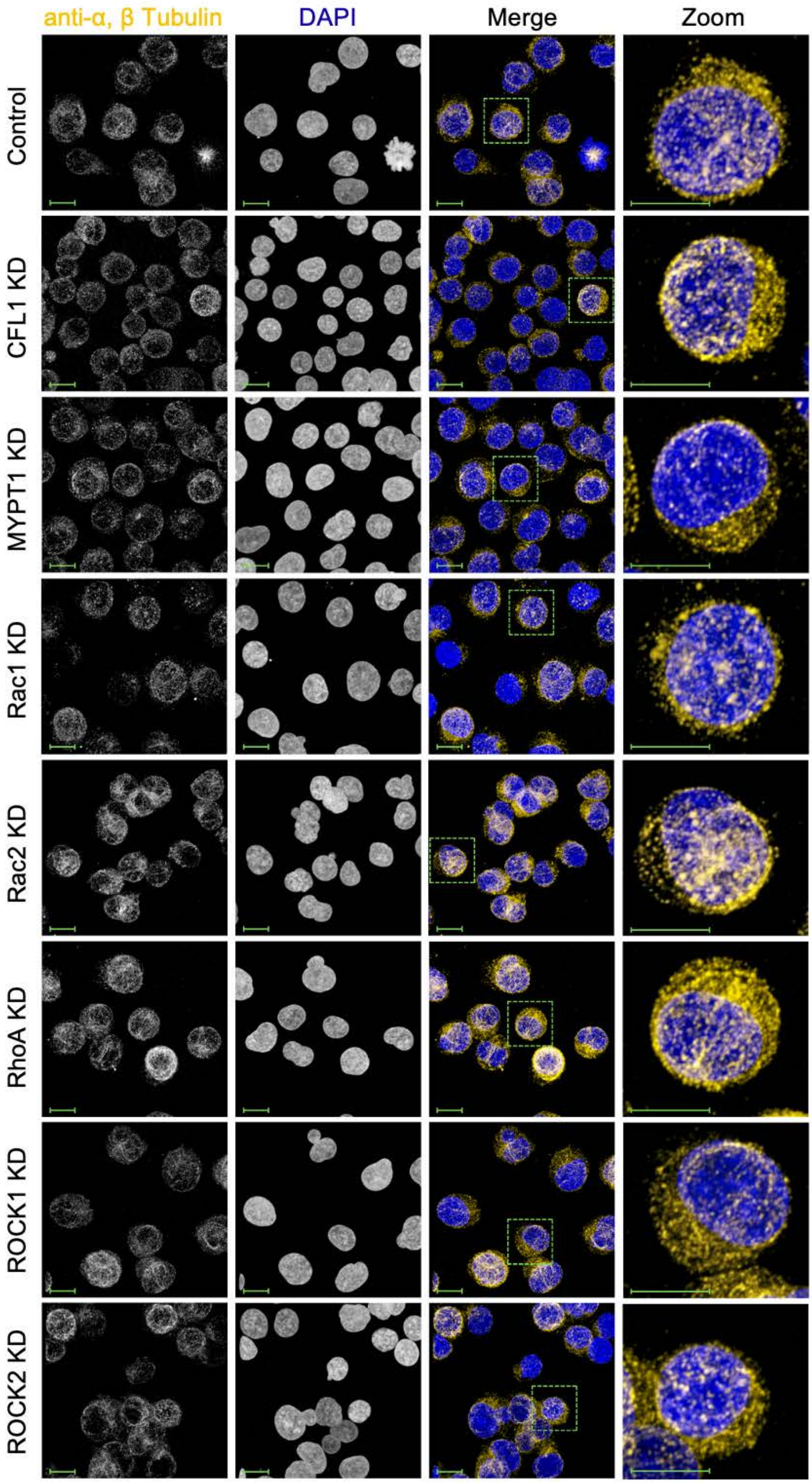

anti-α, β Tubulin
DAPI
Merge
Zoom
Control
CFL1 KD
MYPT1 KD
Rac1 KD
Rac2 KD
RhoA KD
ROCK1 KD
ROCK2 KD

#### Figure S7

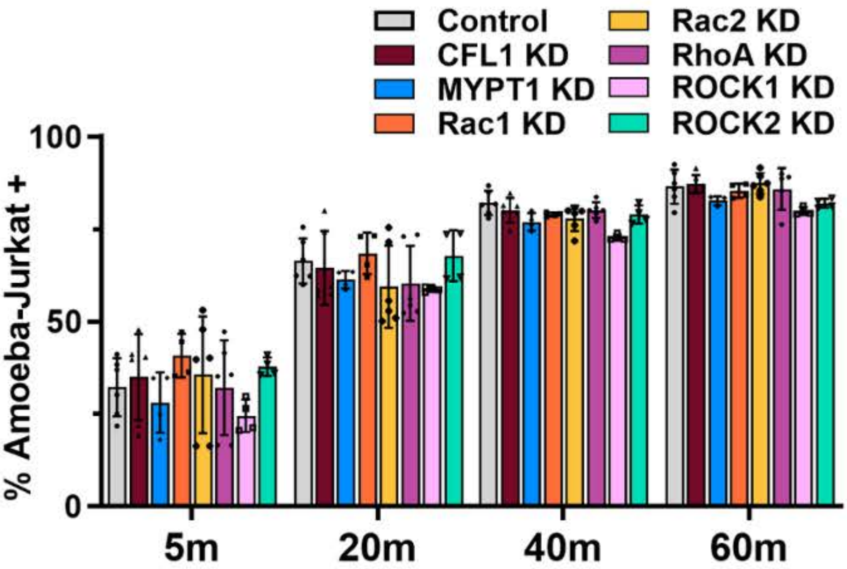

Control
CFL1 KD
MYPT1 KD
Rac1 KD
Rac2 KD
RhoA KD
ROCK1 KD
ROCK2 KD
% Amoeba-Jurkat +
100
50
0
5m
20m
40m
60m

**Figure S8**

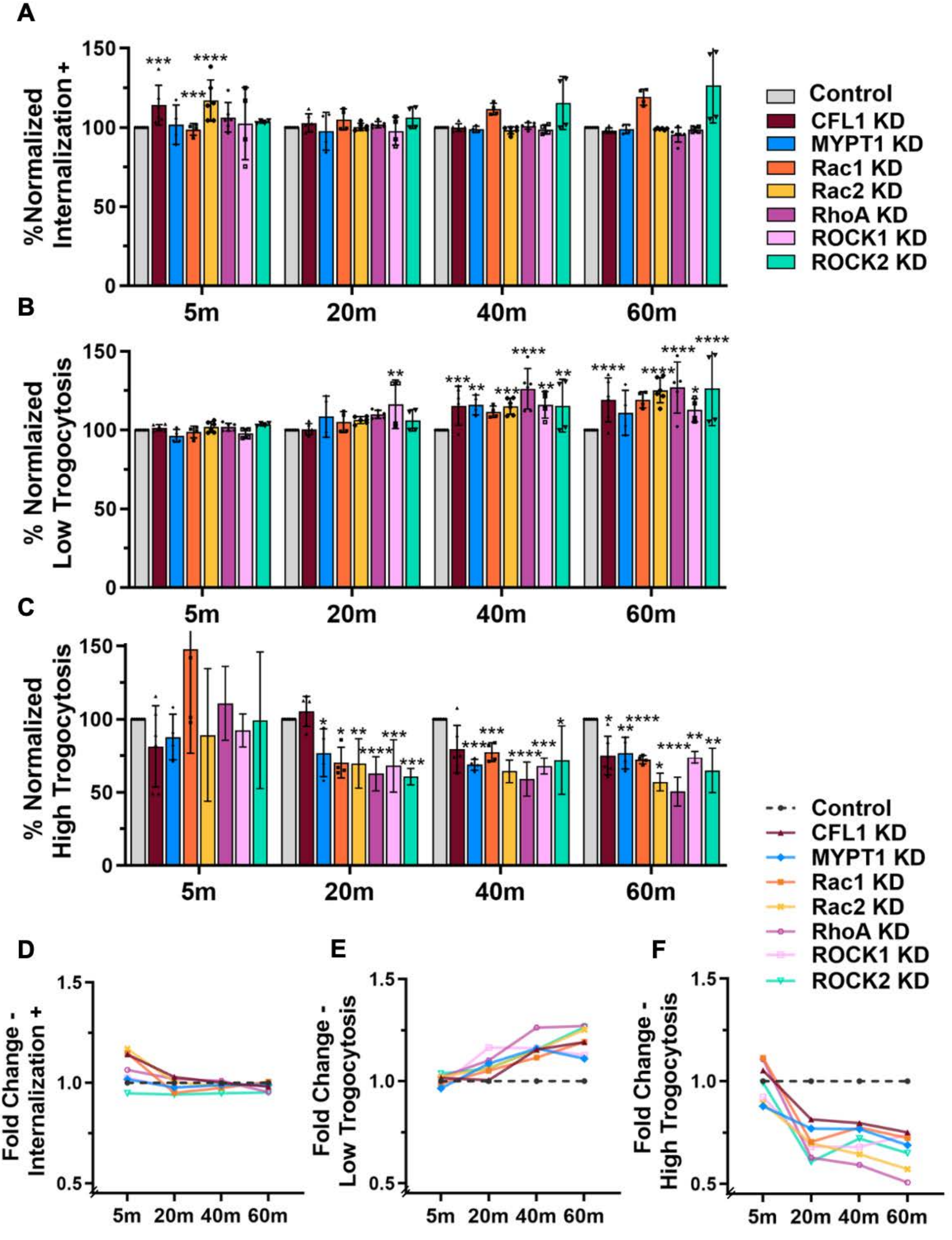
A
%Normalized Internalization +
Control
CFL1 KD
MYPT1 KD
Rac1 KD
Rac2 KD
RhoA KD
ROCK1 KD
ROCK2 KD
5m
20m
40m
60m
B
% Normalaized Low Trogocytosis
C
% Normalized High Trogocytosis
D
Fold Change - Internalization +
E
Fold Change - Low Trogocytosis
F
Fold Change - High Trogocytosis

#### Figure S9

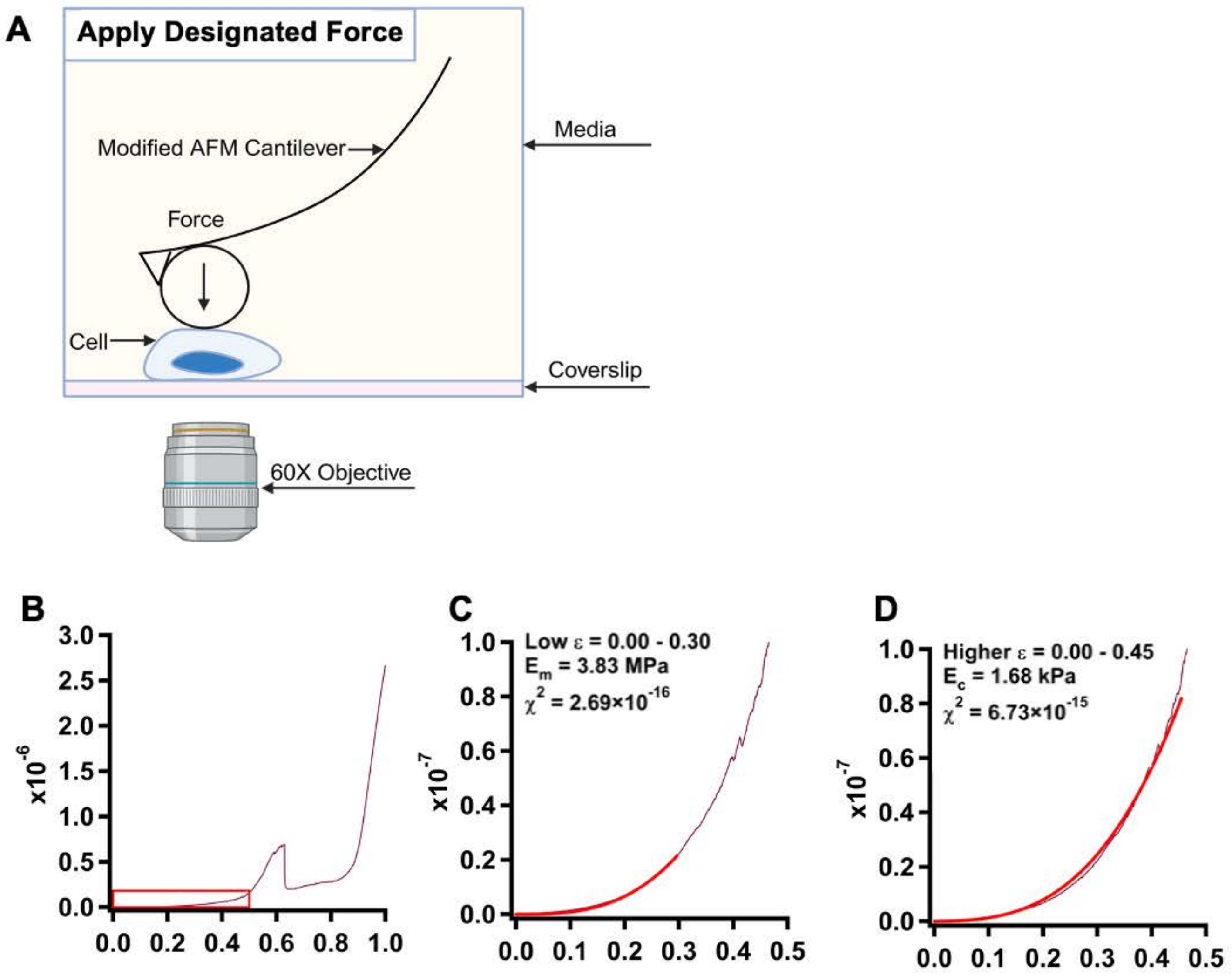
A
Apply Designated Force
Modified AFM Cantilever
Media
Force
Cell
Coverslip
60X Objective
B
x10$^{-6}$
3.0
2.5
2.0
1.5
1.0
0.5
0.0
0.0 0.2 0.4 0.6 0.8 1.0
C
Low ε = 0.00 - 0.30
$E_m$ = 3.83 MPa
$\chi^2$ = 2.69×10$^{-16}$
x10$^{-7}$
1.0
0.8
0.6
0.4
0.2
0.0
0.0 0.1 0.2 0.3 0.4 0.5
D
Higher ε = 0.00 - 0.45
$E_c$ = 1.68 kPa
$\chi^2$ = 6.73×10$^{-15}$
x10$^{-7}$
1.0
0.8
0.6
0.4
0.2
0.0
0.0 0.1 0.2 0.3 0.4 0.5

**Figure S10**

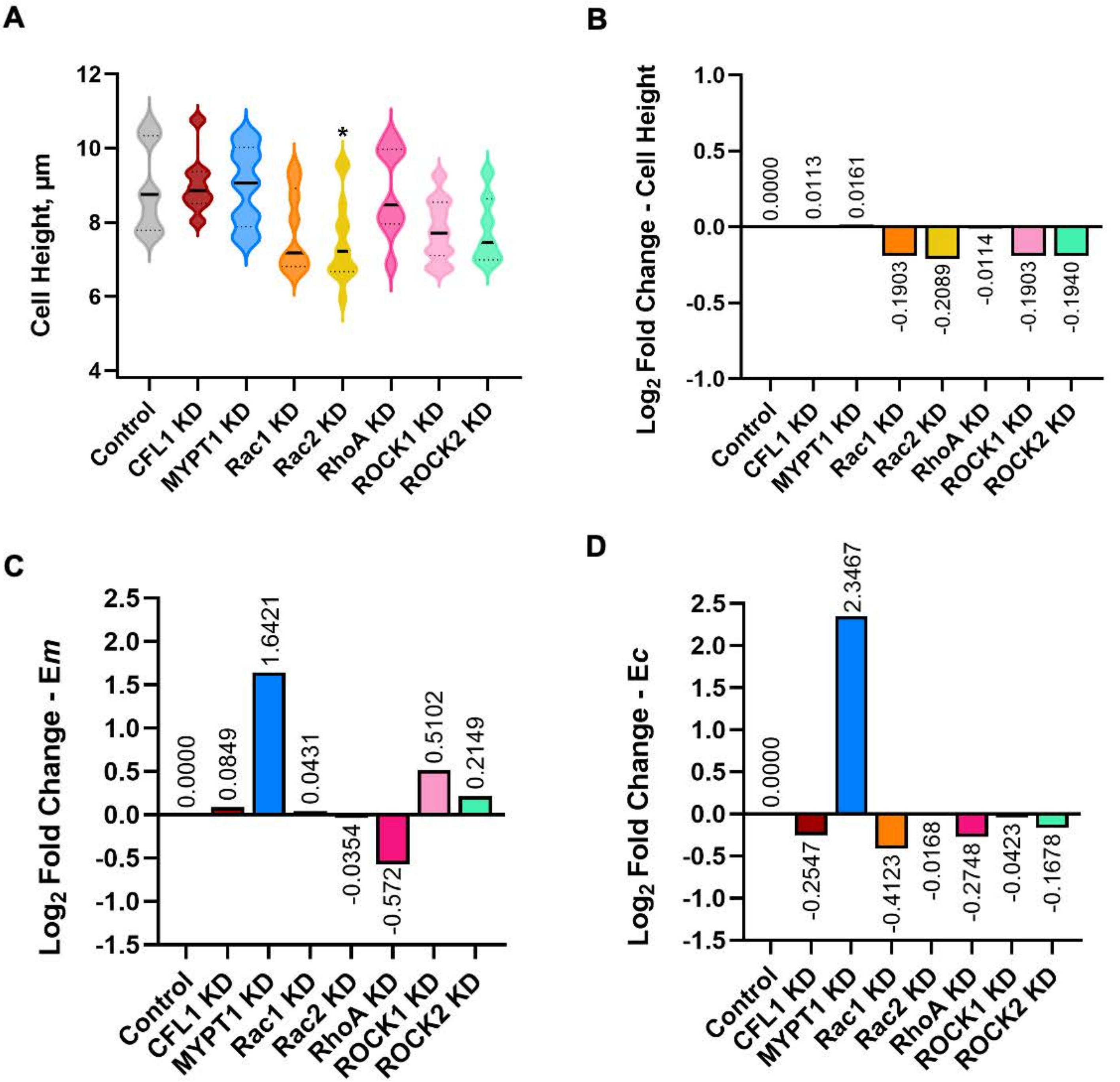
A
B
C
D
Cell Height, μm
12
10
8
6
4
*
Control
CFL1 KD
MYPT1 KD
Rac1 KD
Rac2 KD
RhoA KD
ROCK1 KD
ROCK2 KD
Log2 Fold Change - Cell Height
1.0
0.5
0.0
-0.5
-1.0
0.0000
0.0113
0.0161
-0.1903
-0.2089
-0.0114
-0.1903
-0.1940
Log2 Fold Change - Em
2.5
2.0
1.5
1.0
0.5
0.0
-0.5
-1.0
-1.5
0.0000
0.0849
1.6421
0.0431
-0.0354
-0.572
0.5102
0.2149
Log2 Fold Change - Ec
0.0000
-0.2547
2.3467
-0.4123
-0.0168
-0.2748
-0.0423
-0.1678

### Figure S11

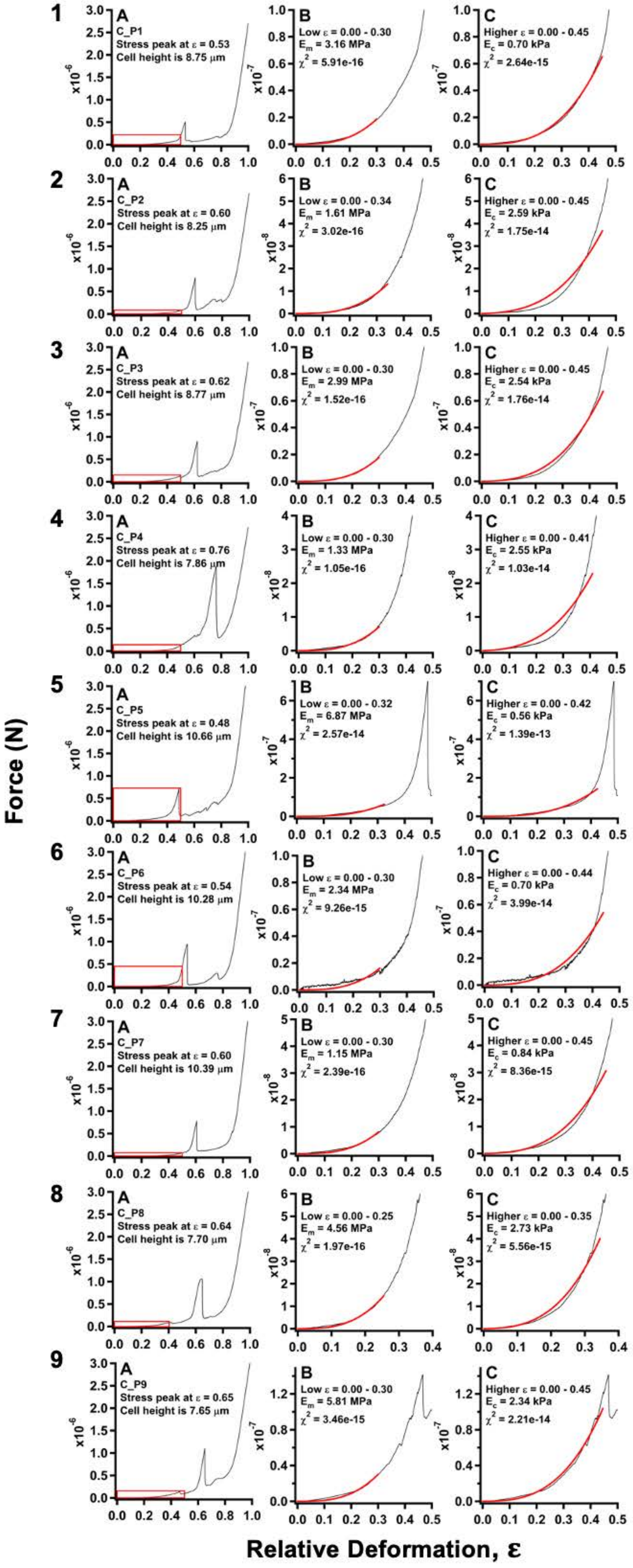

C_P1
Stress peak at ε = 0.53
Cell height is 8.75 μm
Low ε = 0.00 - 0.30
Em = 3.16 MPa
χ2 = 5.91e-16
Higher ε = 0.00 - 0.45
Ec = 0.70 kPa
χ2 = 2.64e-15
C_P2
Stress peak at ε = 0.60
Cell height is 8.25 μm
Low ε = 0.00 - 0.34
Em = 1.61 MPa
χ2 = 3.02e-16
Higher ε = 0.00 - 0.45
Ec = 2.59 kPa
χ2 = 1.75e-14
C_P3
Stress peak at ε = 0.62
Cell height is 8.77 μm
Low ε = 0.00 - 0.30
Em = 2.99 MPa
χ2 = 1.52e-16
Higher ε = 0.00 - 0.45
Ec = 2.54 kPa
χ2 = 1.76e-14
C_P4
Stress peak at ε = 0.76
Cell height is 7.86 μm
Low ε = 0.00 - 0.30
Em = 1.33 MPa
χ2 = 1.05e-16
Higher ε = 0.00 - 0.41
Ec = 2.55 kPa
χ2 = 1.03e-14
C_P5
Stress peak at ε = 0.48
Cell height is 10.66 μm
Low ε = 0.00 - 0.32
Em = 6.87 MPa
χ2 = 2.57e-14
Higher ε = 0.00 - 0.42
Ec = 0.56 kPa
χ2 = 1.39e-13
C_P6
Stress peak at ε = 0.54
Cell height is 10.28 μm
Low ε = 0.00 - 0.30
Em = 2.34 MPa
χ2 = 9.26e-15
Higher ε = 0.00 - 0.44
Ec = 0.70 kPa
χ2 = 3.99e-14
C_P7
Stress peak at ε = 0.60
Cell height is 10.39 μm
Low ε = 0.00 - 0.30
Em = 1.15 MPa
χ2 = 2.39e-16
Higher ε = 0.00 - 0.45
Ec = 0.84 kPa
χ2 = 8.36e-15
C_P8
Stress peak at ε = 0.64
Cell height is 7.70 μm
Low ε = 0.00 - 0.25
Em = 4.56 MPa
χ2 = 1.97e-16
Higher ε = 0.00 - 0.35
Ec = 2.73 kPa
χ2 = 5.56e-15
C_P9
Stress peak at ε = 0.65
Cell height is 7.65 μm
Low ε = 0.00 - 0.30
Em = 5.81 MPa
χ2 = 3.46e-15
Higher ε = 0.00 - 0.45
Ec = 2.34 kPa
χ2 = 2.21e-14
Force (N)
Relative Deformation, ε

#### Figure S12

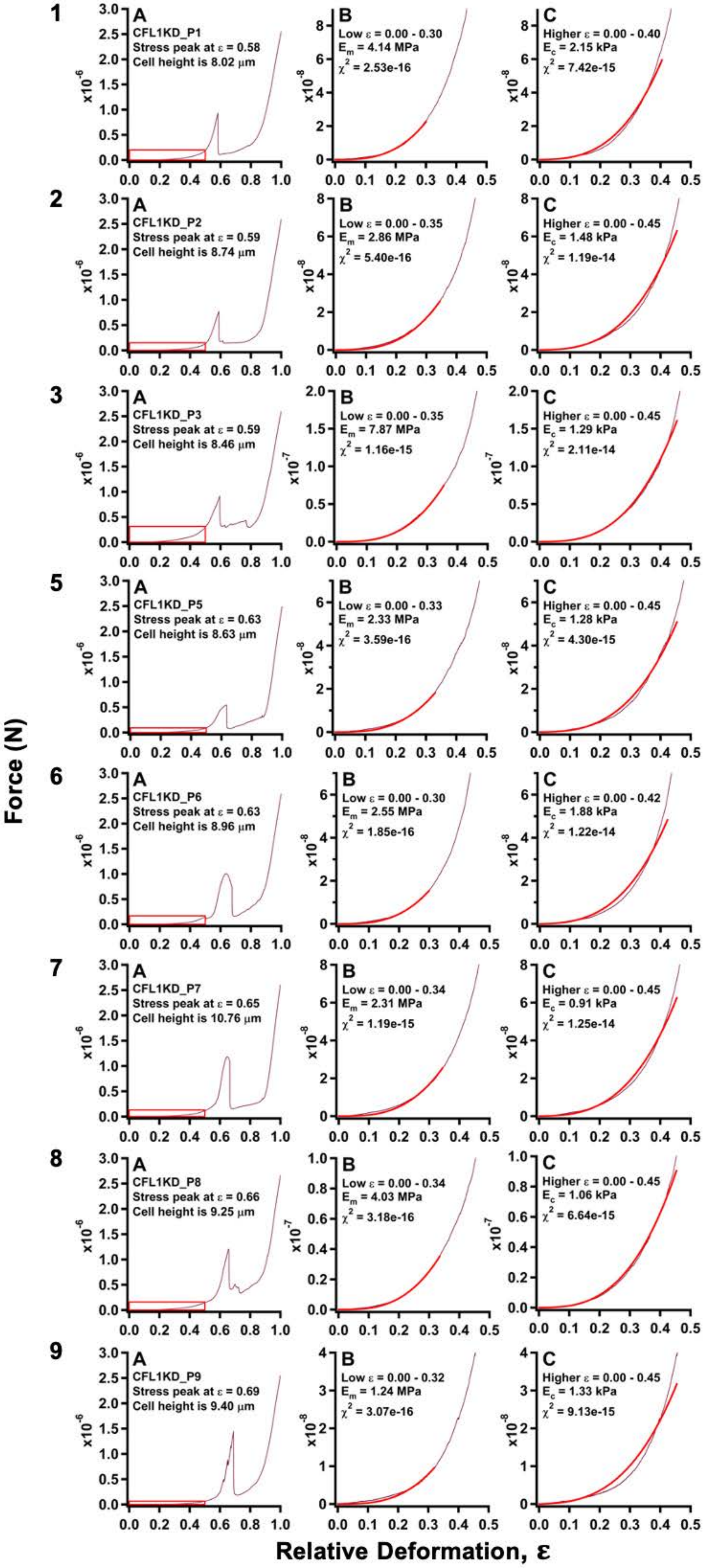
1
A
CFL1KD_P1
Stress peak at ε = 0.58
Cell height is 8.02 μm
B
Low ε = 0.00 - 0.30
Em = 4.14 MPa
χ2 = 2.53e-16
C
Higher ε = 0.00 - 0.40
Ec = 2.15 kPa
χ2 = 7.42e-15
2
A
CFL1KD_P2
Stress peak at ε = 0.59
Cell height is 8.74 μm
B
Low ε = 0.00 - 0.35
Em = 2.86 MPa
χ2 = 5.40e-16
C
Higher ε = 0.00 - 0.45
Ec = 1.48 kPa
χ2 = 1.19e-14
3
A
CFL1KD_P3
Stress peak at ε = 0.59
Cell height is 8.46 μm
B
Low ε = 0.00 - 0.35
Em = 7.87 MPa
χ2 = 1.16e-15
C
Higher ε = 0.00 - 0.45
Ec = 1.29 kPa
χ2 = 2.11e-14
5
A
CFL1KD_P5
Stress peak at ε = 0.63
Cell height is 8.63 μm
B
Low ε = 0.00 - 0.33
Em = 2.33 MPa
χ2 = 3.59e-16
C
Higher ε = 0.00 - 0.45
Ec = 1.28 kPa
χ2 = 4.30e-15
6
A
CFL1KD_P6
Stress peak at ε = 0.63
Cell height is 8.96 μm
B
Low ε = 0.00 - 0.30
Em = 2.55 MPa
χ2 = 1.85e-16
C
Higher ε = 0.00 - 0.42
Ec = 1.88 kPa
χ2 = 1.22e-14
7
A
CFL1KD_P7
Stress peak at ε = 0.65
Cell height is 10.76 μm
B
Low ε = 0.00 - 0.34
Em = 2.31 MPa
χ2 = 1.19e-15
C
Higher ε = 0.00 - 0.45
Ec = 0.91 kPa
χ2 = 1.25e-14
8
A
CFL1KD_P8
Stress peak at ε = 0.66
Cell height is 9.25 μm
B
Low ε = 0.00 - 0.34
Em = 4.03 MPa
χ2 = 3.18e-16
C
Higher ε = 0.00 - 0.45
Ec = 1.06 kPa
χ2 = 6.64e-15
9
A
CFL1KD_P9
Stress peak at ε = 0.69
Cell height is 9.40 μm
B
Low ε = 0.00 - 0.32
Em = 1.24 MPa
χ2 = 3.07e-16
C
Higher ε = 0.00 - 0.45
Ec = 1.33 kPa
χ2 = 9.13e-15
Force (N)
Relative Deformation, ε

#### Figure S13

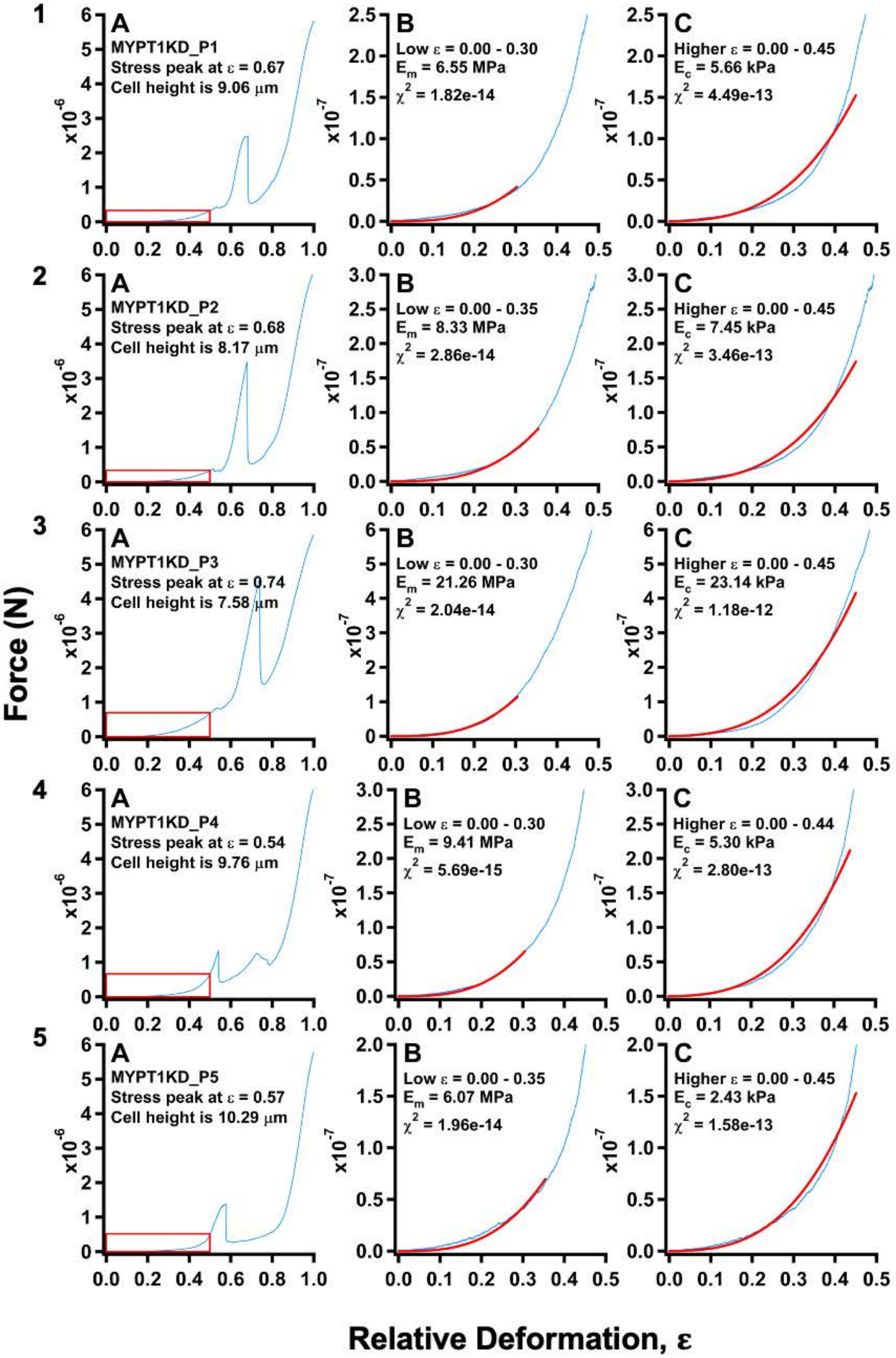

1
A
MYPT1KD_P1
Stress peak at ε = 0.67
Cell height is 9.06 μm
B
Low ε = 0.00 - 0.30
E_m = 6.55 MPa
χ^2 = 1.82e-14
C
Higher ε = 0.00 - 0.45
E_c = 5.66 kPa
χ^2 = 4.49e-13
2
A
MYPT1KD_P2
Stress peak at ε = 0.68
Cell height is 8.17 μm
B
Low ε = 0.00 - 0.35
E_m = 8.33 MPa
χ^2 = 2.86e-14
C
Higher ε = 0.00 - 0.45
E_c = 7.45 kPa
χ^2 = 3.46e-13
3
A
MYPT1KD_P3
Stress peak at ε = 0.74
Cell height is 7.58 μm
B
Low ε = 0.00 - 0.30
E_m = 21.26 MPa
χ^2 = 2.04e-14
C
Higher ε = 0.00 - 0.45
E_c = 23.14 kPa
χ^2 = 1.18e-12
4
A
MYPT1KD_P4
Stress peak at ε = 0.54
Cell height is 9.76 μm
B
Low ε = 0.00 - 0.30
E_m = 9.41 MPa
χ^2 = 5.69e-15
C
Higher ε = 0.00 - 0.44
E_c = 5.30 kPa
χ^2 = 2.80e-13
5
A
MYPT1KD_P5
Stress peak at ε = 0.57
Cell height is 10.29 μm
B
Low ε = 0.00 - 0.35
E_m = 6.07 MPa
χ^2 = 1.96e-14
C
Higher ε = 0.00 - 0.45
E_c = 2.43 kPa
χ^2 = 1.58e-13
x10^-6
x10^-7
Force (N)
Relative Deformation, ε

#### Figure S14

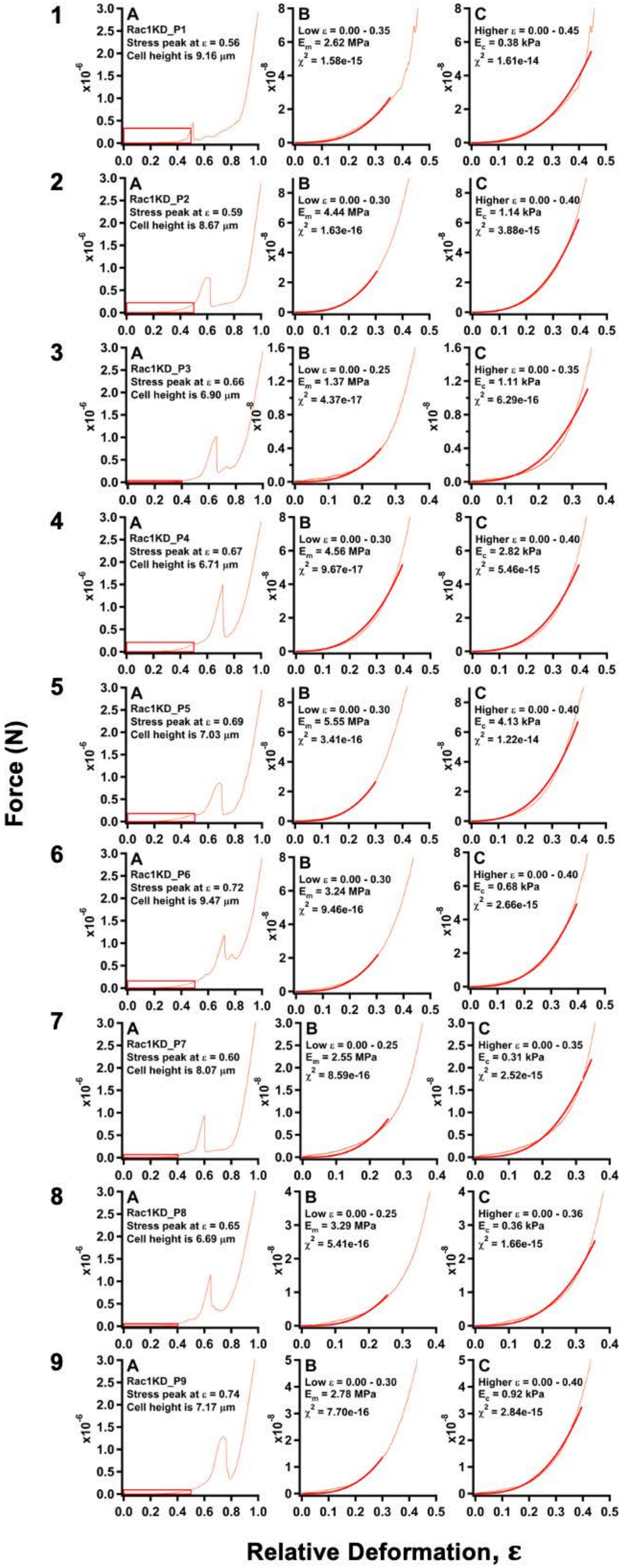

Rac1KD_P1
Stress peak at ε = 0.56
Cell height is 9.16 μm
Low ε = 0.00 - 0.35
E_m = 2.62 MPa
χ^2 = 1.58e-15
Higher ε = 0.00 - 0.45
E_c = 0.38 kPa
χ^2 = 1.61e-14
Rac1KD_P2
Stress peak at ε = 0.59
Cell height is 8.67 μm
Low ε = 0.00 - 0.30
E_m = 4.44 MPa
χ^2 = 1.63e-16
Higher ε = 0.00 - 0.40
E_c = 1.14 kPa
χ^2 = 3.88e-15
Rac1KD_P3
Stress peak at ε = 0.66
Cell height is 6.90 μm
Low ε = 0.00 - 0.25
E_m = 1.37 MPa
χ^2 = 4.37e-17
Higher ε = 0.00 - 0.35
E_c = 1.11 kPa
χ^2 = 6.29e-16
Rac1KD_P4
Stress peak at ε = 0.67
Cell height is 6.71 μm
Low ε = 0.00 - 0.30
E_m = 4.56 MPa
χ^2 = 9.67e-17
Higher ε = 0.00 - 0.40
E_c = 2.82 kPa
χ^2 = 5.46e-15
Rac1KD_P5
Stress peak at ε = 0.69
Cell height is 7.03 μm
Low ε = 0.00 - 0.30
E_m = 5.55 MPa
χ^2 = 3.41e-16
Higher ε = 0.00 - 0.40
E_c = 4.13 kPa
χ^2 = 1.22e-14
Rac1KD_P6
Stress peak at ε = 0.72
Cell height is 9.47 μm
Low ε = 0.00 - 0.30
E_m = 3.24 MPa
χ^2 = 9.46e-16
Higher ε = 0.00 - 0.40
E_c = 0.68 kPa
χ^2 = 2.66e-15
Rac1KD_P7
Stress peak at ε = 0.60
Cell height is 8.07 μm
Low ε = 0.00 - 0.25
E_m = 2.55 MPa
χ^2 = 8.59e-16
Higher ε = 0.00 - 0.35
E_c = 0.31 kPa
χ^2 = 2.52e-15
Rac1KD_P8
Stress peak at ε = 0.65
Cell height is 6.69 μm
Low ε = 0.00 - 0.25
E_m = 3.29 MPa
χ^2 = 5.41e-16
Higher ε = 0.00 - 0.36
E_c = 0.36 kPa
χ^2 = 1.66e-15
Rac1KD_P9
Stress peak at ε = 0.74
Cell height is 7.17 μm
Low ε = 0.00 - 0.30
E_m = 2.78 MPa
χ^2 = 7.70e-16
Higher ε = 0.00 - 0.40
E_c = 0.92 kPa
χ^2 = 2.84e-15
Force (N)
Relative Deformation, ε

Figure S15

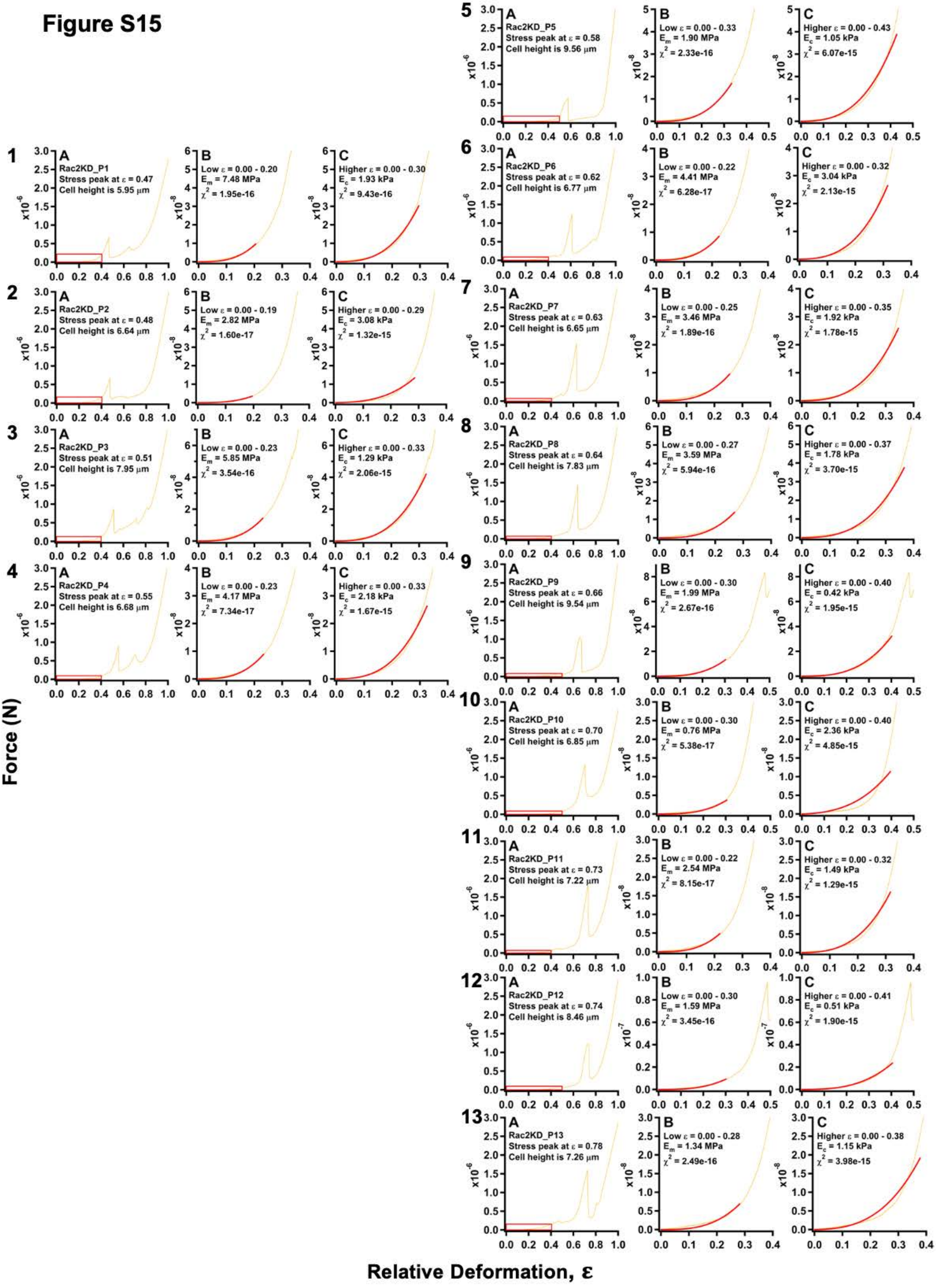

Rac2KD_P1
Stress peak at ε = 0.47
Cell height is 5.95 μm
Low ε = 0.00 - 0.20
E_m = 7.48 MPa
χ² = 1.95e-16
Higher ε = 0.00 - 0.30
E_c = 1.93 kPa
χ² = 9.43e-16
Rac2KD_P2
Stress peak at ε = 0.48
Cell height is 6.64 μm
Low ε = 0.00 - 0.19
E_m = 2.82 MPa
χ² = 1.60e-17
Higher ε = 0.00 - 0.29
E_c = 3.08 kPa
χ² = 1.32e-15
Rac2KD_P3
Stress peak at ε = 0.51
Cell height is 7.95 μm
Low ε = 0.00 - 0.23
E_m = 5.85 MPa
χ² = 3.54e-16
Higher ε = 0.00 - 0.33
E_c = 1.29 kPa
χ² = 2.06e-15
Rac2KD_P4
Stress peak at ε = 0.55
Cell height is 6.68 μm
Low ε = 0.00 - 0.23
E_m = 4.17 MPa
χ² = 7.34e-17
Higher ε = 0.00 - 0.33
E_c = 2.18 kPa
χ² = 1.67e-15
Rac2KD_P5
Stress peak at ε = 0.58
Cell height is 9.56 μm
Low ε = 0.00 - 0.33
E_m = 1.90 MPa
χ² = 2.33e-16
Higher ε = 0.00 - 0.43
E_c = 1.05 kPa
χ² = 6.07e-15
Rac2KD_P6
Stress peak at ε = 0.62
Cell height is 6.77 μm
Low ε = 0.00 - 0.22
E_m = 4.41 MPa
χ² = 6.28e-17
Higher ε = 0.00 - 0.32
E_c = 3.04 kPa
χ² = 2.13e-15
Rac2KD_P7
Stress peak at ε = 0.63
Cell height is 6.65 μm
Low ε = 0.00 - 0.25
E_m = 3.46 MPa
χ² = 1.89e-16
Higher ε = 0.00 - 0.35
E_c = 1.92 kPa
χ² = 1.78e-15
Rac2KD_P8
Stress peak at ε = 0.64
Cell height is 7.83 μm
Low ε = 0.00 - 0.27
E_m = 3.59 MPa
χ² = 5.94e-16
Higher ε = 0.00 - 0.37
E_c = 1.78 kPa
χ² = 3.70e-15
Rac2KD_P9
Stress peak at ε = 0.66
Cell height is 9.54 μm
Low ε = 0.00 - 0.30
E_m = 1.99 MPa
χ² = 2.67e-16
Higher ε = 0.00 - 0.40
E_c = 0.42 kPa
χ² = 1.95e-15
Rac2KD_P10
Stress peak at ε = 0.70
Cell height is 6.85 μm
Low ε = 0.00 - 0.30
E_m = 0.76 MPa
χ² = 5.38e-17
Higher ε = 0.00 - 0.40
E_c = 2.36 kPa
χ² = 4.85e-15
Rac2KD_P11
Stress peak at ε = 0.73
Cell height is 7.22 μm
Low ε = 0.00 - 0.22
E_m = 2.54 MPa
χ² = 8.15e-17
Higher ε = 0.00 - 0.32
E_c = 1.49 kPa
χ² = 1.29e-15
Rac2KD_P12
Stress peak at ε = 0.74
Cell height is 8.46 μm
Low ε = 0.00 - 0.30
E_m = 1.59 MPa
χ² = 3.45e-16
Higher ε = 0.00 - 0.41
E_c = 0.51 kPa
χ² = 1.90e-15
Rac2KD_P13
Stress peak at ε = 0.78
Cell height is 7.26 μm
Low ε = 0.00 - 0.28
E_m = 1.34 MPa
χ² = 2.49e-16
Higher ε = 0.00 - 0.38
E_c = 1.15 kPa
χ² = 3.98e-15
Force (N)
Relative Deformation, ε

**Figure S16**

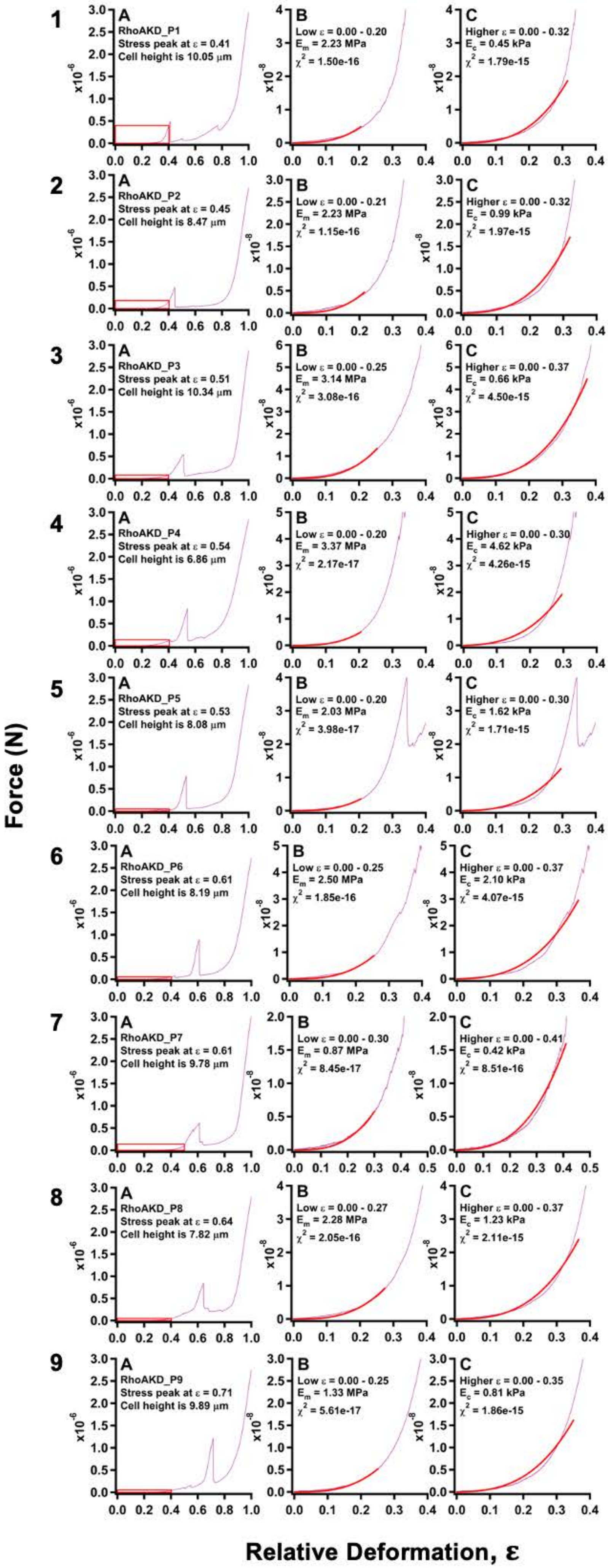
RhoAKD_P1
Stress peak at ε = 0.41
Cell height is 10.05 μm
Low ε = 0.00 - 0.20
E_m = 2.23 MPa
χ^2 = 1.50e-16
Higher ε = 0.00 - 0.32
E_c = 0.45 kPa
χ^2 = 1.79e-15
RhoAKD_P2
Stress peak at ε = 0.45
Cell height is 8.47 μm
Low ε = 0.00 - 0.21
E_m = 2.23 MPa
χ^2 = 1.15e-16
Higher ε = 0.00 - 0.32
E_c = 0.99 kPa
χ^2 = 1.97e-15
RhoAKD_P3
Stress peak at ε = 0.51
Cell height is 10.34 μm
Low ε = 0.00 - 0.25
E_m = 3.14 MPa
χ^2 = 3.08e-16
Higher ε = 0.00 - 0.37
E_c = 0.66 kPa
χ^2 = 4.50e-15
RhoAKD_P4
Stress peak at ε = 0.54
Cell height is 6.86 μm
Low ε = 0.00 - 0.20
E_m = 3.37 MPa
χ^2 = 2.17e-17
Higher ε = 0.00 - 0.30
E_c = 4.62 kPa
χ^2 = 4.26e-15
RhoAKD_P5
Stress peak at ε = 0.53
Cell height is 8.08 μm
Low ε = 0.00 - 0.20
E_m = 2.03 MPa
χ^2 = 3.98e-17
Higher ε = 0.00 - 0.30
E_c = 1.62 kPa
χ^2 = 1.71e-15
RhoAKD_P6
Stress peak at ε = 0.61
Cell height is 8.19 μm
Low ε = 0.00 - 0.25
E_m = 2.50 MPa
χ^2 = 1.85e-16
Higher ε = 0.00 - 0.37
E_c = 2.10 kPa
χ^2 = 4.07e-15
RhoAKD_P7
Stress peak at ε = 0.61
Cell height is 9.78 μm
Low ε = 0.00 - 0.30
E_m = 0.87 MPa
χ^2 = 8.45e-17
Higher ε = 0.00 - 0.41
E_c = 0.42 kPa
χ^2 = 8.51e-16
RhoAKD_P8
Stress peak at ε = 0.64
Cell height is 7.82 μm
Low ε = 0.00 - 0.27
E_m = 2.28 MPa
χ^2 = 2.05e-16
Higher ε = 0.00 - 0.37
E_c = 1.23 kPa
χ^2 = 2.11e-15
RhoAKD_P9
Stress peak at ε = 0.71
Cell height is 9.89 μm
Low ε = 0.00 - 0.25
E_m = 1.33 MPa
χ^2 = 5.61e-17
Higher ε = 0.00 - 0.35
E_c = 0.81 kPa
χ^2 = 1.86e-15
Force (N)
Relative Deformation, ε

**Figure S17**

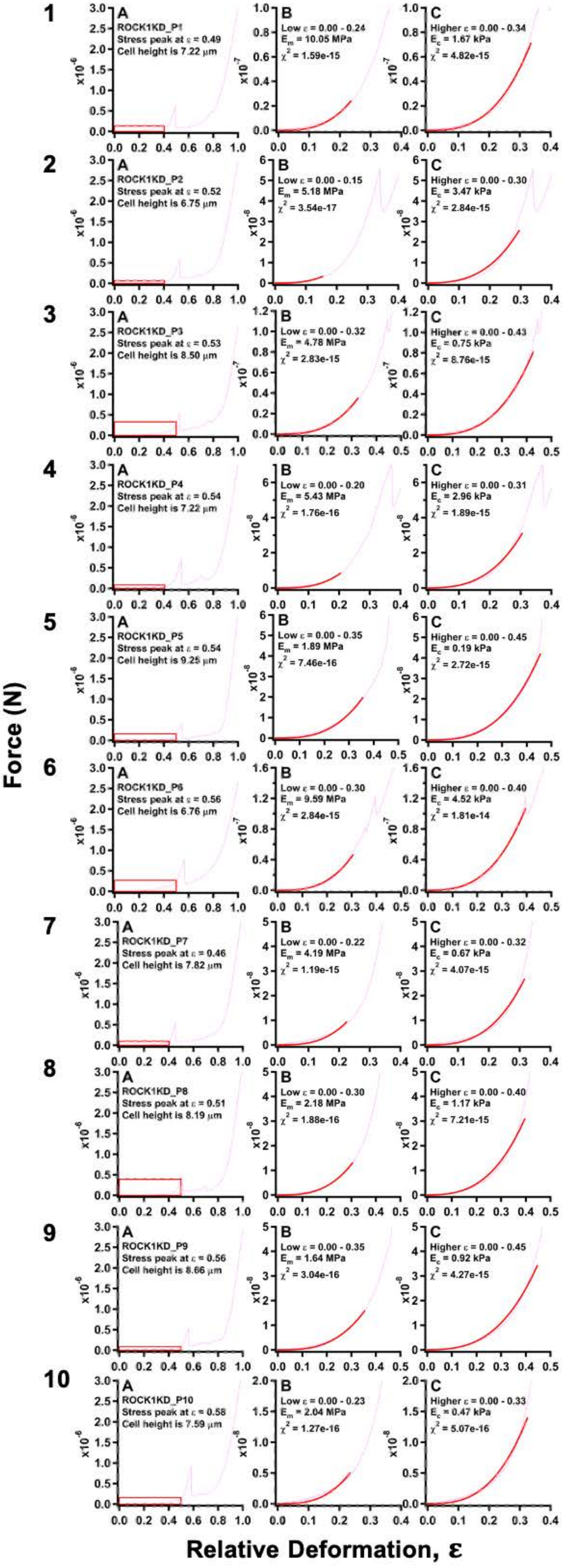

Force (N)
Relative Deformation, ε
ROCK1KD_P1
Stress peak at ε = 0.49
Cell height is 7.22 μm
Low ε = 0.00 - 0.24
E_m = 10.05 MPa
χ² = 1.59e-15
Higher ε = 0.00 - 0.34
E_c = 1.67 kPa
χ² = 4.82e-15
ROCK1KD_P2
Stress peak at ε = 0.52
Cell height is 6.75 μm
Low ε = 0.00 - 0.15
E_m = 5.18 MPa
χ² = 3.54e-17
Higher ε = 0.00 - 0.30
E_c = 3.47 kPa
χ² = 2.84e-15
ROCK1KD_P3
Stress peak at ε = 0.53
Cell height is 8.50 μm
Low ε = 0.00 - 0.32
E_m = 4.78 MPa
χ² = 2.83e-15
Higher ε = 0.00 - 0.43
E_c = 0.75 kPa
χ² = 8.76e-15
ROCK1KD_P4
Stress peak at ε = 0.54
Cell height is 7.22 μm
Low ε = 0.00 - 0.20
E_m = 5.43 MPa
χ² = 1.76e-16
Higher ε = 0.00 - 0.31
E_c = 2.96 kPa
χ² = 1.89e-15
ROCK1KD_P5
Stress peak at ε = 0.54
Cell height is 9.25 μm
Low ε = 0.00 - 0.35
E_m = 1.89 MPa
χ² = 7.46e-16
Higher ε = 0.00 - 0.45
E_c = 0.19 kPa
χ² = 2.72e-15
ROCK1KD_P6
Stress peak at ε = 0.56
Cell height is 6.76 μm
Low ε = 0.00 - 0.30
E_m = 9.59 MPa
χ² = 2.84e-15
Higher ε = 0.00 - 0.40
E_c = 4.52 kPa
χ² = 1.81e-14
ROCK1KD_P7
Stress peak at ε = 0.46
Cell height is 7.82 μm
Low ε = 0.00 - 0.22
E_m = 4.19 MPa
χ² = 1.19e-15
Higher ε = 0.00 - 0.32
E_c = 0.67 kPa
χ² = 4.07e-15
ROCK1KD_P8
Stress peak at ε = 0.51
Cell height is 8.19 μm
Low ε = 0.00 - 0.30
E_m = 2.18 MPa
χ² = 1.88e-16
Higher ε = 0.00 - 0.40
E_c = 1.17 kPa
χ² = 7.21e-15
ROCK1KD_P9
Stress peak at ε = 0.56
Cell height is 8.66 μm
Low ε = 0.00 - 0.35
E_m = 1.64 MPa
χ² = 3.04e-16
Higher ε = 0.00 - 0.45
E_c = 0.92 kPa
χ² = 4.27e-15
ROCK1KD_P10
Stress peak at ε = 0.58
Cell height is 7.59 μm
Low ε = 0.00 - 0.23
E_m = 2.04 MPa
χ² = 1.27e-16
Higher ε = 0.00 - 0.33
E_c = 0.47 kPa
χ² = 5.07e-16

**Figure S18**

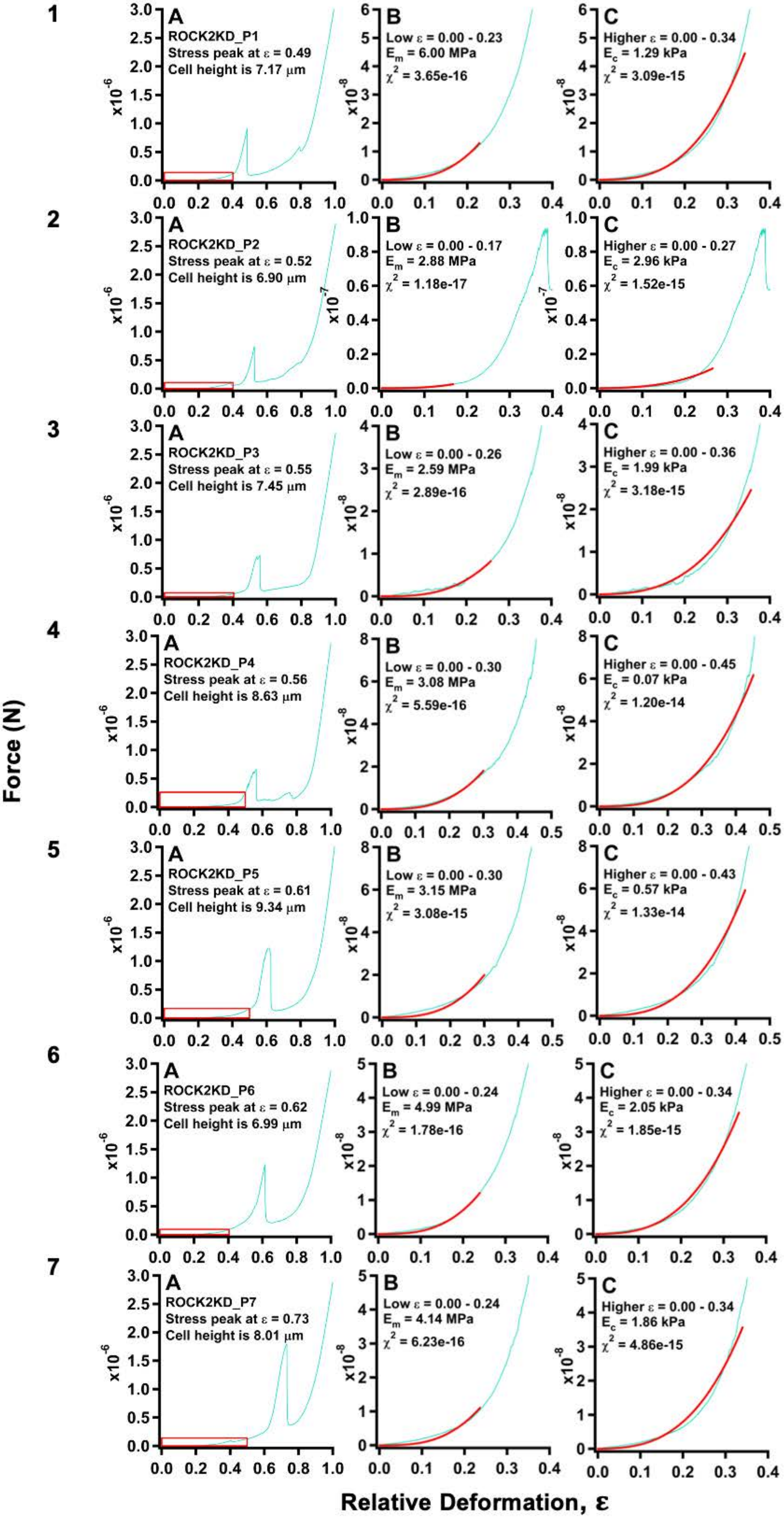

1
A
ROCK2KD_P1
Stress peak at ε = 0.49
Cell height is 7.17 μm
B
Low ε = 0.00 - 0.23
Em = 6.00 MPa
χ2 = 3.65e-16
C
Higher ε = 0.00 - 0.34
Ec = 1.29 kPa
χ2 = 3.09e-15
2
A
ROCK2KD_P2
Stress peak at ε = 0.52
Cell height is 6.90 μm
B
Low ε = 0.00 - 0.17
Em = 2.88 MPa
χ2 = 1.18e-17
C
Higher ε = 0.00 - 0.27
Ec = 2.96 kPa
χ2 = 1.52e-15
3
A
ROCK2KD_P3
Stress peak at ε = 0.55
Cell height is 7.45 μm
B
Low ε = 0.00 - 0.26
Em = 2.59 MPa
χ2 = 2.89e-16
C
Higher ε = 0.00 - 0.36
Ec = 1.99 kPa
χ2 = 3.18e-15
4
A
ROCK2KD_P4
Stress peak at ε = 0.56
Cell height is 8.63 μm
B
Low ε = 0.00 - 0.30
Em = 3.08 MPa
χ2 = 5.59e-16
C
Higher ε = 0.00 - 0.45
Ec = 0.07 kPa
χ2 = 1.20e-14
5
A
ROCK2KD_P5
Stress peak at ε = 0.61
Cell height is 9.34 μm
B
Low ε = 0.00 - 0.30
Em = 3.15 MPa
χ2 = 3.08e-15
C
Higher ε = 0.00 - 0.43
Ec = 0.57 kPa
χ2 = 1.33e-14
6
A
ROCK2KD_P6
Stress peak at ε = 0.62
Cell height is 6.99 μm
B
Low ε = 0.00 - 0.24
Em = 4.99 MPa
χ2 = 1.78e-16
C
Higher ε = 0.00 - 0.34
Ec = 2.05 kPa
χ2 = 1.85e-15
7
A
ROCK2KD_P7
Stress peak at ε = 0.73
Cell height is 8.01 μm
B
Low ε = 0.00 - 0.24
Em = 4.14 MPa
χ2 = 6.23e-16
C
Higher ε = 0.00 - 0.34
Ec = 1.86 kPa
χ2 = 4.86e-15
Force (N)
Relative Deformation, ε
